# Genomic and dietary transitions during the Mesolithic and Early Neolithic in Sicily

**DOI:** 10.1101/2020.03.11.986158

**Authors:** Marieke S. van de Loosdrecht, Marcello A. Mannino, Sahra Talamo, Vanessa Villalba-Mouco, Cosimo Posth, Franziska Aron, Guido Brandt, Marta Burri, Cäcilia Freund, Rita Radzeviciute, Raphaela Stahl, Antje Wissgott, Lysann Klausnitzer, Sarah Nagel, Matthias Meyer, Antonio Tagliacozzo, Marcello Piperno, Sebastiano Tusa, Carmine Collina, Vittoria Schimmenti, Rosaria Di Salvo, Kay Prüfer, Jean-Jacques Hublin, Stephan Schiffels, Choongwon Jeong, Wolfgang Haak, Johannes Krause

## Abstract

Southern Italy is a key region for understanding the agricultural transition in the Mediterranean due to its central position. We present a genomic transect for 19 prehistoric Sicilians that covers the Early Mesolithic to Early Neolithic period. We find that the Early Mesolithic hunter-gatherers (HGs) are a highly drifted sister lineage to Early Holocene western European HGs, whereas a quarter of the Late Mesolithic HGs ancestry is related to HGs from eastern Europe and the Near East. This indicates substantial gene flow from (south-)eastern Europe between the Early and Late Mesolithic. The Early Neolithic farmers are genetically most similar to those from the Balkan and Greece, and carry only a maximum of ∼7% ancestry from Sicilian Mesolithic HGs. Ancestry changes match changes in dietary profile and material culture, except for two individuals who may provide tentative initial evidence that HGs adopted elements of farming in Sicily.

**One-sentence summary:** Genome-wide and isotopic data from prehistoric Sicilians reveal a pre-farming connection to (south-) eastern Europe, and tentative initial evidence that hunter-gatherers adopted some Neolithic aspects prior to near-total replacement by early farmers.

## Introduction

Southern Italy and Sicily feature some of the earliest evidence for agricultural food production in the Central Mediterranean, starting as early as ∼6,000 calBCE (*1*) or earlier (∼6,200 calBCE (*2*)). In the Mediterranean area, two Early Neolithic meta-horizons had developed in parallel by ∼5,500 calBCE (*3–5*). In the eastern and central Mediterranean, Early Neolithic farmers produced Impressa Wares with various decorative impressed designs made with a wide selection of tools. In contrast, in the western Mediterranean the decorative designs were preferentially made with *Cardium* seashell impressions, resulting in the typical Cardial Ware pottery (*6*). In Sicily and southern Italy two Impressa Ware horizons appeared rapidly in a timeframe of ∼500 years. The very first aspect of Impressa Wares appeared 6,000-5,700 calBCE followed by the Impressed Ware of the Stentinello group (Stentinello/Kronio) around 5,800-5,500 calBCE (*1, 7, 8*). The Early Neolithic horizons in Sicily may have their origin in the early farming traditions in the Balkans (*9–11*).

Grotta dell’Uzzo, in northwestern Sicily, is a key site for understanding human prehistory in the Central Mediterranean, and has provided unique insights into the cultural, subsistence and dietary changes that took place in the transition from hunting and gathering to agro-pastoralism (*12–14*). The cave stratigraphy covers the late Upper Palaeolithic through the Mesolithic and up to the Middle Neolithic, with traces of later occupation. Quite uniquely in the Mediterranean region, deposits at Grotta dell’Uzzo show a continuous occupation during the Mesolithic (*12*). Zooarchaeological and isotopic investigations indicated shifts in the economy and diet of the cave occupants, who subsisted by hunting and gathering for most of the Mesolithic, started to exploit marine resources towards the end of the Mesolithic, and combined all the previous activities with the exploitation of domesticates and increased fishing during the Early Neolithic (*12–14*).

Genome-wide data has been published for six (Epi-)Gravettian HGs and one tentatively Sauveterrian Mesolithic HG from peninsular Italy, and one Late Epigravettian HG from *OrienteC* in Sicily (*15–17*). To date, no ancient genomes are available for Early Neolithic and Late Mesolithic individuals from Sicily or southern Italy. The question of whether the agricultural tradition was adopted by local HGs or brought to Sicily by incoming farmers, thus, remains open.

## Results

Here, we investigated the biological processes underlying the transition from hunting and gathering to agropastoralism in Sicily. We reconstructed the genomes for 19 individuals from Grotta dell’Uzzo dating to a period from the Early Mesolithic ∼8,810 calBCE to the Early Neolithic ∼5,210 calBCE (Data file 1). We obtained a direct accelerator mass spectrometry (AMS) radiocarbon (^14^C) date on the skeletal elements that were used for genetic analysis for 15 individuals, and determined carbon (δ^13^C) and nitrogen (δ^15^N) isotope values from the same bone collagen for dietary reconstruction (Data file 1).

We extracted DNA from bone and teeth in a dedicated clean room, built DNA libraries and enriched for ∼1240k single nucleotide polymorphisms (SNPs) in the nuclear genome and independently for the complete mitogenome (*18*) using in-solution capture (*19*). We restricted our analyses to individuals with evidence of authentic DNA, and removed ∼300k SNPs on CpG islands to minimize the effects of residual ancient DNA damage (*20*). The final data set includes 868,755 intersecting autosomal SNPs for which our newly reported individuals cover 53,352-796,174 SNP positions with an average read depth per target SNP of 0.09-9.39X (Data file 1). We compared our data to a global set of contemporary (*21*) and 377 ancient individuals from Europe, Asia and Africa (*15-17, 21-51*).

### Genetic grouping of the ancient Sicilians

First, we aimed to group the individuals for genetic analysis. For this we co-analysed one Epigravettian HG (*OrienteC*) from the Grotta d’Oriente site on Favignana island in southwestern Sicily (12,250-11,850 calBCE, ^14^C date on charcoal from the deposit (*15, 17*). The ancient Sicilians form three genetic groups that we distinguished based on the individuals’ ^14^C dates (Fig. 1B, Data file 1), position in Principal Component Analysis (PCA, Fig. 1C), mtDNA haplogroups (Supplementary Section S7), and degree of allele sharing in outgroup-*f_3_* statistics and qpWave-based ancestry models (Supplementary Section S2).

**Fig. 1.**
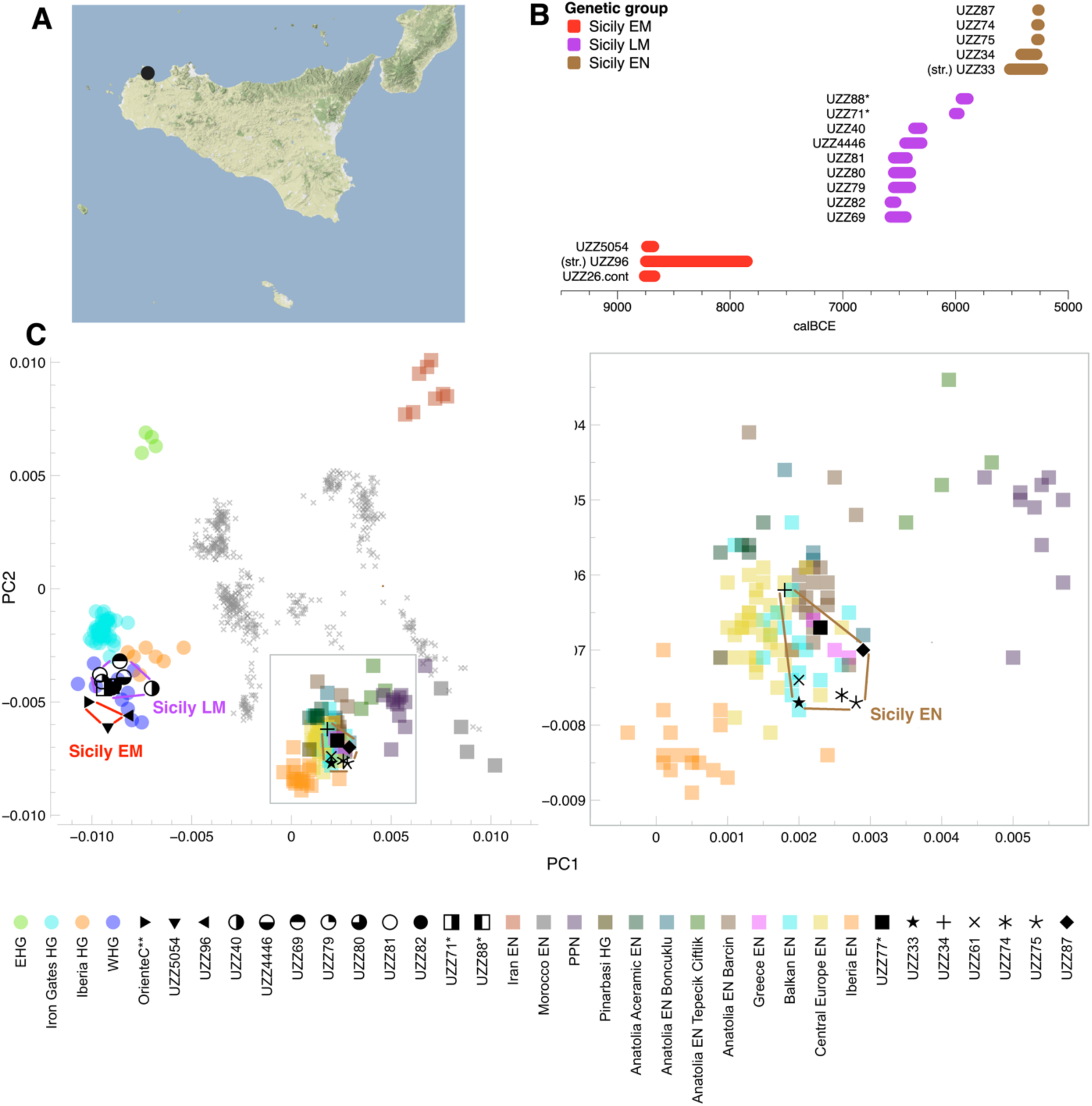
Genetic structure of ancient West Eurasians. **(A)** The geographical location of Grotta dell’Uzzo in Sicily. (**B)** Dating of the ancient Sicilians. Dates were determined from direct radiocarbon (^14^C) measurements, and for *UZZ96* and *UZZ33* from the stratigraphy (str.) (Supplementary Section S1). Dates could not be inferred for individuals *UZZ61*, and *-77*. The individuals fall into three temporal groups, and are coloured according to their assigned genetic group. *UZZ71, -88* and *-77* may be contemporaneous with early Impressa Ware aspects at the site (marked by *). **(C)** PCA plots for the genetic distances between West-Eurasians with principal components constructed from individuals from 43 modern Eurasian groups (grey crosses). The ancient Sicilians are projected (black symbols) together with relevant previously published hunter-gatherers (coloured dots) and early farmers (coloured squares). We co-analyzed an Epigravettian HG from OrienteC (*15*) (marked by **). The genetic variation of the ancient Sicilians forms three genetic groups, that we labelled Sicily EM, LM and EN. The individuals in the Sicily EM and LM genetic group fall close to individuals from the Villabruna cluster that are characterized by high levels of WHG ancestry. In contrast, those from the Sicily EN genetic group contain substantially more Near Eastern ancestry, and fall among early farmers from the Balkan and Central Europe but not from Iberia.

The first two groups consist of individuals that fall close to a cluster of western European Mesolithic hunter-gatherers (‘WHG’) that includes the 14,000-year-old individual from the Villabruna site in northern Italy (‘*Villabruna*’) (*16*) (Fig. 1C). The first and oldest genetic group, which we labelled Sicily Early Mesolithic (Sicily EM, n=3), contains the previously published Epigravettian *OrienteC* (12,250-11,850 calBCE (*15, 17*)) and the two oldest HGs from Grotta dell’Uzzo (∼8,800-8,630 calBCE). These three individuals carried mitogenome lineages that fall within the U2’3’4’7’8’9 branch (Supplementary Section S7, and (*15, 17*) for *OrienteC*). From that haplogroup node they shared nine mutations specific to their lineage and were differently related to each other with regard to three additional private mutations. U2’3’4’7’8’9 mitogenome lineages have already been reported for Upper Palaeolithic European HGs, such as *Paglicci108* associated with the Gravettian in Italy (26,400-25,000 calBCE (*52*)). The second genetic group, which we labelled as Sicily Late Mesolithic (Sicily LM, n=9), contains nine individuals dated to ∼6,750-5,850 calBCE, The mitogenome haplogroups carried by the Sicily LM HGs are U4a2f (n=1), U5b2b (n=2), U5b2b1a (n=1), U5b3(d) (n=3), U5a1 (n=1), and U5a2+16294 (n=1), which are typical for European Late Mesolithic WHGs (*52, 53*).

The third and most recent genetic group, which we labelled as Sicily Early Neolithic (Sicily EN, n=7), contains seven individuals dated to ∼5,460-5,220 calBCE. In PCA, these individuals show substantially Near-Eastern-related ancestry and fall close to early farmers from the Balkans (Croatia, Greece), Hungary, and Anatolia, but not Iberia (Fig. 1C) (*17, 38, 39, 54*). All the individuals in the Sicily EN group, with sufficient coverage for genome reconstruction, carried mitogenome haplogroups characteristic for European early farmers: U8b1b1 (n=2), K1a2 (n=1), N1a1a1 (n=1), J1c5 (n=1) and H (n=1) (*55*).

### Mesolithic substructure and dynamics

Previous research has shown that the genetic diversity among European HGs after the Last Glacial Maximum (LGM) was shaped by various deeply diverged ancestries (*16, 17, 32, 38, 50, 56*). One such ancestry came from a group of pre-LGM individuals dating to ∼30,000 calBCE and associated with the Gravettian industry (Věstonice cluster). Another one was from individuals associated with the Magdalenian industry (El Mirón cluster) that appeared in Europe by ∼17,000 calBCE (*16*). Individuals of the Villabruna cluster, also referred to as western European hunter-gatherers (WHGs), appeared ∼12,000 calBCE throughout continental Europe, and replaced most of the ancestry of the earlier clusters in European HGs. In Mesolithic HGs from eastern Europe (EHGs, ∼6,000 calBCE) and the Iron Gates HGs from southeastern Europe (∼9,500-5,800 calBCE), ancestry related to Upper Palaeolithic Siberians (Ancient North Eurasians, ANE) was found in addition to WHG ancestry.

To visualize the genetic differentiation among the Sicily EM and LM HGs, and their relation to other West Eurasian HGs, we plotted pairwise genetic distances calculated as *f_3_(Mbuti; HG1, HG2)* in a Multidimensional Scaling (MDS) plot (Fig. 2A). The genetic variation among post-LGM European HGs is structured along two clines: 1) a WHG-EHG-ANE cline, confirming the genetic gradient found in Mesolithic HGs from western to eastern Europe and 2) a WHG-GoyetQ2 cline between WHG and Central European Magdalenian-associated individuals on which Iberian HGs take an intermediate position (*15-17, 38, 50, 56*). As previously reported for *OrienteC* (*15*), the Sicily EM HGs *UZZ5054* and *UZZ96* fall at the extreme WHG-end of both ancestry clines, slightly outside the genetic variation of the Villabruna cluster, named after the site name of its oldest representative individual (∼12,230-11,830 calBCE) (Fig. 2A). The position of the Sicily EM HGs on the MDS plot either hints at a WHG ancestry component that is more basal than that found in Villabruna cluster individuals, and/or at substantial genetic drift. Compared to the Sicily EM HGs, the Sicily LM HGs fall closer to the Villabruna cluster in between Sicily EM HGs and Mesolithic Iron Gates HGs (Fig. 2A).

**Fig. 2.**
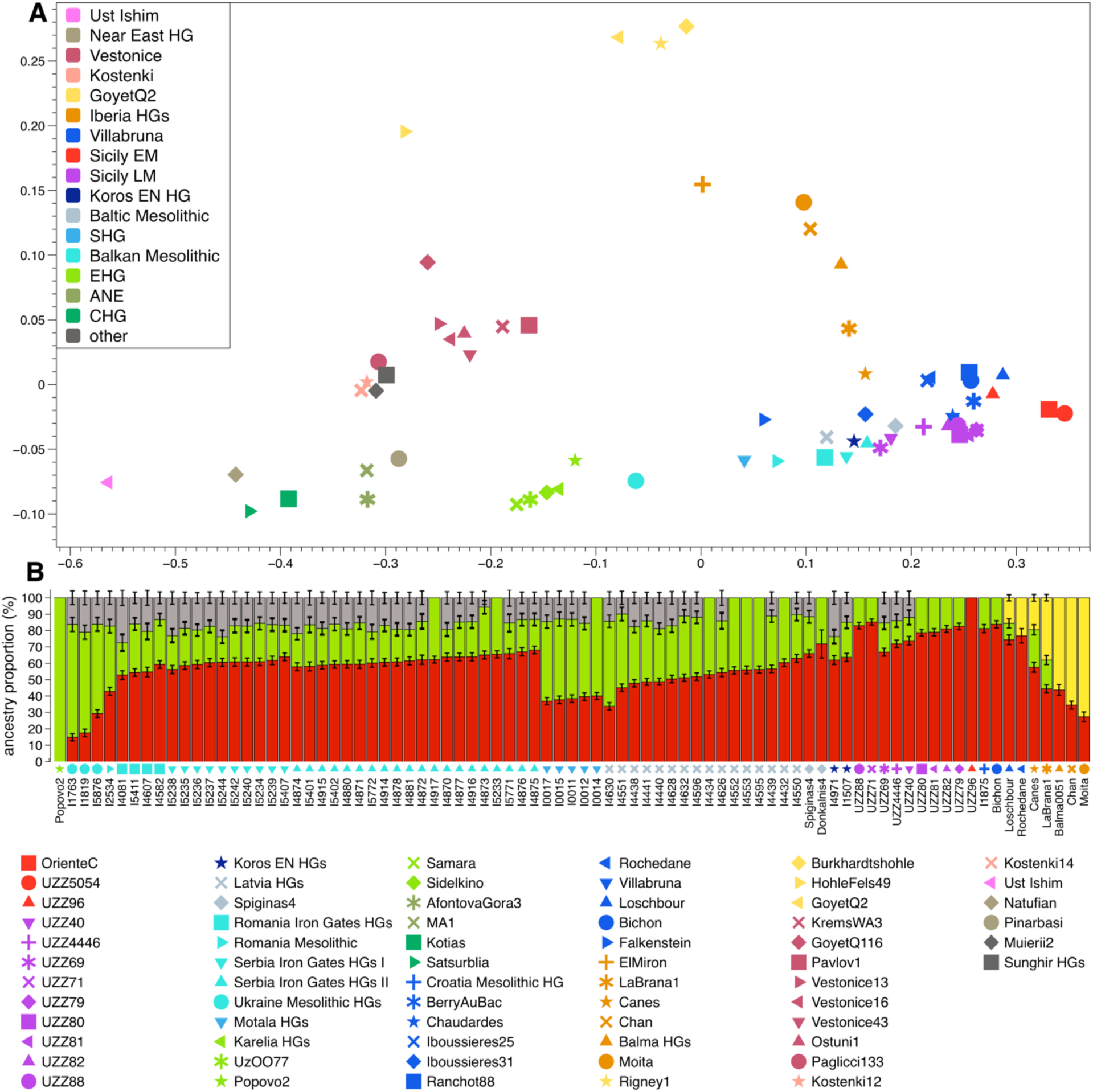
Genetic profile characterization for the Sicilian HGs. **(A)** MDS plot showing structure in the genetic variation among West Eurasian HGs. Genetic distances are based on pairwise *f3*-outgroup statistics of the form *f_3_(Mbuti; HG1, HG2).* Colours reflect various ancestry clusters or geographical groups. The genetic variation among West Eurasian HGs is structured along a WHG-EHG-ANE and WHG-GoyetQ2 ancestry cline. The Sicily EM HGs (red) fall at the extreme WHG end of both ancestry clines. Sicily LM HGs (purple) fall among Villabruna cluster HGs (blue) in between Sicily EM HGs and Mesolithic HGs from the Balkan (turquoise). **(B)** Individual ancestry profiles for Sicily LM HGs and other relevant West-Eurasian HGs. Results are from qpWave- and qpAdm-based admixture models that inferred the ancestry of each target HG as a one-, two- or three-way source mixture of ancestry approximated by Sicily EM (red: *UZZ5054, OrienteC), GoyetQ2* (yellow), EHG (lime green: Karelia HGs, *Uz0077, Samara, Sidelkino*), and *Pınarbaşı* (brown). Sicily EM, *GoyetQ2* and *Pınarbaşı* are taken as proxies for WHG-, Magdalenian- and Near Eastern HG-related ancestry, respectively. Error bars reflect 1 standard error (SE). Individuals with >150k SNPs covered are plotted. Sicily LM HGs, *Bichon* and the Croatia Mesolithic HG contain the highest proportions of Sicily EM ancestry. The Near Eastern-related ancestry is found in some Sicily LM HGs, and frequently among Mesolithic HGs from the Balkan, Baltic and Scandinavia. Congruent to their position in the MDS plot, the Sicily LM HGs ancestry profiles appear intermediate to those of *Bichon* and the Croatia Mesolithic HG (Villabruna cluster), and Mesolithic HGs from the Iron Gates in southeastern Europe.

First, we investigated whether the Sicily EM HGs contain substantial lineage-specific genetic drift. We hence determined the nucleotide diversity (π) by calculating the average proportion of nucleotide mismatches for all possible combinations of individual pairs within the Sicily EM HG group, and compared that to the average of other HG groups. We indeed found significant lower nucleotide diversity for the Sicily EM HGs (95% confidence interval (95CI) π = 0.161-0.170), compared to HGs from Italy from the preceding Upper Palaeolithic (95CI π = 0.227-0.239), and the subsequent Sicily LM HGs (95CI π = 0.217-0.223), and later farmers (Fig. 3 and fig. S3.1). In addition, the nucleotide diversity for the Sicily EM HGs is ∼20% lower compared to contemporaneous HG groups from Central Europe, Iberia and the Iron Gates (Fig. 3). The reduced genetic diversity of Sicily EN HG hence appears to be both geographical- and temporal-specific.

**Fig. 3.**
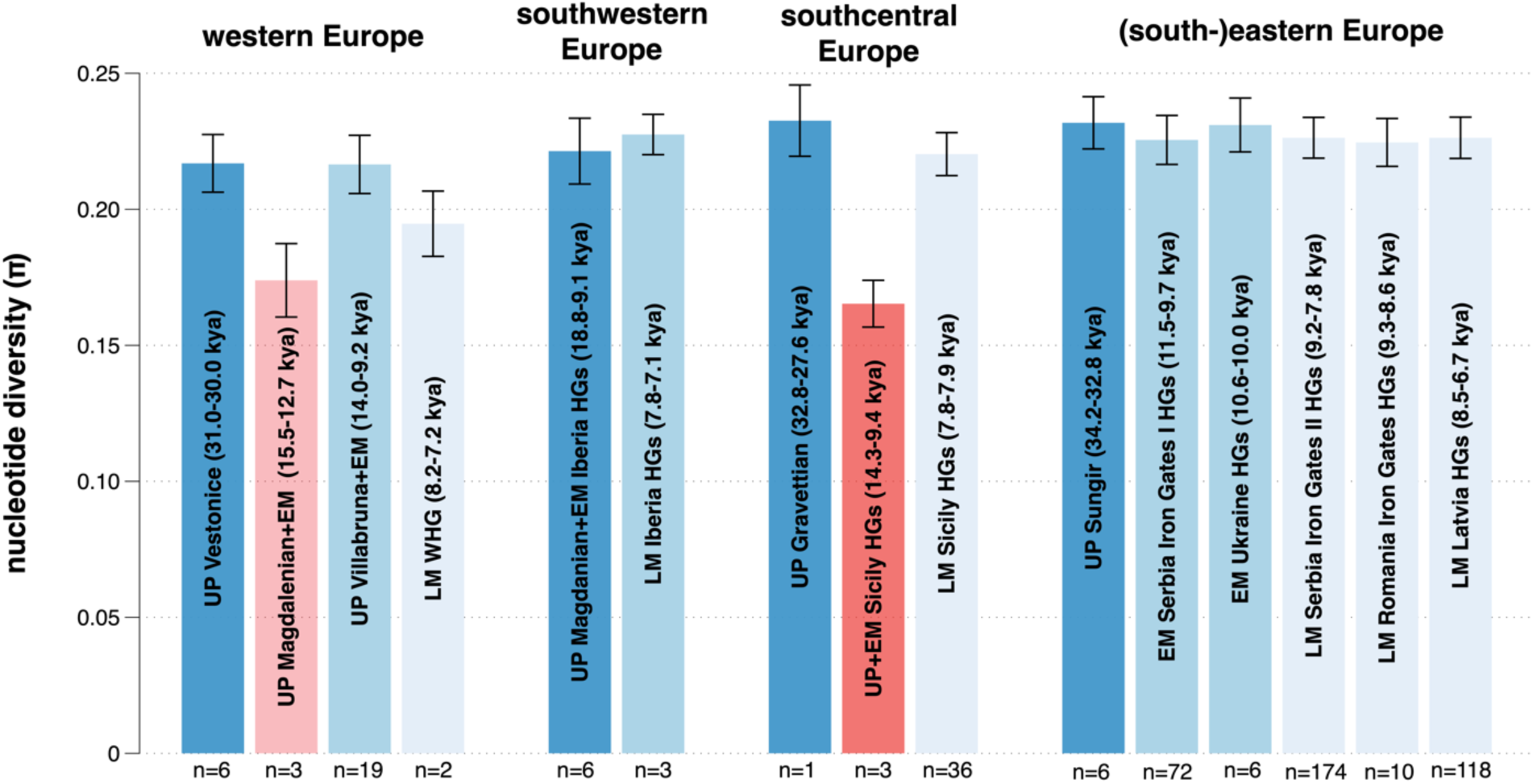
The genetic diversity in Sicilian Early Mesolithic HGs is significantly reduced. Nucleotide diversity (π) was inferred from pseudo-haploid genotypes, calculated as the average proportion of nucleotide mismatches for autosomal SNPs covered in individual pairs within a given HG group. The averages for HG groups from different time periods in different regions in Europe are plotted. UP = Upper Palaeolithic, EM = Early Mesolithic, LM = Late Mesolithic. Individuals are grouped based on their assigned genetic cluster, geographical and temporal proximity (see Data file 1). The number of individual pairs (*n*) that is used to determine the average for each time period is given. Error bars reflect 95% confidence intervals from 5Mb jackknifing. The nucleotide diversity for UP + EM Sicily HGs (red) is significantly lower compared to that for HGs from other time periods or regions in Europe, except for Magdalenian-associated individuals from western Europe (UP Magdalenian + EM: *Rigney1*, *Burkhardshole*, *Hohlefels49, GoyetQ2*).

Secondly, we compared the ancestry component as found in Sicily EM and LM HGs, and their respective affinities to West Eurasian HGs, using an *f_4_-*cladality test of the form *f_4_(Chimp, X; Sicily EM HGs, Sicily LM HGs).* We found a pattern that is linked with geography (Fig. 4A): Sicily EM HGs share significantly more alleles with HGs from western Europe, including Villabruna cluster HGs, and Iberian Upper Palaeolithic and Mesolithic HGs that carry Magdalenian-associated ancestry (*16, 50*). In contrast, the Sicily LM HGs share significantly more alleles with Upper Palaeolithic and Mesolithic HGs from (south-)eastern Europe and Russia, including *AfontovaGora3*, EHGs, *Mal’ta1* and Iron Gates HGs. Notably, comparing the ancestry in Sicily EM HGs with that of *Villabruna* with the cladality statistic *f_4_(Chimp, X; Sicily EM HGs, Villabruna)* results in a similar geographical pattern (fig. S4.3). Also here, Sicily EM HGs share an excess of alleles with western European HGs, including the majority of Villabruna cluster individuals, whereas *Villabruna* does with (south-) eastern European HGs. Fu et al. (*16*) already showed an East Asian affinity for some individuals of the Villabruna cluster individuals compared to older individuals. However, using the Sicily EM HGs as a baseline for WHG ancestry pulls out the difference in genetic affinities to Magdalenian-associated and EHG/ANE-related ancestry more strongly between western and eastern West Eurasian HGs (Supplementary Section S4).

**Fig. 4.**
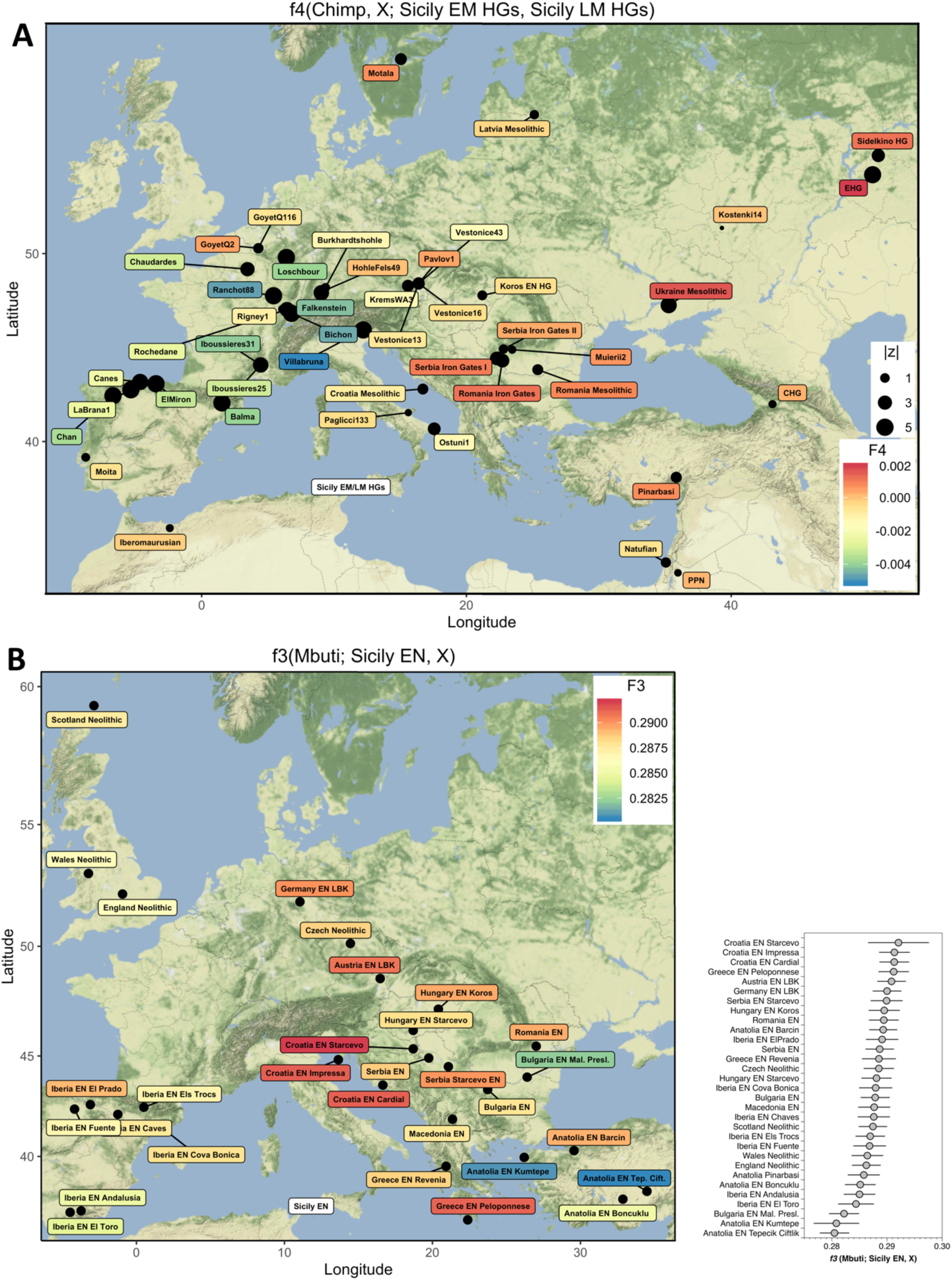
Genomic affinity of the ancient Sicilians. **(A)** Comparing the ancestry in Sicily EM and LM HGs to various West Eurasian HGs (*X*), as measured by *f4(Chimp, X; Sicily EM HGs, Sicily LM HGs*). Cooler colours indicate that *X* shares more genetic drift with Sicily EM HGs than with Sicily LM HGs, and warmer colours indicate the opposite. Dot sizes reflect |z|-scores. Not plotted: *AfontovaGora3* (*f_4_* = 0.0023, z = 3.73), *BerryAuBac* (*f_4_* = 0.0063, z = 5.10). Whereas Upper Palaeolithic and Mesolithic HGs from western Europe, including Villabruna cluster individuals, are genetically closer to Sicily EM HGs, those from eastern Europe are closer to Sicily LM HGs. **(B)** Early Neolithic Sicilian farmers show high genetic affinity to contemporaneous farmers from the Balkan (Croatia and Greece EN Peloponnese), as measured by *f_3_(Mbuti; Sicily EN, X).* Warmer colours indicate higher levels of allele sharing. Error bars in the bar plot indicate 1 SE.

With more explicit modelling using qpGraph (*57*) we further examined the phylogenetic position of Sicily EM HGs. In the least complex scaffold tree Sicily EM HGs and *Villabruna* form a clade on an unadmixed branch, with *GoyetQ2*, a Magdalenian-associated HG, as an outgroup to both of them (fig. S5.3). However, trees that place either *Villabruna* or Sicily EM HGs on an admixed branch with an additional ancestry contribution from *AfontovoGora3* and *GoyetQ2*, respectively, fit the allele frequencies approximately equally well (Supplementary Section S5). Overall, the results suggest that the Sicily EM HGs represent a highly drifted branch closely related to the Villabruna cluster. We can however not rule out that Sicily EM HGs derived an ancestry contribution from Magdalenian-associated individuals or that Sicily EM HGs descended from a more basal lineage that admixed into Iberian HGs and Villabruna cluster individuals (fig. S5.5).

Subsequently, we characterized the ancestry profile of the Sicily LM HGs in more detail. Since the position of Sicily LM HGs on the MDS and PCA plots (Fig. 1D, 2A) is closer to *Villabruna* cluster individuals and EHGs, we investigated whether their gene pool is the result of admixture between the preceding Sicily EM HGs and a group high in EHG-ancestry, or is genetically drifted from Villabruna cluster HGs.

First, we used the outgroup *f_3_*-statistic *f_3_(Mbuti; Sicily LM HGs, X)* to investigate for various West-Eurasian HGs (*X)* which one is genetically closest to Sicily LM HGs (fig. S4.1). Sicily LM HGs shows the highest degree of allele sharing with Sicily EM HGs, followed by other individuals from the Villabruna cluster. Moreover, the statistic *f_4_* (Chimp, Sicily LM HGs; Sicily EM HGs, X) is strongly significantly negative for all tested West-Eurasian HGs, including Villabruna cluster individuals (fig. S4.2). This implies that the Sicily LM HGs form a clade with Sicily EM HGs to the exclusion of other West-Eurasian HGs. Taken together, these statistics indicate substantial continuity in ancestry between the Sicily EM and LM HGs. However, when Sicily EM HGs is used as baseline for the ancestry in Sicily LM HGs in the *f_4_*-statistic *f_4_(Chimp, X; Sicily EM HGs, Sicily LM HGs)*, additional admixture signals are found for various HGs from (south)-eastern Europe and Russia (Fig. 4A). Therefore, Sicily EM HGs do not represent the full gene pool of the Sicily LM HGs.

Subsequently, using qpAdm-based admixture models (*36*) we aimed to more explicitly model the gene pool of the Sicily LM HGs. We found that a three-way ancestry combination of 75.0±1.6% Sicily EM, 15.5±2.4% EHG and 9.5±2.8% Pınarbaşı, a ∼13,300 calBCE HG from central Anatolia (*25*), results in a good fit (P = 0.123, Table 1). Notably, replacing the Sicily EM ancestry with that of *Loschbour* resulted in poorly fitting models (P_Adm_ = 4.27E-09, Table 1). The assigned ancestry components confirm both the substantial continuation of the local Sicily EM ancestry, and the influx of a non-local ancestry frsm (south-)eastern Europe, in Sicily during the Mesolithic. Moreover, the Upper Palaeolithic Pınarbaşı-related ancestry in the Sicily LM HGs is striking, and underlines previous indications for a pre-Neolithic genetic connection between the Near East and European HGs by at least 12,000 calBCE (*16, 17, 25*).

**Table 1.** Overview of key qpWave- and qpAdm-based ancestry models for Sicily EM HGs, Sicily LM HGs and Sicily EN farmers referred to in the main text. For a more comprehensive overview, see Data file 7. For all qpAdm-based admixture models, the P_Wave_-value for the *Sources* is small, indicating that that our used *Outgroups* can distinguish the ancestries between the *Sources*. Note A: The allele frequencies of the Sicily EM and LM HG gene pools do not fully overlap, and hence could not be fitted via one ancestry stream (P_Wave_-value marked by * indicates a model for *Target* and *Source1*). B: A three-way mixture of Sicily EM, EHG and one of various Near Eastern sources results in a full ancestry fit to the Sicily LM HG gene pool. C: Modeling Sicily LM HGs either as a two-way mixture of Sicily EM and EHG or Near Eastern-related ancestry is rejected as a full model fit. D: Replacing the ancestry from EHG with that of Mesolithic Iron Gates HGs marginally improves the model fit. The proportion of assigned Sicily EM ancestry is lower compared to the model that uses EHG as a second ancestry source, due to the substantial amount of WHG ancestry in Iron Gates HGs ((*17*) and see Fig. 2B). E: Replacing the ancestry of Sicily EM with that of *Loschbour* in two or three-way mixtures are strongly rejected as full ancestry fits to the Sicily LM gene pool. F: A two-way mixture of the early farmer ancestry in Anatolia EN Barcin and a West-Eurasian HG source does not adequately fit the Sicily EN gene pool. G: Modelling the early farmer ancestry as a combination of Anatolia EN Barcin and an additional basal ancestry, approximated here by Iran EN Ganj Dareh, and the local preceding Sicilian Mesolithic HGs does result in a full fit to the Sicilian early farmer gene pool. H: The Sicily EN ancestry can also be adequately modelled using a non-local HG ancestry source in addition to early farmer ancestry as approximated by a combination Anatolia EN Barcin and Iran EN Ganj Dareh.

| Target | Source1 | Source2 | Source3 | P<sub>Wave</sub> (Sources) | P<sub>Adm</sub> (Target + Sources) | Source1 (%) | Source2 (%) | Source3 (%) | SE1 (%) | SE2 (%) | SE3 (%) | Note |
| --- | --- | --- | --- | --- | --- | --- | --- | --- | --- | --- | --- | --- |
| Sicily EM | Sicily LM | NA | NA | 7.60E-34* | NA | NA | NA | NA | NA | NA | NA | A |
| Sicily LM | Sicily EM | EHG | Pınarbaşı | 5.70E-113 | 0.12 | 75 | 15.5 | 9.5 | 1.6 | 2.4 | 2.8 | B |
| Sicily LM | Sicily EM | EHG | Natufian | 3.16E-172 | 0.17 | 75.2 | 17.3 | 7.5 | 1.6 | 1.9 | 2.1 |  |
| Sicily LM | Sicily EM | EHG | Iran EN Ganj Dareh | 2.57E-135 | 0.09 | 78 | 15 | 7.1 | 1.4 | 2.7 | 2.2 |  |
| Sicily LM | Sicily EM | EHG | PPN | 3.67E-218 | 0.15 | 76.1 | 16.5 | 7.4 | 1.5 | 2.1 | 2.1 |  |
| Sicily LM | Sicily EM | EHG | NA | 2.19E-299 | 5.56E-03 | 78 | 22 | NA | 1.5 | NA | NA | C |
| Sicily LM | Sicily EM | Pınarbaşı | NA | 1.42E-224 | 4.72E-08 | 76.5 | 23.5 | NA | 1.7 | NA | NA |  |
| Sicily LM | Sicily EM | Natufian | NA | 1.32E-310 | 2.03E-14 | 80.6 | 19.4 | NA | 1.9 | NA | NA |  |
| Sicily LM | Sicily EM | Iran EN Ganj Dareh | NA | 0 | 4.68E-06 | 82.8 | 17.2 | NA | 1.3 | NA | NA |  |
| Sicily LM | Sicily EM | PPN | NA | 0 | 9.44E-11 | 81.4 | 18.6 | NA | 1.5 | NA | NA |  |
| Sicily LM | Sicily EM | Serbia Iron Gates HG I | NA | 7.17E-68 | 0.02 | 37.3 | 62.7 | NA | 2.9 | NA | NA | D |
| Sicily LM | Sicily EM | Serbia Iron Gates HG II | NA | 4.28E-68 | 0.05 | 35.4 | 64.6 | NA | 2.8 | NA | NA |  |
| Sicily LM | Sicily EM | Romania Iron Gates HG | NA | 1.61E-69 | 4.56E-03 | 44.6 | 55.4 | NA | 2.9 | NA | NA |  |
| Sicily LM | Loschbour | EHG | NA | 7.98E-255 | 2.90E-09 | 89.8 | 10.2 | NA | 1.8 | NA | NA | E |
| Sicily LM | Loschbour | EHG | Pınarbaşı | 3.07E-110 | 4.27E-09 | 88.1 | 7.4 | 4.6 | 2 | 2.6 | 3 |  |
| Sicily LM | Loschbour | EHG | Natufian | 1.95E-194 | 1.60E-07 | 86.9 | 6.5 | 6.6 | 1.9 | 2.1 | 2 |  |
| Sicily LM | Loschbour | EHG | Iran EN Ganj Dareh | 4.32E-125 | 1.59E-07 | 89.6 | 3.7 | 6.7 | 1.8 | 2.8 | 2.1 |  |
| Sicily LM | Loschbour | EHG | PPN | 1.20E-213 | 4.15E-08 | 88 | 6.5 | 5.6 | 1.8 | 2.2 | 2 |  |
| Sicily EN | Anatolia EN Barcin | Sicily EM | NA | 0 | 0.02 | 95.7 | 4.3 | NA | 1 | NA | NA | F |
| Sicily EN | Anatolia EN Barcin | Sicily LM | NA | 0 | 0.02 | 94.6 | 5.4 | NA | 1.2 | NA | NA |  |
| Sicily EN | Anatolia EN Barcin | Loschbour | NA | 0 | 7.74E-03 | 95 | 5 | NA | 1.2 | NA | NA |  |
| Sicily EN | Anatolia EN Barcin | Serbia Iron Gates HG I | NA | 0 | 0.01 | 93.8 | 6.2 | NA | 1.4 | NA | NA |  |
| Sicily EN | Anatolia EN Barcin | Serbia Iron Gates HG II | NA | 0 | 0.01 | 93.9 | 6.1 | NA | 1.4 | NA | NA |  |
| Sicily EN | Anatolia EN Barcin | Romania Iron Gates HG | NA | 0 | 0.03 | 92.6 | 7.4 | NA | 1.6 | NA | NA |  |
| Sicily EN | Anatolia EN Barcin | Iran EN Ganj Dareh | Sicily EM | 1.02E-20 | 0.47 | 69.6 | 21.2 | 9.2 | 7.6 | 6.2 | 1.7 | G |
| Sicily EN | Anatolia EN Barcin | Iran EN Ganj Dareh | Sicily LM | 1.15E-24 | 0.34 | 71 | 18.6 | 10.5 | 7.3 | 5.7 | 1.9 |  |
| Sicily EN | Anatolia EN Barcin | Iran EN Ganj Dareh | Loschbour | 7.39E-16 | 0.4 | 63 | 24.8 | 12.2 | 9.1 | 7 | 2.4 | H |
| Sicily EN | Anatolia EN Barcin | Iran EN Ganj Dareh | Serbia Iron Gates HG I | 7.47E-27 | 0.09 | 73.5 | 15.5 | 11 | 7.6 | 5.7 | 2.3 |  |
| Sicily EN | Anatolia EN Barcin | Iran EN Ganj Dareh | Serbia Iron Gates HG II | 1.46E-26 | 0.15 | 72.8 | 16.2 | 11 | 7.6 | 5.7 | 2.2 |  |
| Sicily EN | Anatolia EN Barcin | Iran EN Ganj Dareh | Romania Iron Gates HG | 4.01E-30 | 0.15 | 75 | 13.3 | 11.7 | 7.1 | 5.2 | 2.3 |  |

In a last step, we characterised the ancestry profiles of the Sicily LM HGs on an individual level (Fig. 2B and Fig. 5A). The Sicily LM HGs form a heterogeneous group, with some individuals containing both the EHG and Near Eastern-related ancestry in addition to the preceding local Sicily EM ancestry, whereas others contain solely the additional EHG-related ancestry. The summed proportion for the non-local (EHG + Pınarbaşı) ancestry component in Sicily LM HGs ranges between 17±2% and 33±4% (Fig. 2B, Fig. 5A, Data file 1). Interestingly, whereas a Near Eastern-related ancestry component is discerned in many of the Mesolithic HGs from the Iron Gates and other areas in the Balkan, Baltic or Scandinavia, this is not the case for any of the Villabruna cluster individuals. In contrast to previous statements that the Villabruna cluster individuals form a genetically homogenous group (*16, 25*), here the individuals show rather diverse ancestry profiles with various combinations of Sicily EM, EHG and *GoyetQ2*-related ancestry. Since the Sicily LM HG gene pool contains the distinct Near Eastern-related ancestry but not the *GoyetQ2*-related ancestry, it is unlikely that the diversity of this group originated solely from genetic drift from these Villabruna cluster individuals. We speculate that a single hitherto unsampled population, with an ultimate origin perhaps in the Near East or Caucasus, might harbor the genetic diversity that fits the combined EHG- and Near Eastern-related ancestry in Sicily LM HGs and additional West-Eurasian HGs (e.g. related to the ∼24.000 calBCE Caucasus HGs from Dzudzuana Cave (*56*).

**Fig. 5.**
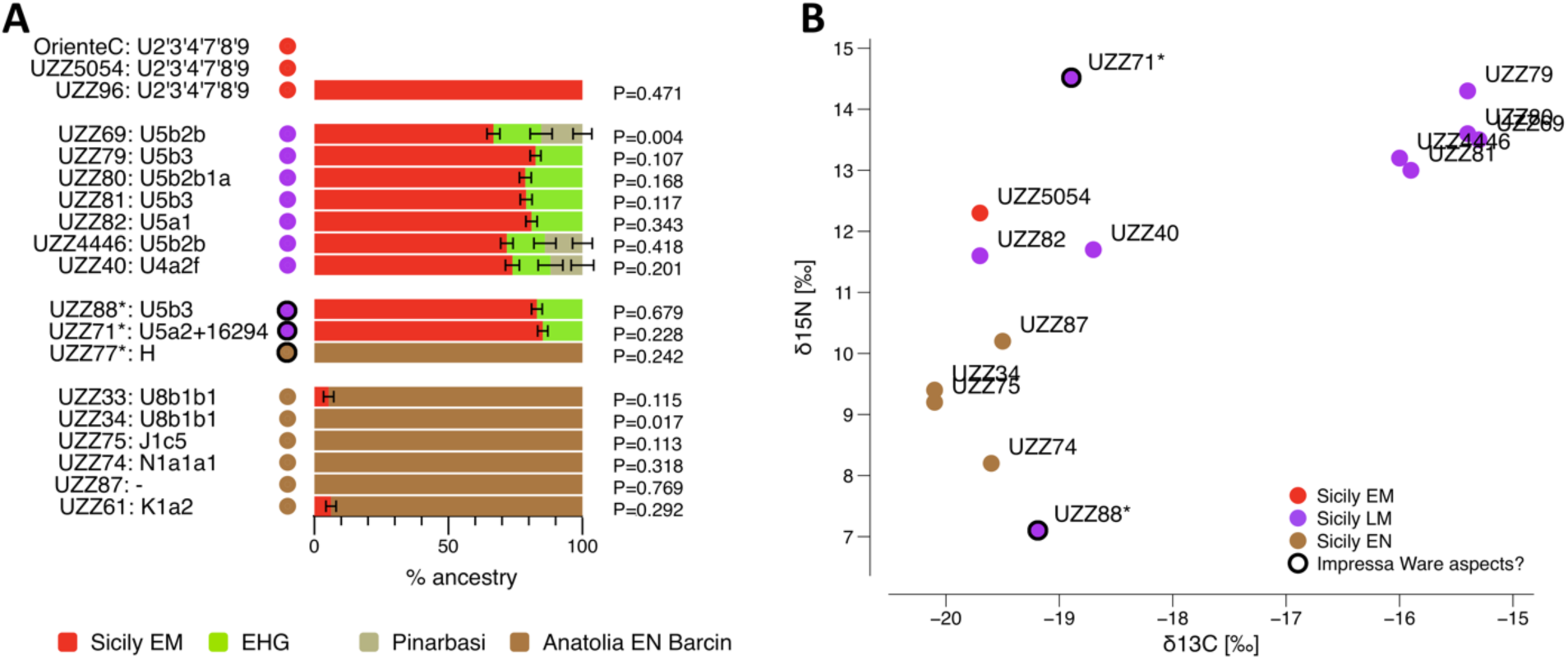
Genomic and dietary turnovers in Sicily during the Mesolithic and Early Neolithic. The coloured dots indicate the individuals’ assigned genetic group with dark outlines those individuals that may be contemporaneous to the earliest Impressa Ware aspects. **(A)** Individual ancestry profiles for the ancient Sicilians determined from qpWave- and qpAdm-based ancestry models. The pre-Neolithic ancestry proportion that is approximated by Sicily EM (*OrienteC/UZZ5054*) is in red, EHG in lime green, Pınarbaşı in light brown, and the early farmer ancestry approximated by Anatolia EN Barcin in dark brown. The 5 cM jackknifing standard errors are marked by horizontal bars. The mitogenomes haplogroups are given. Compared to the preceding Sicily EM HGs, 15-37% of the ancestry in the Sicily LM HGs is from a non-local source that is deeply related to both EHGs and Near Eastern HGs, such as Pınarbaşı. The Sicilian early farmers contain almost entirely early farmer ancestry, indicating an almost complete ancestry replacement during the Early Neolithic. **(B)** Isotope values for diet reconstruction of the ancient Sicilians. The values for the Sicilian early farmers are indicative of a predominantly terrestrial-based farming diet. The Sicily LM HGs associated with the Castelnovian *sensu lato* relied for around half of their protein on seafood. *UZZ71* and *UZZ88*, two individuals that are tentatively contemporaneous to Impressa Ware aspects, show dietary profiles that are strikingly different from the preceding Late Mesolithic Castelnovian HGs and later Early Neolithic Stentinello farmers. *UZZ71* consumed a much higher proportion of freshwater protein, similarly to what is recorded for Mesolithic HGs from the Iron Gates. *UZZ88* consumed more terrestrial plants and less animal protein than the Sicilian early farmers.

### Expanding early farmers replace local HGs in Sicily

Recent ancient human DNA studies have shown that the genetic variation in Early Neolithic groups from central and southwestern Europe is a subset from that found in early farmers from Barcin in northwestern Anatolia and Revenia in northern Greece (*30, 34-37, 39*). Interestingly, early farmers from Diros in Peloponnese Greece might harbour an ancestry component that places them outside of the genetic diversity represented by those from Anatolia Barcin (*17*).

In PCA (Fig. 1C, Data file 1) the Sicilian early farmers (Sicily EN) plot closest to Early Neolithic groups from the Balkan, Serbia, Hungary, Greece and Anatolia, but not from Iberia. With the *f_3_*-outgroup statistic *f_3_(Mbuti; Sicily EN, X)* we aimed to determine the best genetic proxies for the overall gene pool related to Sicily EN. Congruent to the PCA results, the Sicilian early farmers share most genetic drift with various Early Neolithic farmers from the Balkan and Central Europe (Fig. 4B). In addition, in the PCA the Sicilian early farmers do not fall on a cline towards the Sicilian Mesolithic HGs. This suggests that HG ancestry is either absent or low, which would imply a large population replacement in Sicily with the appearance of the Early Neolithic horizons.

To investigate this more formally, we first tested whether Sicilian early farmers contain a distinct HG ancestry component that is not found in early farmers from Anatolia EN Barcin (*38*). For this we used the admixture *f_4_*-statistic *f_4_(Chimp, X; Anatolia EN Barcin, Sicily EN)* that tests for an excess of shared alleles between Sicilian early farmers and various West Eurasian HGs (*X*) when Anatolia EN Barcin is used as a baseline for the early farmer ancestry (fig. S6.1). Sicily EN shows strongly significant signals for admixture for the preceding local Sicily EM (z = 4.12) and Sicily LM HGs (z = 4.09).

To test whether the HG component in Sicily EN is genetically closer to the ancestry of the local preceding Sicily LM HGs or to that of a specific non-local HG source, we performed the *f_4_*-admixture statistic *f_4_(Chimp, Sicily EN; Sicily LM HG, non-local HG)* (fig. S6.2). For HGs from central and southwestern Europe this statistic is negative, indicating that none of them is genetically closer to the HG ancestry in Sicily EN than Sicily LM HGs are. Contrastingly, Mesolithic HGs from southeastern Europe, including Iron Gates HGs, Croatia Mesolithic and Koros EN HGs, do show an excess of shared genetic drift with Sicily EN (fig. S6.2). However, southeastern European HGs and early farmers from Anatolia Barcin and the Balkan share a part of their ancestry (*17, 25*). The excess genetic attraction for southeastern European HGs might hence be driven by the farmer, rather than HG, ancestry component in Sicily EN.

To further investigate what combination of early farmer and HG ancestry fits the Sicily EN gene pool best, and their respective admixture proportions, we used qpWave (P_Wave_) and qpAdm-based (P_Adm_) ancestry models. We required a test result to be more extreme (larger) than P = 0.1 in order not to reject the null-hypothesis of a full ancestry fit (full rank). We found that the ancestry for approximately half of the Sicilian early farmers can be fitted as entirely early farmer ancestry as found in Anatolia EN Barcin, Greece EN Peloponnese, Croatia EN Cardial and Impressa, Hungary EN Koros or Germany LBK (Fig. 4, Data file 7). However, these early farmer sources were rejected as sole ancestry sources to the combined Sicily EN gene pool (max. P_Wave_ ≤ 0.011 for Croatia EN Cardial, Data file 7). We could improve the model fit to the Sicily EN gene pool by adding ∼4-9% ancestry from Sicily LM or Sicily EM HGs as a second source to early farmer ancestry from either Anatolia EN Barcin (P_Adm_ ≥ 0.016), Greece EN Peloponnese (P_Adm_ ≥ 0.059), Croatia EN Cardial (P_Adm_ ≥ 0.056) or Hungary EN Koros (P_Adm_ ≥ 0.300) (Table 1, Data file 7). Notably, three-way mixture models, in which the ancestry from Anatolia EN Barcin was combined with that from early farmers from Ganj Dareh as a proxy for the early farmer ancestry, improved the fit to the Sicily EN gene pool even further (P_Adm_ ≥ 0.335, Table 1). We can therefore not exclude the possibility that the early farmer ancestry in Sicily EN falls partly outside of the broader genetic diversity of the early farmers from Europe and Anatolia Barcin (Supplementary Section S6).

Replacing the local Sicilian HG ancestry with that from Iron Gates HGs or many other Mesolithic European HGs often resulted in similar fits (Table 1, Data file 7). To the limits of the genetic resolution, we hence could not accurately discern whether the Sicilian early farmers derived their HG ancestry from a Mesolithic local Sicilian HG source or from a geographically more distant one. However, a local contribution appears to be the most plausible explanation, which is in line with the hypothesized continuity in occupation at Grotta dell’Uzzo (*14, 58–60*).

## Discussion

Southern Italy has long been viewed as a southern refugium (*61, 62*) during the LGM, ∼25,000 years ago, from where Europe was repopulated (*16, 52, 63*). The earliest evidence for the presence of *Homo sapiens* in Sicily dates to ∼17,000-16,000 calBCE, following the time when a land bridge connected the island to peninsular Italy (*64, 65*). Some Late Epigravettian sites in Sicily contain rock panels with engraved animal figures, which are indistinguishable from those of the Franco-Cantabrian style typical of the Magdalenian (*66*). Moreover, the presence of some decorated pebbles at sites in northwestern Sicily is also suggestive of cultural links with the Azilian in the French Pyrenees (*66*). Although there are many sites in peninsular Italy and Sicily with evidence of Late Upper Palaeolithic occupation, this decreases during the Mesolithic (*67*). The Early Mesolithic HGs from Grotta dell’Uzzo analysed here produced a lithic industry of Epigravettian tradition ((*68*), Supplementary Section S1) and, in continuity with their predecessors, subsisted mainly by hunting large terrestrial game with important contributions of plant foods and limited consumption of marine resources (*12*) (Fig. 5B). Here we showed that, compared to the ∼12,000 calBCE *Villabruna* individual, the Sicily EM HGs have a higher genetic affinity to Magdalenian-associated Iberian HGs *El Miron* and *Balma Guilanyà.* Given the profoundly reduced genetic diversity in the Sicily EM HGs and their closely related non-identical mitogenomes in the U2’3’4’7’8’9 branch, we speculate that these individuals are unadmixed descendants from a peri-glacial refugium population.

The subsequent ∼6,750-5,850 calBCE Late Mesolithic HGs derive between 15-37% of their ancestry from an EHG-related source with an affinity to the Near East. The substantial influx of non-local ancestry indicates a large genetic turnover in Sicily during the Mesolithic, and is matched by a change in diet that is characterized by a statistically significantly higher intake of marine-based protein (Fig. 5B). The seven oldest individuals in this group (dated ∼6,750-6,250 calBCE) are tentatively assigned to the Castelnovian *sensu lato* facies (*12, 14, 69*) (Supplementary Section S1). The Castelnovian is part of the pan-European Late Mesolithic blade and trapeze lithic complex, and appeared throughout Italy ∼6,800-6,500 calBCE ((*70*), D. Binder personal communication). These lithic industries have been argued to originate from the Circum Pontic area (*71, 72*), with a possible ultimate origin from as far as eastern Asia (*73, 74*), or alternatively from the Capsian culture in northwestern Africa (*75, 76*). The ancestry profiles of the individuals associated with the Castelnovian *sensu lato* show a similarity to those of Mesolithic HGs from the Iron Gates, eastern Europe and the Baltic, hence providing support for a connection to the East.

The Sicilian early farmers carried almost exclusively ancestry characteristic for early European farmers. The preceding Late Mesolithic HGs may have contributed only a maximum of ∼7% ancestry (Fig. 5A). Six individuals in this group are from layers that chronologically coincide with the presence of Early Neolithic Stentinello Wares, and one individual (*UZZ77*, undated) tentatively with older aspects of Impressa Ware (Supplementary Section S1). The isotope values of the Stentinello Ware associated early farmers are congruent with them having a terrestrial-based farming diet (Fig. 5B). Their ancestry composition points to a full-scale demographic transition during the Early Neolithic, similar to the Mesolithic-Neolithic transition in other regions in Europe (*30, 34, 36, 37, 53, 54, 77*). Intriguingly however, two individuals *UZZ71* and *UZZ88*, dated ∼6,050-5,850 calBCE, chronologically coincide with layers at the site that may contain the very first aspects of Impressa Wares (*12*). These two individuals fall fully within the genetic diversity of the Late Mesolithic HGs associated with the Castelnovian *sensu lato*, despite postdating them by ∼200 years (Fig. 1C, 5A, Supplementary Section S1, S2). Both these individuals show isotope values that are strikingly different from both the later Sicilian Early Neolithic farmers associated with Stentinello pottery and preceding Late Mesolithic HGs (Fig. 5B, Data file 1). The diet of the *UZZ71* individual included a large proportion of freshwater protein, similarly to what is recorded for Mesolithic hunter-gatherers from the Iron Gates on the Balkan peninsula (e.g. (*78, 79*)). On the other hand, *UZZ88* has an isotopic composition (δ^13^C = −19.2‰, δ^15^N = 7.1‰) that suggests a terrestrial-based farming diet with very low levels of animal protein consumption and that is significantly different from the hunter-gatherers it descended from (mean δ^13^C = −16.6±1.8‰, mean δ^15^N = 13.0±1.0‰), who relied for around half of their protein on seafood. Given the distinct diets and the intermediate ^14^C dates, these two individuals might provide tentative initial evidence that HGs adopted elements of farming in Sicily, as was hypothesized by Tusa (*59, 60*). However, more extensive research on the stratigraphy of Grotta dell’Uzzo is necessary to determine whether the Impressa Ware aspects are indicative of a transitional period (*80*) or should be considered intrusive (*81*).

Although individuals that blur the Mesolithic and Neolithic dichotomy are rather rare, they have been reported before (*17, 29, 78*). Taken together, these individuals could indicate that hunter-gatherers may have met early farmers in different areas of the Mediterranean for which the frequency and exact geographical interaction sphere remains to be unravelled.

## Supporting information

Supplementary Sections S1-7

## Materials and Methods

### aDNA analysis

All pre-amplification laboratory work was performed in dedicated clean rooms (*82*) at the Max Planck Institute (MPI) for the Science of Human history (SHH) in Jena and MPI for Evolutionary Anthropology (EVA) in Leipzig, Germany. At the MPI-SHH the individuals were sampled for bone or tooth powder, originating from various skeletal elements (e.g. petrous, molars, teeth, humerus, phalange, tibia, see Data file 1). The outer layer of the skeletal elements was removed with high-pressured powdered aluminium oxide in a sandblasting instrument, and the element was irradiated with ultraviolet (UV) light for 15 minutes on all sides. The elements were then sampled using various strategies, including grinding with mortar and pestle or cutting and followed by drilling into denser regions (Data file 1). Subsequently, for each individual 1-8 extracts of 100uL were generated from ∼50mg powder per extract, following a modified version of a silica-based DNA extraction method (*83*) described earlier (*50*) (Data file 1). At the MPI-SHH, 20uL undiluted extract aliquots were converted into double-indexed double stranded (ds-) libraries following established protocols (*40, 84*), some of them with a partial uracil-DNA glycosylase (‘ds UDG-half’) treatment (*85*) and others without (‘ds non-UDG’). At the MPI-EVA, 30uL undiluted extract aliquot was converted into double-indexed single-stranded (ss-) libraries (*86*) with minor modifications detailed in (*87*), without UDG treatment (‘ss non-UDG’) (Data file 1). At the MPI-SHH, all the ds- and ss-libraries were shotgun sequenced to check for aDNA preservation, and subsequently enriched using in-solution capture probes following a modified version of (*19*) (described in (*25*)) for ∼1240k single nucleotide polymorphisms (SNPs) in the nuclear genome (*17, 18, 54*) and independently for the complete mitogenome. Then the captured libraries were sequenced on an Illumina 224 HiSeq4000 platform using either a single end (1×75bp reads) or paired end configuration (2×50bp reads).

The sequenced reads were demultiplexed according to the expected index pair for each library, allowing one mismatch per 7 bp index, and subsequently processed using EAGER v1.92.21 (*88*). We used AdapterRemoval v2.2.0 (*89*) to clip adapters and Ns stretches of the reads. We merged paired end reads into a single sequence for regions with a minimal overlap of 30 bp, and single end reads smaller than 30 bp in length were discarded. The reads obtained from the nuclear capture were aligned against the human reference genome (hg19), and those from the mitogenome captured against the revised Cambridge Reference Sequence (rCRS). For mapping we used the Burrows-Wheeler Aligner (BWA v0.7.12) *aln* and *samse* programs (*90*) with a lenient stringency parameter of ‘-n 0.01’ that allows more mismatches, and ‘-l 16500’ to disable seeding. We excluded reads with Phred-scaled mapping quality (MAPQ) <25. Duplicate reads, identified by having identical strand orientation, start and end positions, were removed using DeDup v.0.12.1 (*88*).

### aDNA authentication and quality control

We assessed the authenticity and contamination levels in our ancient DNA libraries (unmerged and merged per-individual) in several ways. First, we checked the cytosine deamination rates at the end of the reads (*91*) using DamageProfiler v0.3 (https://github.com/apeltzer/DamageProfiler). After merging the libraries for each individual, we observed 21-52% C>T mismatch rates at the first base in the terminal nucleotide at the 5’-end, an observation that is compatible with the presence of authentic ancient DNA molecules. Second, we tested for contamination of the nuclear genome in males based on the X-chromosomal polymorphism rate. We determined the genetic sex by calculating the X-rate (coverage of X-chromosomal SNPs/ coverage of autosomal SNPs) and Y-rate (coverage of Y-chromosomal SNPs/ coverage of autosomal SNPs) (*16*). Four individuals for which the libraries showed a Y-rate ≥ 0.49 we assigned the label ‘male’ and 14 individuals with Y-rates ≤ 0.07 as ‘female’. The individual *UZZ26.cont* with an intermediate Y-rate of 0.17 we excluded from further genetic analyses. Then we tested for heterozygosity of the X-chromosome using ANGSD v0.910 (*92*) (≥ 200 X-chromosomal SNPs, covered at least twice (*16*). We found a nuclear contamination of 1.7-5.3% for the four male individuals (Data file 1), based on new Method1 (*93*). Third, we obtained two mtDNA contamination estimates for genetic males and females, using ContaMix v1.0.10 (*19*) and Schmutzi v1.0 (*94*) (Data file 1). Before running Schmutzi, we realigned the reads to the rCRS using CircularMapper v1.93.4 filtering with MAPQ < 30. After removing duplicate reads, we downsampled to ∼30,000 reads per library. With Schmutzi we found low contamination estimates of 1-3% for all individuals with sufficient coverage (Data file 1). ContaMix returned estimates in the range of 0.0-5.6% for all individuals except for *UZZ69* (3.7-10.6%) and the lower coverage individual *UZZ096* (0.3-13.5%).

### Dataset

For genotyping we extracted reads with high mapping quality (MAPQ ≥ 37) to the autosomes using samtools v1.3. The DNA damage plots indicated that misincorporations could extend up to 10 bp from the read termini in non-UDG treated and up to 3bp in UDG-half treated libraries. We hence clipped the reads accordingly, thereby removing G>A transitions from the terminal read ends in ds-libraries and C>T transitions in both ss- and ds-libraries. For each individual, we randomly chose a single base per SNP site as a pseudo-haploid genotype with our custom program ‘pileupCaller’ (https://github.com/stschiff/sequenceTools). We intersected our data with a global set of high-coverage genomes from the Simon Genome Diversity Project (SGDP) for ∼1240k nuclear SNP positions (*21*), including previously reported ancient individuals from (*15-17, 21-51*). To minimize the effects of residual ancient DNA damage, we removed ∼300k SNPs on CpG islands from the data set. CpG dinucleotides, where a cytosine is followed by a guanine nucleotide, are frequent targets of DNA methylation (*95*). Post-mortem cytosine deamination was shown to occur more frequently at methylated than unmethylated CpGs (*20*) resulting in excess of CpG → TpG conversions. The final data set includes 868,755 intersecting autosomal SNPs for which our newly reported individuals cover 53,352-796,174 SNP positions with an average read depth per SNP of 0.09-9.39X (Data file 1). For principal component analyses (PCA) we intersected our data and published ancient genomes with a panel of worldwide present-day populations, genotyped on the Affymetrix Human Origins (HO) (*37, 57*). After filtering out CpG dinucleotides this data set includes 441,774 SNPs.

### Kinship relatedness and individual assessment

We determined pairwise mismatch rates (PMMRs) (*34, 96*) for pseudo-haploid genotypes to check for genetic duplicate individuals and first-degree relatives. If two individuals show similar low PMMRs for inter- and intra-individual library comparisons, then this indicates a genetic duplicate. Moreover, the expected PMMR for two first-degree related individuals falls approximately in the middle of the baseline values for comparison between genetically unrelated and identical individuals (*97*). We found a genetic triplicate (UZZ44, -45, -46) and quintuplicate (UZZ50-54), and merged the respective libraries into *UZZ4446* and *UZZ5054*, respectively. In addition, *UZZ79* and *UZZ81* showed an elevated PMMR indicative of a kinship relation (Data file 4).

### Mitogenome haplogroup determination

We could reconstruct the mitochondrial genomes for 17 individuals (Data file 1). To obtain an automated mitochondrial haplogroup assignment we imported the consensus sequences from Schmutzi into HaploGrep2 v2.1.1 ((*98*); available via: https://haplogrep.uibk.ac.at/) based on phylotree (*99*) (mtDNA tree build 17, available via http://www.phylotree.org/). In parallel, we manually haplotyped the reconstructed mitogenomes, based on a procedure described in (*52*). We imported the bam.files for the merged libraries into Geneious v.9.0.5 (http://www.geneious.com) (*100*). After reassembling the reads against the revised Cambridge Reference Sequence (rCRS) we called SNP variants with a minimum variant frequency of 0.7 and 2.0X coverage. Using phylotree, we double-checked whether the called variants matched the expected diagnostic ones based on the automated HaploGrep assignment. We did not consider known unstable nucleotide positions 309.1C(C), 315.1C, AC indels at 515-522, 16093C, 16182C, 16183C, 16193.1C(C) and 16519. We extracted the consensus sequences based on a minimum of 75% base similarity. Using this approach, we identified a total of twelve lineage-specific and private variants in the high coverage *UZZ5054* mitogenome. Four of the lineage-specific variant positions were covered by only one or two reads in the low coverage *UZZ96* and *OrienteC* genomes, and hence fell initially below our frequency threshold for variant detection. However, since these variants were covered by a large number of reads in the closely related *UZZ5054* mitogenome, for *UZZ96* and *OrienteC* we based the variant calls at these positions on the few reads available and adjusted their consensus sequences accordingly (table S7.3).

### Y-chromosome haplogroup determination

To determine the Y chromosome haplogroup for genetic males we used the *yHaplo* program ((*101*), available via: https://github.com/23andMe/yhaplo). We based our haplogroup assignment on 13,581 strand-unambiguous ancestry informative SNPs from the ISOGG (International Society of Genetic Genealogy) data set. We called genotypes for these SNP sites by randomly choosing one allele with ‘pileupCaller’ (https://github.com/stschiff/sequenceTools). Using an in-house script (Choongwon Jeong) the genotypes were converted to an input format for *yHaplo*. In ancient genomes, missing data for key diagnostic sites may halt the automated search before the most derived haplogroup is reached. We therefore manually checked the coverage for informative SNPs further downstream for the haplogroup that was assigned to each individual. Also, our sequence reads from the non-UDG treated ss-libraries have an expected 3-5% C<->T mismatches due to residual ancient damage. C<->T mismatches at diagnostic SNP positions may result *yHaplo* to incorrectly assign a more derived haplogroup. We therefore checked whether the ancestral state variants matched those that are expected for the assigned haplogroup.

### Principal component analysis (PCA)

We computed principal components from individuals from 43 modern West Eurasian groups in the Human Origin panel (*37, 57*) using the *smartpca* program in the EIGENSOFT package v6.0.1 (*102*) with the parameter ‘numeroutlieriter:0’. Ancient individuals were projected using ‘lsqproject:YES’ and ‘shrinkmode:YES’.

### *f*-statistics

We performed *f*-statistics on the 1240k data set using ADMIXTOOLS (*57*). For *f_3_*-outgroup statistics (*103*) we used *qp3Pop* and for *f_4_*-statistics *qpDstat* with f4mode:YES. Standard errors (SEs) were determined using a weighted block jackknife over 5Mb blocks. *F_3_*-outgroup statistics of the form *f_3_(O;A,B)* test the null hypothesis that *O* is a true outgroup to *A* and *B*. The strength of the *f_3_*-statistic is a measure for the amount of genetic drift that *A* and *B* share after they branched off from a common ancestor with *O*, provided that *A* and *B* are not related by admixture. *F_4_*-statistics of the form *f_4_(X,Y; A,B)* test the null hypothesis that the unrooted tree topology ((*X*,*Y*)(*A*,*B*)), in which (*X*,*Y*) form a clade with regard to (*A*,*B*), reflects the true phylogeny. A positive value indicates that either *X* and *A*, or *Y* and *B*, share more drift than expected under the null hypothesis. A negative value indicates that the tree topology under the null-hypothesis is rejected into the other direction, due to more shared drift between *Y* and *A*, or *X* and *B*.

### Multidimensional scaling (MDS)

We performed MDS using the R package *cmdscale*. Euclidean distances were computed from the genetic distances among West-Eurasian HGs, as measured by *f_3_(Mbuti; HG1, HG2)* for all possible pairwise combinations (*16*). The first two principal components are plotted. We restricted the analyses to individuals with >30,000 autosomal SNPs covered. Relevant previously published West Eurasian HGs were pooled in groups according to their geographical or temporal context, following their initial publication labels (Data file 1). We deviated from the original labels with regard to Iron Gates HGs from Serbia by splitting them into an early and late subgroup, labelled here as ‘Iron Gates HG Serbia I’ (RC date: 10,000-7,500 calBCE) and ‘Iron Gates HG Serbia II’ (RC date: 7,500-5,700 calBCE) (*17*).

### Nucleotide diversity

We selected a total of 120 West-Eurasian HGs with >100k SNPs covered, of which 103 were previously published (Data file 1), from four broad geographical regions “western” (n=18), “south-western” (n=7), “southern-central” (n=33), and “(south)-eastern” (n=62) Europe. We subgrouped the individuals further based on similar ^14^C-dates and genetic cluster assignment (for an overview of the HG groups, see (Data file 1). E.g. we analysed the individuals associated with the Villabruna cluster and those high in Magdalenian-related ancestry in separate groups. We determined the nucleotide diversity (π) from pseudo-haploid genotypes by calculating the proportion of nucleotide mismatches for overlapping autosomal SNPs covered by at least one read in both individuals. We hence determined π from all possible combinations of individual pairs, rather than from all possible chromosome pairs, within a given group. We filtered out individual pairs that shared less than 30,000 SNPs covered. We calculated an average over all the individual pairs within a group and determined standard errors from block jackknifes over 5Mb windows and 95% confidence intervals (95CIs) from 1,000 bootstraps

### Inference of mixture proportions

To characterize the ancestry of the ancient Sicilians we used the qpWave (*54*) and qpAdm (*36*) programs from the Admixtools v3.0 package, with the ‘allsnps: YES’ option. qpWave tests whether a set of *Left* populations is consistent with being related via as few as N streams of ancestry to a set of *Outgroup* populations. qpAdm tries to fit a *Target* as a linear combination of the *Left/Source* populations, and estimates the respective ancestry proportions that each of the *Left* populations contributed to the *Target*. Both qpWave and qpAdm are based on *f_4_*-statistics of the form *f_4_(X,O1;O2,O3)*, where *O1, O2, O3* are all the triplet combinations of the *Outgroup* populations, and *X* is a *Target* or *Left/Source* population. We used an *Outgroup* set from Mathieson et al. (*17*): *El Miron*, *Mota*, Mbuti, *Ust Ishim, Mal’ta, AfontovaGora3, GoyetQ116, Villabruna, Kostenki14, Vestonice16*, Karitiana, Papuan, Onge. We grouped individuals with a similar ancestry for Sicily EM, Sicily LM, EHG and CHG (Data file 1). Since missing data may inflate the P-values for this test, we required a test result to be smaller (less extreme) than P = 0.1 in order to reject the null-hypothesis of a full ancestry fit between the *Target* and the *Left/Source* population(s). Prior to running qpAdm we used qpWave to check whether the pairs of *Left/Source* populations are not equally related to the *Outgroups*.

### Phylogeny modelling

We used the qpGraph program (*57*) to construct a phylogeny of ancestry lineages found among Palaeolithic and Mesolithic West-Eurasian HGs to clarify the genetic history of Sicily EM and LM HGs. For our modelling we used the parameters ‘useallsnps: YES’, ‘forcezmode: YES’, ‘terse: NO’. We built the phylogeny models with increasing complexity by fitting representative West Eurasian HG ancestry lineages in the order: 1) Mbuti, 2) *Ust Ishim*, 3) *Kostenki14*, 4) *Mal’ta*, 5) *Vestonice16*, 6) *GoyetQ2*, 7) *AfontovaGora3*, 8) *Villabruna* or Sicily EM, 9) Sicily EM or *Villabruna* (Supplementary Section S5). We fitted the lineages one by one by adding them to all possible nodes as a branch without admixture or as a binary admixture between two branches. We preferred the former over the latter if both models fitted the observed *f-*statistics equally well. We selected models that did not include trifurcations or 0% ancestry stream estimates, and for which the difference between the observed and fitted *f-*statistics were less extreme than 3.5 SEs.

### Direct AMS ^14^C bone dates

For 15 individuals we obtained a direct ^14^C date from the skeletal element that was used for genetic analysis (Data file 1). All bone samples were pretreated at the Department of Human Evolution at the Max Planck Institute for Evolutionary Anthropology (MPI-EVA), Leipzig, Germany, using the method described in (*104*). For each skeletal element, 200-500mg of bone/tooth powder was decalcified in 0.5M HCl at room temperature ∼4 hours until no CO_2_ effervescence was observed. To remove humics, in a first step 0,1M NaOH was added for 30 minutes, followed by a final 0.5M HCl step for 15 minutes. The resulting solid was gelatinized following a protocol of (*105*) at pH 3 in a heater block at 75°C for 20h. The gelatin was then filtered in an Eeze-Filter™ (Elkay Laboratory Products (UK) Ltd.) to remove small (> 80mm) particles, and ultrafiltered (*106*) with Sartorius “VivaspinTurbo” 30 KDa ultrafilters. Prior to use, the filter was cleaned to remove carbon containing humectants (*107*). The samples were lyophilized for 48 hours. In order to monitor contamination introduced during the pre-treatment stage, a sample from a cave bear bone, kindly provided by D. Döppes (MAMS, Germany), was extracted along with the batch from La Ferrassie samples (*108*). In marine environments the radiocarbon is older than the true age, usually by ∼400 years (marine reservoir effect). The specimens for which a correction was necessary are *UZZ4446* (40±10% marine), *UZZ81* (45±10% marine), *UZZ69*, *UZZ79* and *UZZ80* (50±10% marine). Corrections were made using the reservoir correction estimated for the Mediterranean Basin by (*109*), which is ΔR = 58±85 ^14^C yr.

### Isotope analysis

For 14 individuals we determined the carbon (δ^13^C) and nitrogen (δ^15^N) isotope values for dietary inference (Data file 1). To assess the preservation of the collagen yield, C:N ratios, together with isotopic values are evaluated following the limits of (*110*).

## H2: Supplementary Materials

Section S1. Grotta dell’Uzzo: archaeology and stratigraphic sequence

Section S2. Genetic grouping and substructure of the ancient Sicilians

Section S3. Elevated lineage-specific genetic drift in the Sicilian Early Mesolithic

Section S4. Characterizing the Sicilian Mesolithic HGs ancestry using F-statistics

Section S5. Investigating the phyologenetic position of the Early Mesolithic Sicilian HGs

Section S6. Characterizing the Sicilian early farmer ancestry using F-statistics

Section S7. Uniparental marker haplotyping

Data file S1. Summary table results ancient Sicilians, labels used in analyses, data in Supplementary Sections

Data file S2. Data underlying Fig. 1

Data file S3. Data underlying Fig. 2

Data file S4. Data underlying Fig. 3

Data file S5. Data underlying Fig. 4

Data file S6: Data underlying Fig. 5.

Data file S7. Admixture models for ancient Sicilians and West Eurasian HGs

## Acknowledgements

For helpful comments we thank Stephanie Eisenmann, Maite Rivollat, Thiseas Lamnidis and other members of the Department of Archaeogenetics, and Barbara Pavlek of the Minds & Traditions research group of the Max Planck Institute for the Science of Human History. We are grateful for comments on the manuscript from Didier Binder from the French National Center for Scientific Research and Detlef Gronenborn from the Leibniz Research Institute for Archaeology. We thank David Reich, Shop Mallick and Ian Mathieson for access to unpublished data. We thank Sven Steinbrenner of the Department of Human Evolution from the Max Planck Institute for Evolutionary Anthropology for undertaking the stable isotope analysis.

## Author contributions

M.Ma, W.H, and J.K conceived the study. M.Ma, A.T, M.P, S.T, C.C, V.S and R.dS provided the ancient human remains and input for the archaeological interpretation. M.vdL, C.F, S.N and L.K performed laboratory work with the help of F.A, M.B, R.R, R.S, A.W, G.B and M.Me. M.vdL conducted the population genetic analyses with the help of K.P, C.J and W.H. S.T performed the AMS radiocarbon dating analysis, and M.Ma the isotope analysis. M.vdL, M.Ma, W.H, V.V-M, C.P, S.S, C.J, K.P and J.K. wrote the paper with input from all co-authors.

## Data and materials availability

Genomic data (BAM format) are available through the Sequence Read Archive (accession number X) and consensus mitogenome sequences (FASTA format) in GenBank (accession numbers X to X).

