## Supplementary Sections S1-7 for "Genomic and dietary transitions during the Mesolithic and Early Neolithic in Sicily"

S1. Grotta dell'Uzzo: archaeology and stratigraphic sequence …………………………..……… p.1

S2. Genetic grouping and substructure of the ancient Sicilians ……………………….............. p.10

S3. Elevated lineage-specific genetic drift in the Sicilian Early Mesolithic HGs ……..………… p.13

S4. Characterizing the Sicilian Mesolithic HGs ancestry using F-statistics …………….......... p.18

S5. Investigating the phyologenetic position of the Early Mesolithic Sicilian HGs ………...... p.20

S6. Characterizing the Sicilian early farmer ancestry using F-statistics …………….……..... p.26

S7. Uniparental marker haplotyping ……………………………………………….…….. p.32

**S1: Grotta dell'Uzzo: archaeology and stratigraphic sequence**

**1. The site, its burial ground and human remains**

Grotta dell'Uzzo is a large shelter-like cave located in northwestern Sicily, along the eastern cliffs of the San Vito lo Capo peninsula (fig. S1.1). The site was visited hastily in 1927 by the French archaeologist Raymond Vaufrey, who did not realize its importance. The discovery of the deposit and its stratification was made in the early 1970s by Giovanni Mannino (*111*), who excavated a small test trench in the cave, exposing a sequence of *in situ* Mesolithic deposit (identified as *epipaleolitico*). Prehistoric deposits have been excavated during the 1970s, 1980s and in 2004 within a number of trenches both inside and outside the overhang of the cave (fig. S1.2). This revealed that the site was occupied from at least the late Upper Palaeolithic through the Mesolithic and into the Neolithic (*14, 112-115*). The cave was also occupied during the Bronze Age and throughout history, and until recently used by shepherds as a stable for sheep.

The main reason why Grotta dell'Uzzo is a key site for Mediterranean prehistory is that its long stratigraphic sequence covers the transition from hunter-gatherer to agro-pastoral economies (*12-14, 59, 60, 115*). It is one of few such sites, given that the number of sites in the Mediterranean with sequences from the late Mesolithic to the early Neolithic is rare, probably as a consequence of a decrease in hunter-gatherer populations at the end of the Mesolithic (*67*).

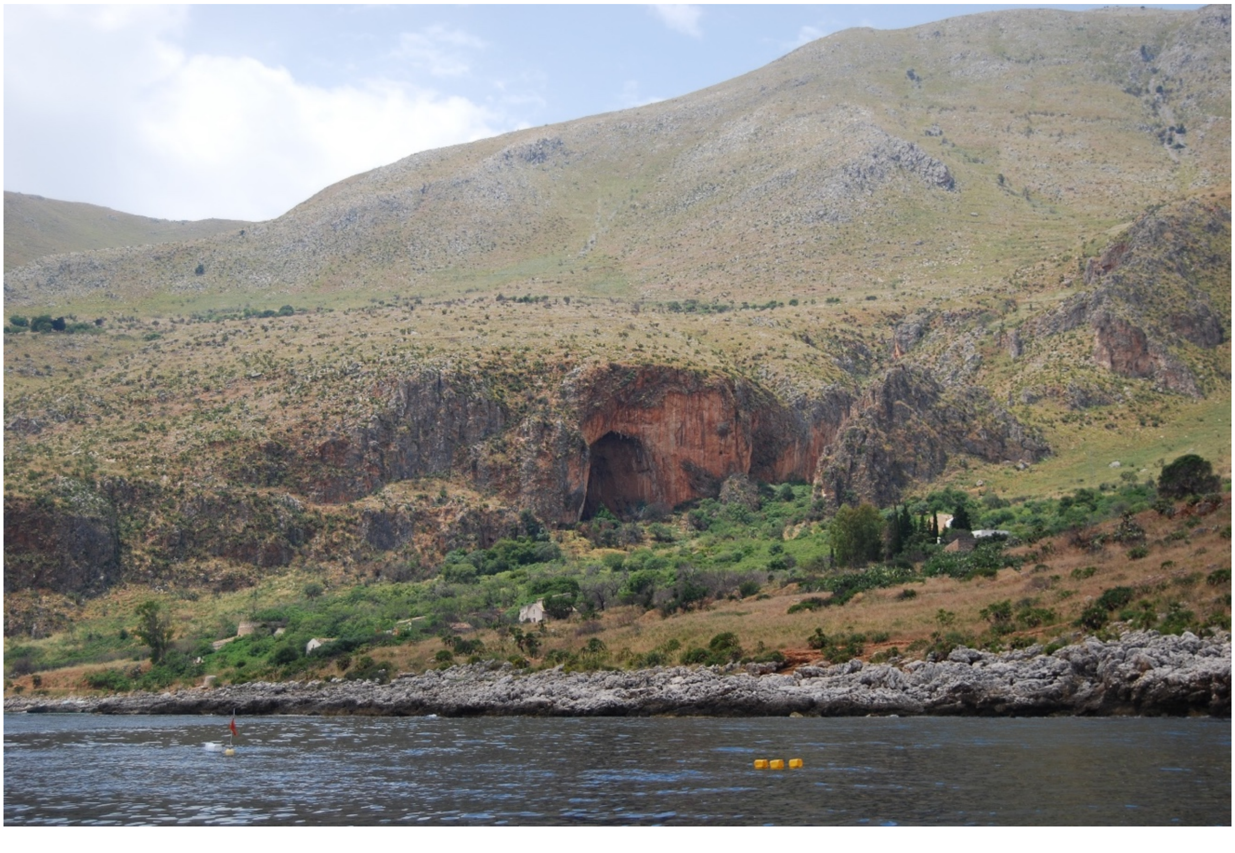

**fig. S1.1. View of Grotta dell'Uzzo from the sea** (photo by Marcello A. Mannino)

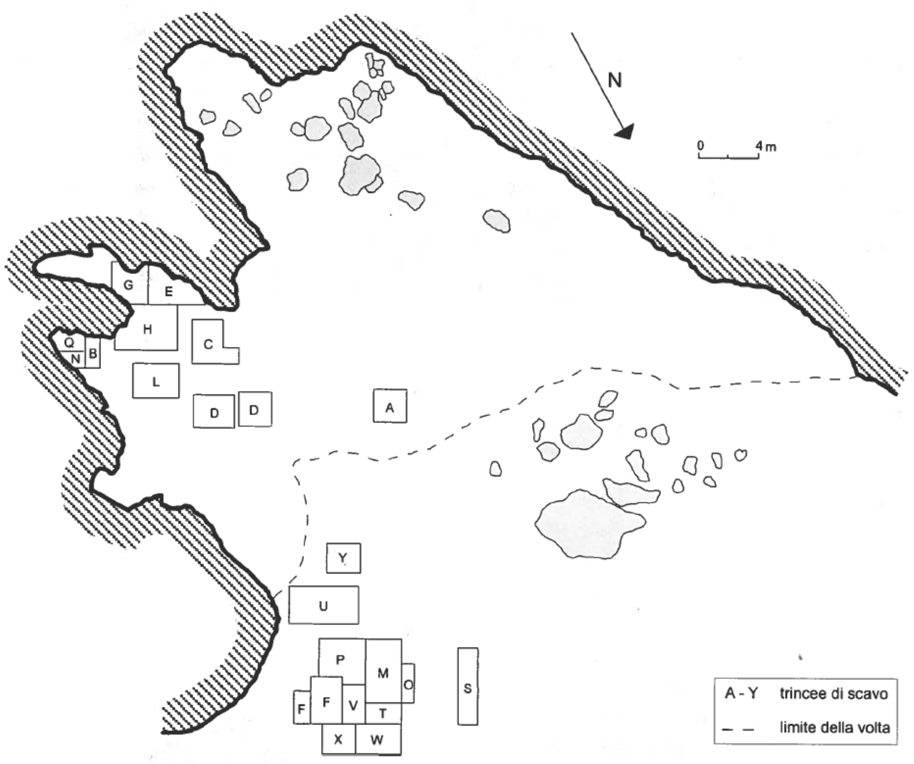


**fig. S1.2. Plan of Grotta dell'Uzzo with the trenches excavated in the 1970s and 1980s** (from: (*14*))

Another important feature of this cave site is that during the Mesolithic it was used as a burial ground. A total of 11 burials and 13 inhumated individuals (six males, four females and three infants) have been recovered at Grotta dell'Uzzo in the course of excavations in the 1970s, 1980s and 2004 close to the walls of the 'inner part' of the cave (*112, 116, 117*). Studies on the pathologies of the inhumated humans established that plant foods were an important component of the diet of the Mesolithic hunter-gatherers (*118*). On the other hand, isotopic and zooarchaeological investigations show that the occupants of Grotta dell'Uzzo relied heavily on animal protein, which through time originated increasingly from marine ecosystems (*12, 14*).

Human remains at the cave were, however, also found scattered through the deposits of Grotta dell'Uzzo. Radiocarbon dating ascertained that these remains were not only Mesolithic but dated to all the main phases of cave occupation, including the so-called Mesolithic-Neolithic transition phase and Neolithic phases (*12*). As part of that same study 70 human bones were sampled, of which 57 recovered from the burials and 13 commingled within the deposits. In total only 33 bones yielded collagen extracts and 10 of these were from the bones recovered outside of the burials. Only 40% of the bones from the burials yielded collagen extracts and not all of these met the quality criteria established by van Klinken (*110*), which is indicative of the poor state of preservation of the human skeletal remains from the burials (*12*). On the other hand, 77% of the commingled bones yielded collagen extracts, all of which are well preserved. For this reason, and because our aim was to obtain genetic and further isotopic information on the main periods of cave occupation, we decided to target the loose human remains, most of which have been directly dated within the remit of this project.

### 2. Cultural succession at Grotta dell'Uzzo from the Mesolithic to Early Neolithic

*Mesolithic*

The lithic industries from the two oldest two phases of the Mesolithic at Grotta dell'Uzzo have not been studied in detail, but they are contemporary to the occurrence in Sicily of facies of Epigravettian tradition across the island and the Undifferentiated Epipalaeolithic in the east, followed by Sauveterrian-like facies (*68*). In north-western Sicily microlithic industries of Epigravettian tradition (labelled as *Epigravettiano indifferenziato*) have been identified at Grotta dell'Uzzo in the Mesolithic phases I and II (*119*), as well as at Grotta dell'Isolidda (*68*) and Grotta di Cala Mancina (*120*) on the western coast of the San Vito lo Capo peninsula. These industries demonstrate strong techno-cultural affinities between the Late Epigravettian and early Holocene hunter-gatherers of Sicily. On the other hand, Sauveterrian industries have not been clearly identified at Grotta dell'Uzzo, but Sauveterrian-like facies have been retrieved from the nearby site of Grotta dell'Isolidda (*68*) and at the site at the westernmost end of Sicily on the island of Favignana at Grotta d'Oriente (*120*).

Levels 14 to 11 in both Trench F and Trench M have previously been defined as the so-called 'Mesolithic-Neolithic transition phase' (e.g. (*12-14, 59, 113, 114, 121, 122*), because this was an essentially Mesolithic phase with some Neolithic elements in its upper spits. A recent study of the lithic industry from these layers attributes this phase of cave occupation to the blade-and-trapeze techno-complex of the western Mediterranean Castelnovian tradition (*69*). The oldest date available for the lowermost spits of this phase obtained on charcoal from spits 14 and 13 of Trench F attributes this part to 7,000-6,590 calBCE ((*58*); P-2734, 7,910 ± 70 BP), which is one of the oldest chronological attributions for a blade-and-trapeze industry (Castelnovian *sensu lato*). The most recent reassessment of the radiocarbon chronology for Grotta dell'Uzzo, based on Bayesian modelling of the sequence of dates available for Trench F, suggests that the phase associated with the Castelnovian facies may have spanned from around 6,770 to 5,850 yrs calBCE (~8,770-7,850 yrs calBP (*12*)). The following phase in chronological continuity is the Neolithic phase I, which according to the above-mentioned Bayesian model may have spanned from around 6,050 to 5,400 yrs calBCE (~8,050-7,400 yrs cal BP; (*12*).

The blade-and-trapeze Castelnovian (*sensu lato*) complex of the VII millennium BCE is in techno-economic continuity with the Neolithic complexes of the archaic Impressed Ware and Stentinello culture of the VI millennium BCE. The production of trapezes constitutes the defining element of the lithic techno-complexes between the VII and VI millennia BCE. This was achieved through a notable standardization of the production processes, particularly through the application of pressure by different modalities (*69*). Nevertheless, the variability in some technical behaviours (e.g. bladelet fracturing techniques, presence/absence of the microburin technique, façonage processes of the trapeze truncations) is linked with a break and discontinuity in the Mesolithic-Neolithic technical traditions (*69*).

*Early Neolithic*

The early Neolithic in Sicily has been defined based on sites in the western part of the island (i.e. Grotta dell'Uzzo, Grotta del Kronio) and is characterized by three main cultural horizons, which in chronological order are: 'Archaic Impressed Ware' (*ceramiche impresse arcaiche*), 'Advanced Impressed Ware' (*ceramiche impresse evolute*) of facies Stentinello I and 'Advanced Impressed Ware' (*ceramiche impresse evolute*) of facies Stentinello I (*8*). The chronology of these horizons is largely based on the dating at Grotta dell'Uzzo, which for this part of the sequence does not see full consensus between the different scholars who worked on the site, depending on whether the beginning of the Neolithic is taken to coincide with Spit 12 of Trench F, as proposed by Tiné and Tusa (*80*), or with Spit 10 of Trench F, as proposed by Tagliacozzo (*14*) and Collina (*69*).

#### 3. Human remains sampled: cultural attribution and chronology

*The interior of the cave*

The first trench excavated at the site was Trench A, which was located within the overhang of the cave. This part of the deposit includes the oldest levels of occupation, which can be attributed to the Late Epigravettian. These are covered by Mesolithic layers, which have been divided into two horizons. The more precise chrono-typological attribution has not been refined due to the partial study of the lithic industries. Horizon 1, the most recent Mesolithic horizon, is characterized by the presence of a specific type of scraper with a reduced front adjacent to a deep laterally-retouched indentation (*123*). This tool is associated to a laminar industry with geometrics, represented by triangles, circular segments and rare trapezes. These characters are not present in Horizon 2, the oldest of the two Mesolithic horizons, in which the geometrics are rarer and replaced by a lithic industry with less differentiated (or more undifferentiated) characters. Below Horizon 2, in a different sedimentary layer, terminal Upper Palaeolithic finds have been recovered. Most trenches within the overhang of the cave (including Trench H) contained deposits of Mesolithic age, given that the Neolithic layers had been removed within it.

We sampled the following individuals from within the overhang of Grotta dell'Uzzo:

- *UZZ26.cont*: cranial fragment retrieved from Spit 8 in Trench A, can be attributed to the phase characterized by microlithic industries of Epigravettian tradition (Mesolithic I phase I). DNA and collagen were extracted from this specimen, but DNA preservation was insufficient to include this individual in our genetic analyses. The direct $^{14}$C date on this individual is 9,436 ± 36 BP (8,810-8,620 calBCE).
- *UZZ61*: phalanx retrieved from the topsoil layer of Trench H, for which we analysed the ancient DNA and collagen. Direct $^{14}$C date: TBA.

  Given the risk of post-depositional disturbance in top soil layers the stratigraphical position for this individual could not be used for a reliable archaeological assignment. However, since this individual has both a genetic profile and isotopic compositon which we consider this individual compatible with the Stentinello culture.

*The deposits outside the cave: Trench F and M*

Most of the human skeletal remains selected for our genetic and isotopic investigation (table S1.1) originate from trenches beyond the overhang of the cave (fig. S1.2). The outside of the cave, and in particular trenches F and M (*69*), contained thick stratigraphic sequences spanning through the Mesolithic and up to the pre-Stentinello (Impressed Ware) and Stentinello Neolithic phases. Trench F contained a Mesolithic deposit of 1,50m in thickness (*113*), overlain by a similarly thick deposit that in chronological order included the so-called 'Mesolithic-Neolithic transition phase' and two Neolithic phases. A recent study of the lithic industries from the transitional layers shows that this phase was associated with the blade-and-trapeze complex attributable to the Castelnovian *sensu lato* (*69*).

The sequence from Trench F is the reference stratigraphy for Grotta dell'Uzzo. The original stratigraphic scheme was proposed by Tagliacozzo (*14*). Based on the findings of Collina (*69*) and on-going investigations, we here classified_the different phases as follows, using the age ranges for each phase as generated by Mannino et al. (*12*):

- Basal stratum / Late Upper Palaeolithic (Spits 48-33): Late Epigravettian
- Mesolithic I, phase I (Spits 32-23): Industries of Epigravettian tradition
- Mesolitihic I, phase II (Spits 22-15): (~11,100-8,500 yrs calBP)
- Mesolithic II (Spits 14-11): Castelnovian facies *sensu lato* (~8,770-7,850 yrs calBP)
- Neolithic phase I (Spits 10-6): Impressed Ware horizon (~8,050-7,400 yrs calBP)
- Neolithic phase II (Spits 5-1): Stentinello horizon (~7,520-7,130 yrs calBP)

Here, we present genomic and isotope data for the following individuals from the F-trench:

- *UZZ33:* tooth retrieved from Spit 4. Although we we could not obtain a direct $^{14}C$ date, this individual was found in the same layer as *UZZ34* that was directly dated.
- *UZZ34*: tooth retrieved from Spit 4, which corresponds to Neolithic I phase II Stentinello. Direct $^{14}C$ date: 6,351±24 BP, 5,470-5,230 calBCE.
  For both *UZZ33* and *UZZ34* the archaeological contextual attribution and ancestry profile are consistent with them being early farmers, most likely from a Stentinello horizon.
- *UZZ40*: tooth of *infans* retrieved from Spit 13, which corresponds to Mesolithic II, Castelnovian *sensu lato.* Direct $^{14}C$ date: 7,471±26 BP, 6,420-6,250 calBCE. This confirms the attribution based on stratigraphic and archaeological observations.
- *UZZ4446*: retrieved from Spit 15, which is stratigraphically assigned to the Mesolithic I phase I. For this individual DNA was extracted from two teeth (skeletal elements UZZ44 and –45) and a mandible fragment (element UZZ46). Collagen was extracted from element UZZ45. Direct $^{14}C$ date: 7,713±26 BP, 6,500-6,250 calBCE. The calibrated age corrected for a marine dietary contribution of 40±10%, attributes this specimen to the Mesolithic II and not to the Mesolithic I phase II. In relation to some cetacean bones, it is possible that materials moved post-depositionally down the sequence from the layer immediately above (Spits 14-11) into Spits 15 and 16 (*12*).
- *UZZ5054*: retrieved from Spits 19 and 20, which correspond to Mesolithic I phase II. DNA was extracted from five different teeth (skeletal elements UZZ50-54). We obtained a direct $^{14}C$ date on element UZZ51: 9,436±29 BP, 8,790-8,630 calBCE. This confirms the attribution based on stratigraphic and archaeological observations.

Trench M had a very similar stratigraphic sequence to Trench F, albeit including only four phases of cave occupation:

- Mesolithic I, phase II (Spits 18-15):
- Mesolithic II (Spits 14-11): Castelnovian facies *sensu lato* (blade-and-trapeze complex)
- Neolithic phase I (Spits 10-7): Impressed Ware culture
- Neolithic phase II (Spits 6-1): Stentinello culture

From the M trench, we present genomic and isotope data for the following individuals:

- *UZZ69*: for who we sampled a mandible retrieved from Spit 3, which corresponds to Neolithic I Stentinello. Direct $^{14}C$ date: 7,848±26 BP, 6,630-6,390 calBCE.
- *UZZ71,* for who we sampled a tooth retrieved from Spit 10, which corresponds to the Neolithic I Impressa. This individual shows an ancestry profile chacteristic for the individuals associated with the Mesolithic II Castelnovian lithic industry. Direct $^{14}C$ date: 7,127±25 BP, 6,060-5,920 calBCE.

The lack of an adequate freshwater isotopic baseline for Grotta dell'Uzzo, and the possibility that this individual may not be local, complicate issues linked to accurate reservoir correction. We have, thus, only calibrated the radiocarbon date for *UZZ71*, but not corrected the calibrated age for possible freshwater effects.

*Trenches S, T, U, W and burial 8*

As discussed above, only few trenches from Grotta dell'Uzzo have been studied and dated more in detail (i.e. trenches A, F and M). For this reason, and because radiocarbon dating has not been applied much on materials from other trenches, the stratigraphically and archaeologically based attribution to cultural phase hinges on observations recorded in the excavation notebooks. However, we have radiocarbon dated almost all the specimens from these poorly-studied trenches, so that their calibrated ages can be related to the stratigraphical and chronological 'master sequence' published for Trench F (*12*).

From Trench S, we present genomic and isotope data for two individuals:

- *UZZ74*: femur retrieved from Spit 5. Direct $^{14}C$ date: 6,310±23 BP, 5,330-5,210 calBCE. The direct $^{14}C$ date, isotopic and ancestry profile are consistent with this individual being an early farmer, most likely from a Stentinello horizon.
- *UZZ75*: petrous bone retrieved from Spit 15. Direct $^{14}C$ date: 6,310±23 BP, 5,330-5,210 calBCE. The direct $^{14}C$ date, isotopic and ancestry profile are consistent with this individual being an early farmer, most likely from a Stentinello horizon.

Although *UZZ74* and *UZZ75* have identical $^{14}C$ dates, we can exclude that these individuals are genetic identicals or have a kinship relation.

From Trench T, we present genomic data for one individual:

- *UZZ77*: tooth, which based on the excavation notebooks and finds recovered from its spit of origin (Spit 13) can be attributed the Neolithic I Impressed Ware horizon. Direct $^{14}C$ date: TBA.

From Trench W, we present genomic and isotope data for two individuals:

- *UZZ87*: humerus retrieved from Spit 2. Direct $^{14}C$ date: 6,286±24 BP, 5,320-5,210 calBCE. The archaeological contextual attribution, direct radiocarbon date, isotopic and ancestry profile are consistent with this individual being an early farmer, most likely from a Stentinello horizon.
- *UZZ88*: phalanx retrieved from Spit 14. Direct $^{14}C$ date: 7,036±25 BP, 6,000-5,840 calBCE. The radiocarbon date is in line with the contextual attribution, both indicating that this individual dates to the Neolithic I Impressed Ware horizon.

From the Mesolithic burial VIII, we present genomic and isotope data for one individual:

- *UZZ96*: molar (M2). This specimen originates from one of the inhumations, which all date to the Mesolithic I (*12*).

  The genetic profile and stratigraphy date are consistent with this individual being a hunter-gatherer from the Early Mesolithic, most likely from the facies Mesolithic I, phase II.

In addition, four individuals were retrieved from the top soil layer in Trench U for which a reliable archaeological assignment could not be ascertained on stratigraphic grounds. However, their direct $^{14}C$ dates, genetic ancestry and isotopic profiles are consistent with them being from the Mesolithic II

Castelnovian facies *sensu lato.* We found genetic evidence for a second-degree kinship relation between *UZZ79* (genetic female) and *UZZ81* (genetic male) (Extended Data Table-4).

- *UZZ79*: petrous bone. Direct $^{14}$C date: 7,809±26 BP, 6,600-6,350 calBCE.
- *UZZ80*: petrous bone. Direct $^{14}$C date: 7,809±26 BP, 6,600-6,350 calBCE.
- *UZZ81*: temporal bone fragment. Direct $^{14}$C date: 7,807±26 BP, 6,600-6,380 calBCE.
- *UZZ82*: petrous bone. Direct $^{14}$C date: 7,729±26 BP, 6,630-6,480 calBCE

| Stratigraphic position | Genetic ID | Cultural phase | Skeletal element |
|---|---|---|---|
| A-8 | UZZ26.cont | Mesolithic I phase I | cranial |
| F-4 | UZZ33 | Early Neolithic Stentinello | tooth |
| F-4 | UZZ34 | Early Neolithic Stentinello | tooth |
| F-13 | UZZ40 | Mesolithic II Castelnovian | tooth (infans) |
| F-15 | UZZ4446 | Mesolithic II Castelnovian | tooth(2x)/mandible |
| F-19/20 | UZZ5054 | Mesolithic I phase II | tooth (5x) |
| H-rim | UZZ61 | Early Neolithic Stentinello | phalanx |
| M-3 | UZZ69 | Mesolithic II Castelnovian | mandible |
| M-7 | UZZ71 | Early Neolithic Impressa? | tooth |
| S-rim | UZZ74 | Early Neolithic Stentinello | femur |
| S-5 | UZZ75 | Early Neolithic Stentinello | temporal/petrous |
| T-13 | UZZ77 | Early Neolithic Impressa? | tooth |
| U-rim | UZZ79 | Mesolithic II Castelnovian | temporal/petrous |
| U-rim | UZZ80 | Mesolithic II Castelnovian | temporal/petrous |
| U-rim | UZZ81 | Mesolithic II Castelnovian | temporal fr |
| U-rim | UZZ82 | Mesolithic II Castelnovian | temporal/petrous |
| W-2 | UZZ87 | Early Neolithic Stentinello | humerus |
| W-14 | UZZ88 | Early Neolithic Impressa? | phalanx |
| burial VIII | UZZ96 | Mesolithic I phase I | M2 upper right |

**table S1.1. Cultural affiliation of the human remains investigated in this study.** This table lists the attribution to cultural phase based on stratigraphic and archaeological grounds. The attribution of samples that have not been clearly assigned to a cultural phase on archaeological grounds is briefly treated in the specimen by specimen list above. In the case of Trench U, although all specimens come from the ‘topsoil layer’, we propose an attribution to the Mesolithic II phase (Castelnovian *sensu lato*) based on a radiocarbon date a delphinid specimen from this part of the deposit ((*12*); 8,083±26 BP: 6,780-6,350 calBCE).

| Individual ID | R-EVA | MAMS | Trench-Spit | Phase | $^{14}$C date BP | Calibrated age BP 2σ | Calibrated age BC 2σ |
|---|---|---|---|---|---|---|---|
| UZZ26.cont | 1918 | 40708 | A-8 | MESO1/1 | 9436±36 | 10760-10570 | 8810-8620 |
| UZZ5054 | 1935 | 40710 | F-19 | MESO1/2 | 9436±29 | 10740-10580 | 8790-8630 |
| UZZ82 | 1960 | 40722 | U-rim | MESO2 | 7809±26 | 8650-8540 | 6690-6590 |
| UZZ69 | 1948 | 40711 | M-3 | MESO2 | 7848±26 | 8580-8340 | 6630-6390 |
| UZZ79 | 1957 | 40719 | U-rim | MESO2 | 7809±26 | 8550-8300 | 6600-6350 |
| UZZ80 | 1958 | 40720 | U-rim | MESO2 | 7809±26 | 8550-8300 | 6600-6350 |
| UZZ81 | 1959 | 40721 | U-rim | MESO2 | 7807±26 | 8550-8330 | 6600-6380 |
| UZZ82 | 1960 | 40722 | U-rim | MESO2 | 7,729±26 | 8650-8540 | 6630-6480 |
| UZZ4446 | 1930 | 40709 | F-15 | MESO2 | 7713±26 | 8450-8200 | 6500-6250 |
| UZZ40 | 2880 | 40726 | F-13 | MESO2 | 7471±26 | 8370-8200 | 6420-6250 |
| UZZ71 | 1950 | 43967 | M-10 | NEO1/1 | 7127±25 | 8010-7870 | 6060-5920 |
| UZZ88 | 1965 | 40712 | W-14 | NEO1/1 | 7036±25 | 7940-7790 | 6000-5840 |
| UZZ34 | 2879 | 40725 | F-4 | NEO1/2 | 6351±24 | 7420-7170 | 5470-5230 |
| UZZ74 | 1953 | 40716 | S-rim | NEO1/2 | 6310±23 | 7280-7160 | 5330-5210 |
| UZZ75 | 1954 | 40717 | S-5 | NEO1/2 | 6310±23 | 7280-7160 | 5330-5210 |
| UZZ87 | 1964 | 40723 | W-2 | NEO1/2 | 6286±24 | 7260-7160 | 5320-5210 |

**table S1.2. Radiocarbon dates, calibrated and corrected ages of humans from Grotta dell'Uzzo**. The AMS radiocarbon dates reported in this table were performed at the Klaus Tschira Laboratory of the Curt-Engelhorn-Zentrum Archaeometrie in Mannheim (MAMS). Dates were calibrated with the OxCal 4.2 software (*124*) using the IntCal13 curve and, in addition, the Marine13 curve for individuals that had consumed marine protein (*125*). The estimation of the amount of marine protein consumed is based on calculations made for specimen S-EVA 8010 (40±10% marine) by (*12*). The individuals for which a correction was necessary are *UZZ4446* (40±10% marine), *UZZ81* (45±10% marine), *UZZ69*, *UZZ79* and *UZZ80* (50±10% marine). Corrections were made using the reservoir correction estimated for the Mediterranean Basin by Reimer and McCormac (*109*), which is $\Delta R = 58\pm85$ $^{14}$C yr. This table is useful to relate the dates from (*12*), which are calibrated BP, with those produced for the present study that are calibrated BCE.

**S2. Genetic grouping and substructure of the ancient Sicilians**

Here, we aimed to investigate genetic substructure among the ancient Sicilians and whether individuals could be grouped for genetic analyses. We co-analyzed an Epigravettian HG from OrienteC (*I2158* (*15*)).

**1. Three genetic groups**

First, we used $f_3$-outgroup statistics of the form $f_3(Mbuti; X1, X2)$ for all individual pairs to quantify their levels of shared genetic drift for SNPs ascertained in Mbuti. We found that the ancient Sicilians in our transect form three major genetic groups that are characterized by elevated levels of shared genetic drift for within-group compared to between-group individual pairs (fig. S2.1). Alternatively, taking an approach presented by Lazaridis et al. (*56*), we used *qpWave (54)* to test for all possible pairs of individuals whether their gene pools are consistent with being derived from one ancestry stream (N = 1) with regard to set of *Outgroups* (table S2.1). If one ancestry stream suffices the model has full rank and the null-hypothesis can not be rejected, hence $f_4(Left1, OG1; OG2, OG3) = f_4(Left2, OG1; OG2, OG3)$ for all triplets. If the latter is true, we assumed that the individuals in the *Left* pair are symmetrically related to the specific combination of *Outgroups* used, and therefore can be grouped for analyses. We found that the qpWave-based cladality models distinguished the same three genetic groups as with the pairwise $f_3$-outgroup statistics.

One group contains the two oldest Mesolithic individuals from Uzzo, *UZZ5054* and *UZZ96* (~8,800-8,630 calBCE) and the Epigravettian *OrienteC* HG (12,250-11,850 calBCE (*15, 17*)). These three individuals carried mitogenome lineages that fall within the U2'3'4'7'8'9 branch (Supplementary Section S7, and (*15, 17*) for *OrienteC*). We labelled this group as **Sicily Early Mesolithic (Sicily EM)**.

A second group contains nine individuals dated to ~6,750-5,850 calBCE. The seven oldest individuals in this group (dated ~6,750-6,250 calBCE) are tentatively assigned to the Mesolithic II Castelnovian archaeological context (Supplementary Section S1). Notably, the two youngest individuals in this group (*UZZ71* and *UZZ88*, dated ~6,050-5,850 calBCE) chronologically coincide with layers at the site that may contain the very first aspects of Impressa Wares (Supplementary Section S1). These two individuals fall fully within the genetic diversity of the other individuals in the Sicily LM group, despite postdating them by ~200 years. The mitogenome haplogroups carried by the all these individuals are typical for European Late Mesolithic WHGs (Supplementary Section S7). We labelled this group as **Sicily Late Mesolithic (Sicily LM)**. Notably, for some individual pairs in this group the qpWave cladality model for one ancestry stream is rejected ($P<0.1$) (table S3.1). This implies that the Sicily LM HGs form a heterogenous group with possibly additional underlying substructure.

The third and most recent genetic group contains seven individuals dated to ~5,460-5,220 calBCE. Six individuals in this group are from layers that chronologically coincide with the presence of Early Neolithic Stentinello Ware, and one individual (*UZZ77*, undated) tentatively with older aspects of Impressa Ware (Supplementary Section S1). All the individuals in the Sicily EN group carried mitogenome haplogroups characteristic for European early farmers (Supplementary Section S7). We labelled this group as **Sicily Early Neolithic (Sicily EN)**.

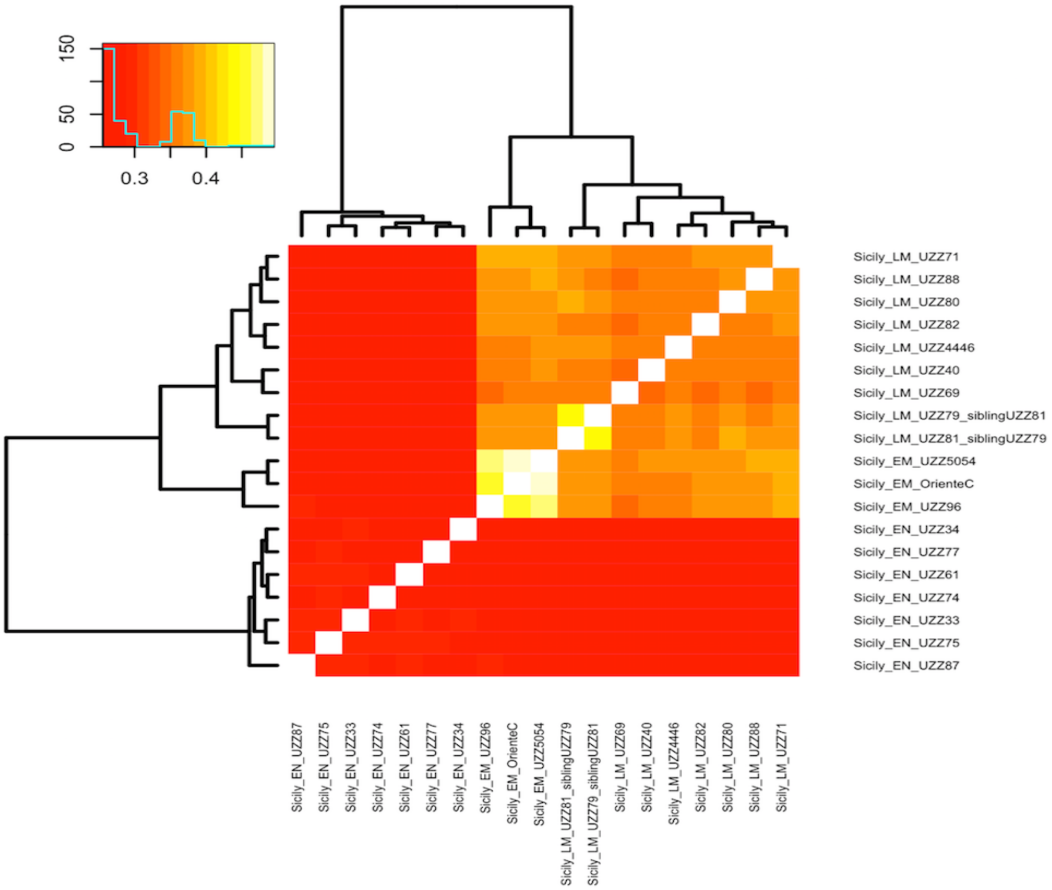


**fig. S2.1. Heat map showing results for *$f_3$(Mbuti; X1, X2)* for all pairwise comparisons between the ancient individuals from Uzzo and one HG from OrienteC.** Larger positive $f_3$-values indicate higher similarity in shared genetic covariance, hence stronger degrees of genetic relatedness between individuals. Three genetic groups appear. Notably, *UZZ5054* and *UZZ96* post-date *OrienteC* by ~3,250-3,450 years yet share very high levels of genetic drift (*$f_3$(Mbuti; OrienteC, UZZ5054)* = 0.495, *$f_3$(Mbuti; OrienteC, UZZ96)* = 0.461, and *$f_3$(Mbuti; UZZ5054, UZZ96)* = 0.465). The higher levels of shared genetic drift between the Sicily LM HGs UZZ79 and UZZ81 are the result of a direct kinship relation (see Extended Data Table-4).

| | OrienteC | UZZ5054 | UZZ096 | UZZ4446 | UZZ40 | UZZ69 | UZZ79 | UZZ80 | UZZ81 | UZZ82 | UZZ71 | UZZ88 | UZZ77 | UZZ33 | UZZ34 | UZZ61 | UZZ74 | UZZ75 | UZZ87 |
|---|---|---|---|---|---|---|---|---|---|---|---|---|---|---|---|---|---|---|---|
| Genetic group | Sicily EM | Sicily EM | | Sicily LM | Sicily LM | Sicily LM | Sicily LM | Sicily LM | Sicily LM | Sicily LM | Sicily LM | Sicily LM | Sicily EN | Sicily EN | Sicily EN | Sicily EN | Sicily EN | Sicily EN | Sicily EN |
| C14 date | | | | | | | | | | | | | | | | | | | |
| # SNPs | | | | | | | | | | | | | | | | | | | |
| **OrienteC** | NR | 0.5660 | 0.9386 | 1.18E-15 | 1.37E-08 | 3.72E-18 | 1.52E-08 | 2.50E-12 | 9.03E-06 | 6.24E-10 | 2.46E-05 | 6.53E-09 | 2.36E-72 | 2.56E-128 | 3.04E-92 | 3.24E-164 | 7.82E-89 | 4.75E-178 | 6.72E-41 |
| **UZZ5054** | 0.5660 | NR | 0.4122 | 1.76E-18 | 2.81E-13 | 4.18E-31 | 3.15E-11 | 5.41E-16 | 1.48E-12 | 4.37E-11 | 9.03E-08 | 1.12E-08 | 7.68E-122 | 5.04E-197 | 2.26E-171 | 2.09E-214 | 8.32E-152 | 6.52E-228 | 3.48E-74 |
| **UZZ096** | 0.9386 | 0.4122 | NR | 0.0201 | 0.5583 | 1.37E-04 | 0.6865 | 0.0281 | 0.3366 | 3.48E-03 | 0.8156 | 0.7681 | 2.13E-22 | 6.99E-44 | 2.03E-26 | 1.09E-51 | 2.72E-32 | 1.74E-54 | 5.92E-10 |
| **UZZ4446** | 1.18E-15 | 1.76E-18 | 0.0201 | NR | 0.8770 | 0.0333 | 0.1123 | 0.1407 | 0.6736 | 0.0803 | 0.0812 | 0.0555 | 6.40E-56 | 1.19E-112 | 1.21E-95 | 2.46E-122 | 1.18E-71 | 8.30E-124 | 3.27E-33 |
| **UZZ40** | 1.37E-08 | 2.81E-13 | 0.5583 | 0.8770 | NR | 8.67E-03 | 0.0589 | 0.4519 | 0.8869 | 0.1467 | 0.0110 | 0.4252 | 9.03E-40 | 6.03E-84 | 1.62E-71 | 1.50E-104 | 9.15E-38 | 8.24E-99 | 1.75E-17 |
| **UZZ69** | 3.72E-18 | 4.18E-31 | 1.37E-04 | 0.0333 | 8.67E-03 | NR | 1.48E-04 | 2.41E-03 | 0.1819 | 3.41E-04 | 8.38E-09 | 1.01E-07 | 1.16E-67 | 1.53E-109 | 7.02E-81 | 4.33E-126 | 6.50E-85 | 1.81E-126 | 5.80E-35 |
| **UZZ79** | 1.52E-08 | 3.15E-11 | 0.6865 | 0.1123 | 0.0589 | 1.48E-04 | NR | 0.4076 | 0.9433 | 0.6787 | 0.8609 | 0.8865 | 8.34E-78 | 1.24E-137 | 9.21E-133 | 1.36E-162 | 5.14E-103 | 3.77E-162 | 5.80E-50 |
| **UZZ80** | 2.50E-12 | 5.41E-16 | 0.0281 | 0.1407 | 0.4519 | 2.41E-03 | 0.4076 | NR | 0.7294 | 0.9650 | 0.0705 | 0.4134 | 9.98E-90 | 1.17E-148 | 1.18E-123 | 1.13E-163 | 2.11E-94 | 4.68E-170 | 3.38E-50 |
| **UZZ81** | 9.03E-06 | 1.48E-12 | 0.3366 | 0.6736 | 0.8869 | 0.1819 | 0.9433 | 0.7294 | NR | 0.8256 | 0.6475 | 0.7192 | 7.99E-51 | 3.96E-93 | 9.51E-76 | 2.18E-123 | 1.05E-70 | 6.61E-118 | 3.40E-28 |
| **UZZ82** | 6.24E-10 | 4.37E-11 | 3.48E-03 | 0.0803 | 0.1467 | 3.41E-04 | 0.6787 | 0.9650 | 0.8256 | NR | 0.0958 | 0.6946 | 7.74E-82 | 1.85E-157 | 1.71E-126 | 2.81E-161 | 4.50E-106 | 8.55E-165 | 7.03E-44 |
| **UZZ71** | 2.46E-05 | 9.03E-08 | 0.8156 | 0.0812 | 0.0110 | 8.38E-09 | 0.8609 | 0.0705 | 0.6475 | 0.0958 | NR | 0.9360 | 1.11E-94 | 4.68E-146 | 2.09E-115 | 1.54E-157 | 3.37E-108 | 3.20E-186 | 2.41E-46 |
| **UZZ88** | 6.53E-09 | 1.12E-08 | 0.7681 | 0.0555 | 0.4252 | 1.01E-07 | 0.8865 | 0.4134 | 0.7192 | 0.6946 | 0.9360 | NR | 9.92E-76 | 8.46E-143 | 7.05E-125 | 2.57E-160 | 5.66E-96 | 1.24E-165 | 3.01E-41 |
| **UZZ77** | 2.36E-72 | 7.68E-122 | 2.13E-22 | 6.40E-56 | 9.03E-40 | 1.16E-67 | 8.34E-78 | 9.98E-90 | 7.99E-51 | 7.74E-82 | 1.11E-94 | 9.92E-76 | NR | 0.6138 | 0.0096 | 0.7563 | 0.9473 | 0.3566 | 0.7585 |
| **UZZ33** | 2.56E-128 | 5.04E-197 | 6.99E-44 | 1.19E-112 | 6.03E-84 | 1.53E-109 | 1.24E-137 | 1.17E-148 | 3.96E-93 | 1.85E-157 | 4.68E-146 | 8.46E-143 | 0.6138 | NR | 0.2600 | 0.1701 | 0.6806 | 0.5095 | 0.4760 |
| **UZZ34** | 3.04E-92 | 2.26E-171 | 2.03E-26 | 1.21E-95 | 1.62E-71 | 7.02E-81 | 9.21E-133 | 1.18E-123 | 9.51E-76 | 1.71E-126 | 2.09E-115 | 7.05E-125 | 0.0096 | 0.2600 | NR | 0.6946 | 0.2384 | 0.5425 | 0.4452 |
| **UZZ61** | 3.24E-164 | 2.09E-214 | 1.09E-51 | 2.46E-122 | 1.50E-104 | 4.33E-126 | 1.36E-162 | 1.13E-163 | 2.18E-123 | 2.81E-161 | 1.54E-157 | 2.57E-160 | 0.7563 | 0.1701 | 0.6946 | NR | 0.6973 | 0.2914 | 0.9762 |
| **UZZ74** | 7.82E-89 | 8.32E-152 | 2.72E-32 | 1.18E-71 | 9.15E-38 | 6.50E-85 | 5.14E-103 | 2.11E-94 | 1.05E-70 | 4.50E-106 | 3.37E-108 | 5.66E-96 | 0.9473 | 0.6806 | 0.2384 | 0.6973 | NR | 0.8842 | 0.0837 |
| **UZZ75** | 4.75E-178 | 6.52E-228 | 1.74E-54 | 8.30E-124 | 8.24E-99 | 1.81E-126 | 3.77E-162 | 4.68E-170 | 6.61E-118 | 8.55E-165 | 3.20E-186 | 1.24E-165 | 0.3566 | 0.5095 | 0.5425 | 0.2914 | 0.8842 | NR | 0.0263 |
| **UZZ87** | 6.72E-41 | 3.48E-74 | 5.92E-10 | 3.27E-33 | 1.75E-17 | 5.80E-35 | 5.80E-50 | 3.38E-50 | 3.40E-28 | 7.03E-44 | 2.41E-46 | 3.01E-41 | 0.7585 | 0.4760 | 0.4452 | 0.9762 | 0.0837 | 0.0263 | NR |

**table S2.1. Ancestry similarity matrix for all individual pairs.** Results are from qpWave-based cladality models. We used an *Outgroup* set from Mathieson et al. (*17*): *El Miron*, *Mota*, Mbuti, *Ust Ishim*, *Mal'ta*, *AfontovaGora3*, *GoyetQ116*, *Villabruna*, *Kostenki14*, *Vestonice16*, Karitiana, Papuan, Onge. Individuals in red have < 150k SNPs covered. Since missing data may inflate the P-values for this test, we required a test result to be smaller (less extreme) than P = 0.1 in order to reject the null-hypothesis of cladality. Models that provide a full ancestry fit (significance threshold: P ≥ 0.1) are highlighted in green, those that approach the boundaries of the model (0.01 < P < 0.1) are in orange, and those for which a full ancestry fit can be rejected (P < 0.01) are in red. We found three genetic groups (boxes) that we labeled as Sicily EM (*OrienteC*, *UZZ5054*), Sicily LM (*UZZ4446*, *UZZ40*, *UZZ69*, *UZZ79*, *UZZ80*, *UZZ81*, *UZZ82*, *UZZ71*, *UZZ88*) and Sicily EN (*UZZ77*, *UZZ33*, *UZZ34*, *UZZ61*, *UZZ74*, *UZZ75*, *UZZ87*). The Sicily LM HGs form a heterogenous group with possible additional underlying substructure. *UZZ96* shows a high smilarity to individuals in both the Sicily EM and Sicily LM genetic group, congruent to its position in the MDS plot (Fig. 2A).

### S3. Elevated lineage-specific genetic drift in the Sicilian Early Mesolithic HGs

#### Nucleotide diversity (π)

We selected a total of 120 West-Eurasian HGs with >150k SNPs covered on the 1240k panel, of which 103 were previously published (Extended Data Table 1), from four broad geographical regions that we labeled as "western" (n=18), "south-western" (n=7), "southern-central" (n=33), and "(south)-eastern" (n=62) Europe. We subgrouped the individuals further based on similar $^{14}$C-dating and genetic cluster assignment (*16, 17, 36, 38, 50*) (for an overview of the HG groups, see Extended Data Table 1. E.g. we made seperate groups for individuals associated with the Villabruna cluster, and those high in Magdalenian-related ancestry. We determined the nucleotide diversity (π) from pseudo-haploid genotypes by calculating the average proportion of nucleotide mismatches for overlapping autosomal SNPs covered by at least one read by both individuals in a pair within a given group. Since individual pairs and not chromosome pairs are considered, this measure of nucleotide diversity does not include the global heterozygosity levels of individuals (e.g. (*32, 34, 48*)). We restricted to the set of ~870k CpG-filtered autosomal SNPs and removed individual pairs that shared less than 35,000 SNPs covered. Standard errors were determined from block jackknifes over 5Mb windows and 95% confidence intervals (95CIs) from 1,000 bootstraps. Then we calculated an average over all the individual pairs within a HG group.

We find a significantly lower nucleotide diversity (π) for individuals from the Early Mesolithic time period (π = 0.165, 95CI = 0.161-0.170), compared to those from the preceding Upper Paleolithic (π = 0.233, 95CI = 0.227-0.239), and subsequent Late Mesolithic (π = 0.220, 95CI = 0.217-0.223), Early Neolithic (π = 0.252, 95CI = 0.248-0.256), and later time periods (fig. S3.1).

In addition, the nucleotide diversity for the Sicily EM HGs is ~20% lower compared to contemporaneous Villabruna-cluster related individuals from Central Europe (π = 0.217, 95CI = 0.211-0.222), Magdalenian individuals from Iberia (π = 0.221, 95CI = 0.216-0.226) and the earliest Iron Gates HGs in Serbia (π = 0.226, 95CI = 0.222-0.229) (Fig. 3). Notably, we also find a reduction in genetic diversity for Upper Paleolithic HGs from Central Europe with Magdalenian-associated ancestry and in related Early Mesolithic HGs that are part of the ElMiron genetic cluster (*16*) ("western Europe UP Magdalenian + EM (15.5-12.7 kya)": π = 0.174, 95CI = 0.167-0.181). Intriguingly, this reduction is not found for the closely related Magdalenian- associated ElMiron genetic cluster HGs from Iberia, nor in contemporaneous individuals from Central Europe that are part of the Villabruna genetic cluster (*16*). These results underline previous suggestions for a possible genetic bottleneck in Central European Magdalenian individuals (*16, 52*).

To check whether our results are driven by differences in ascertainment bias, false positives due to sequencing errors or ancient DNA damage, we repeated the analysis for 94,469 autosomal SNP ascertained in Yoruba, an African outgroup, vis-à-vis (*17, 48*). We find that while the absolute values for the nucleotide diversity changes, the overall trend mirrors that of the full data set (fig. S3.2).

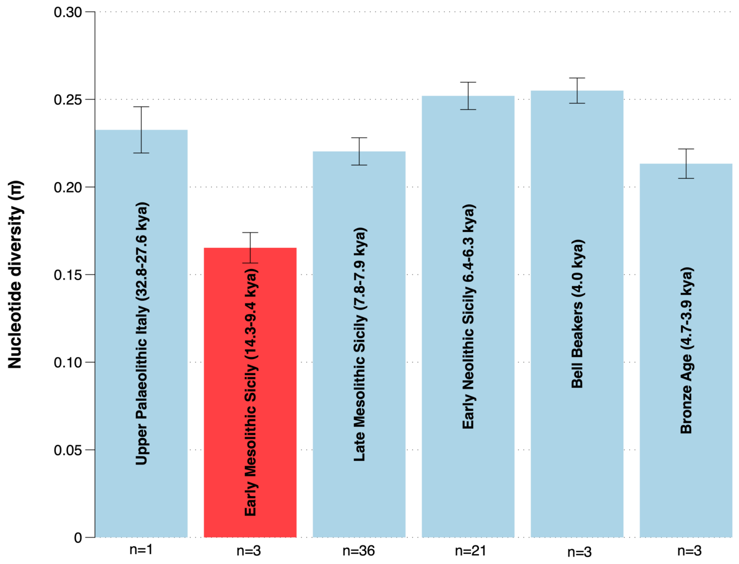


**fig. S3.1. Changes in the nucleotide diversity over time for individuals from peninsular Italy and Sicily**. The nucleotide diversity (π) is plotted for various transect groups in archaeo-chronological order. Upper Palaeolithic (Paglicci13, Ostuni1), Early Mesolithic (Sicily EM HGs), Late Mesolithic (Sicily LM HGs), Early Neolithic (Sicily EN farmers), Bell Beakers (Italy Bell Beakers), and Bronze Age (Italy Remedello), see Extended Data Table 1 for details on the grouping. The number of tests (n) that is used to determine the average for each time period is given. Error bars reflect 3 SEs.

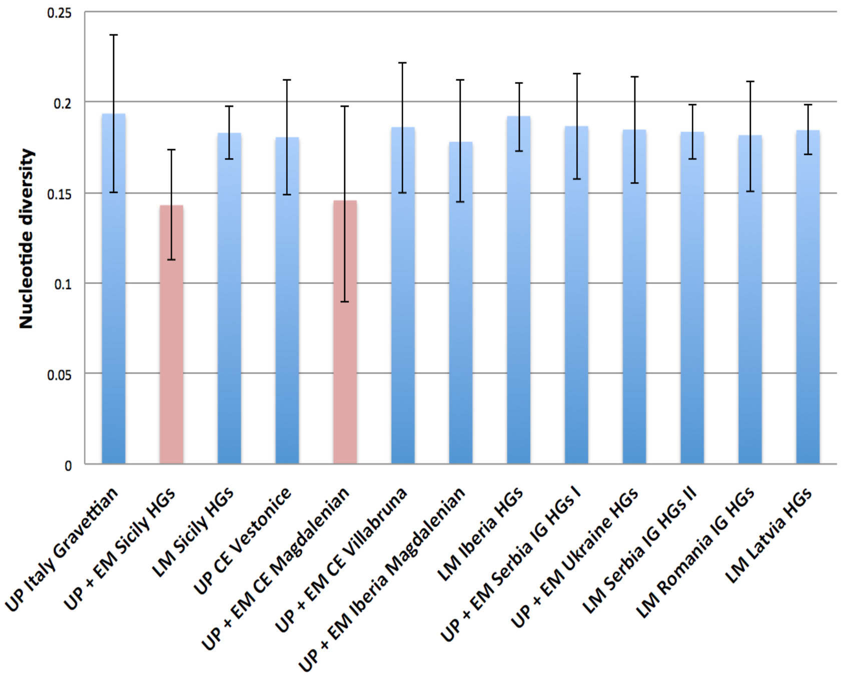


**fig. S3.2. Nucleotide diversity (π) for the various Eurasian HG groups for autosomal SNP sites ascertained in Yoruba**. Error bars reflect 3 SEs.

**Individual Heterozygosity (H)**

Secondly, we investigated whether the genetic diversity within the individual Sicily EM HGs is reduced compared to the later Sicily HGs and early farmers. We hence quantified the **individual heterozygosity (H)**, the proportion of sites considered heterozygous among all sites analysed. We calculated heterozygosity as the sum of heterozygous genotypes estimated using *SnpAD* (*126*) (version 0.3.3 with parameters --max_gtfreq=0.2) (table S3.1). Error profiles were calculated separately for single-stranded and double-stranded libraries, when both types of data were available. Confidence intervals (95%) (95CIs) were calculated using *snpADci*, which determines multiple testing corrected confidence intervals around heterozygous genotype frequencies. These confidence intervals were summed to arrive at confidence intervals for the heterozygosity.

As a quality check we investigated whether differences in calling rate for the alternative allele influenced the calculated heterozygous genotype frequencies. The calling rate may be biased when a heterozygous SNP site is covered by only a few reads. When the SNP depth is low the alternative allele may not be observed. Indeed, when the heterozygosity level (H) is plotted as a function of the read depth (X), individuals with an average SNP depth of 0.08-1.54X have a considerably lower calling rate for the alternative allele at heterozygous sites (fig. S3.3). This bias plateaus in our dataset in individuals with > 1.94X read coverage, and we hence used this as a cutoff for our analysis.

Subsequently, with the *boxplot.stats()* function in R we found that the individual heterozygosity for *UZZ88* is reduced compared to other Sicily LM HGs and does not fall within the variance of this group. We hence did not include this individual in the Sicily LM group average. We found that the average individual heterozygosity (H) for Sicily EM HGs is 30% lower compared to Sicily LM HGs (non-overlapping 95CIs), and 40% compared to Sicilian EN farmers (non-overlapping 95CIs) (fig. S3.4).

| Individual label | # autosomal SNPs covered on 1240k | Average read depth for SNP (X coverage) | Individual Heterozygosity (H) | 95% CI low. bound | 95% CI up. bound |
|---|---|---|---|---|---|
| Sicily EM UZZ05054 | 502,957 | 3.5990 | 0.1422 | 0.1391 | 0.1451 |
| Sicily EM OrienteC | 155,489 | NA | 0.1628 | 0.1519 | 0.1722 |
| Sicily EM UZZ096 | 48,824 | 0.1038 | 0.1042 | 0.0895 | 0.1218 |
| Sicily LM UZZ069 | 449,167 | 2.7736 | 0.2086 | 0.2046 | 0.2119 |
| Sicily LM UZZ079 | 534,685 | 7.1628 | 0.2165 | 0.2135 | 0.2197 |
| Sicily LM UZZ080 | 561,466 | 9.1715 | 0.2139 | 0.2107 | 0.2166 |
| Sicily LM UZZ082 | 517,375 | 4.7542 | 0.2132 | 0.2109 | 0.2179 |
| Sicily LM UZZ040 | 175,573 | 0.5139 | 0.1206 | 0.1139 | 0.1267 |
| Sicily LM UZZ04446 | 356,411 | 1.1895 | 0.2242 | 0.2193 | 0.2294 |
| Sicily LM UZZ071 | 373,717 | 1.5397 | 0.1739 | 0.1703 | 0.1792 |
| Sicily LM UZZ081 | 213,244 | 0.6539 | 0.1628 | 0.1567 | 0.1702 |
| Sicily LM UZZ088 | 406,054 | 1.9261 | 0.1999 | 0.1963 | 0.2045 |
| Sicily EN UZZ061 | 472,518 | 3.5955 | 0.2481 | 0.2438 | 0.2525 |
| Sicily EN UZZ075 | 539,032 | 5.8031 | 0.2484 | 0.2458 | 0.2521 |
| Sicily EN UZZ033 | 248,381 | 0.7065 | 0.2126 | 0.2061 | 0.2192 |
| Sicily EN UZZ034 | 182,339 | 0.5185 | 0.1879 | 0.1802 | 0.1954 |
| Sicily EN UZZ074 | 132,318 | 0.3468 | 0.0744 | 0.0667 | 0.0840 |
| Sicily EN UZZ077 | 116,020 | 0.2542 | 0.2129 | 0.2022 | 0.2273 |
| Sicily EN UZZ087 | 39,706 | 0.0806 | 0.1421 | 0.1209 | 0.1657 |

**table S3.1. Individual heterozygosity (H) levels for the ancient Sicilians in our transect.** 95% confidence intervals (95CI) were determined using block jackknifes over 5Mb windows and corrected for multiple testing. The total number of autosomal SNPs covered on the 1240k panel and average read depth for the SNPs are given. Individuals that were excluded from analysis are in red.

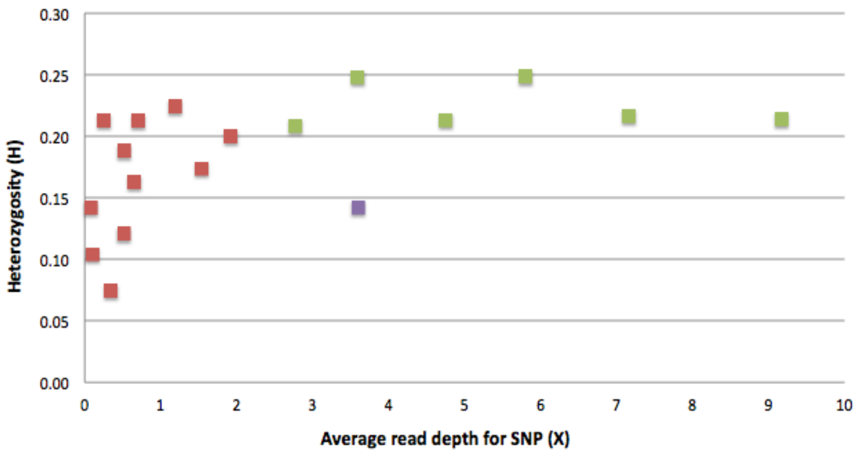


**fig. S3.3. Plot for the individual Heterozygosity (H) levels in our ancient Sicilians as a function of their average SNP depth.** Red: low coverage individuals that were excluded from analysis. For these individuals we found a systematic bias in the calling rate for the alternative allele at heterozygous sites. Green: individuals > 1.94X coverage that passed our quality threshold filters. Purple: Sicily EM HG *UZZ5054*. The observed lower heterozygosity resulted from its population genetic history. *OrienteC* is not plotted (H = 0.163, average SNP depth unknown).

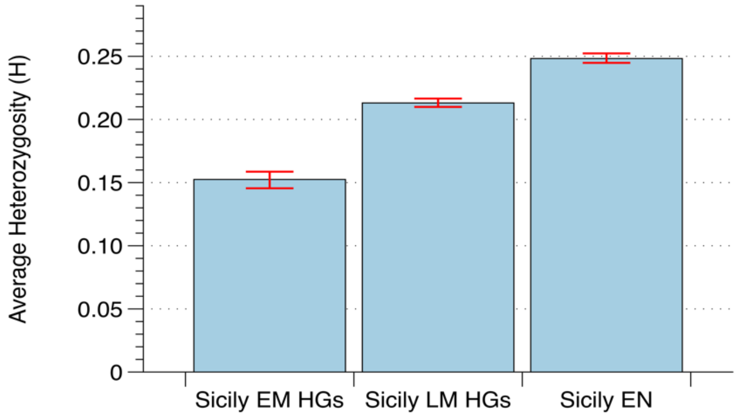


**fig. S3.4. Barplot showing the average individual heterozygosity (H) for Sicily EM HGs, LM HGs and early farmers (EN).** The 95% confidence intervals are given in red. The average individual heterozygosity (H) for Sicily EM HGs is 0.153 (95CI: 0.146-0.157), for Sicily LM HGs 0.213 (95CI: 0.210-0.217) and for Sicily early farmers (EN) 0.248 (95CI: 0.245-0.252). The confidence interval for Sicily EM HGs does not overlap with that for either Sicily LM HGs or Sicily EN.

### S4. Characterizing the Sicilian Mesolithic HGs ancestry using F-statistics

First, we used outgroup $f_3$-statistics to investigate for various West-Eurasian HGs (*X*) which one is genetically closest to Sicily EM HGs and Sicily LM HGs, using $f_3$*(Mbuti; Sicily EM HGs, X)* and $f_3$*(Mbuti; Sicily LM HGs, X),* respectively (fig. S4.1). The highest amount of shared genetic drift for Sicily EM HGs is with Sicily EM HG *UZZ96*, followed by Villabruna cluster individuals and Sicily LM HGs. Sicily LM HGs show the highest degree of allele sharing with Sicily EM HGs, followed by other individuals from the Villabruna cluster.

Secondly, we performed $f_4$-cladality statistics of the form $f_4$*(Chimp, Sicily LM HGs; Sicily EM HGs, X)* and $f_4$*(Chimp, Sicily EM HGs; Sicily LM HGs, X)* (fig. S4.2). For almost all tested HGs *X* the statistic is $\leq 0$, implying that Sicily LM and EM HGs form a clade to the exclusion of other West-Eurasian HGs. This suggests that the shared genetic drift level measured in the above $f_3$-outgroup statistics most likely reflects a direct ancestry connection between Sicily EM HGs and Sicily LM HGs. However, Sicily EM HGs do not represent all the ancestry in the Sicily LM HGs, since in the $f_4$- statistic $f_4$*(Chimp, X; Sicily EM HGs, Sicily LM HGs)* additional admixture signals are found for various HGs from (south)-eastern Europe and Russia (Fig. 4A).

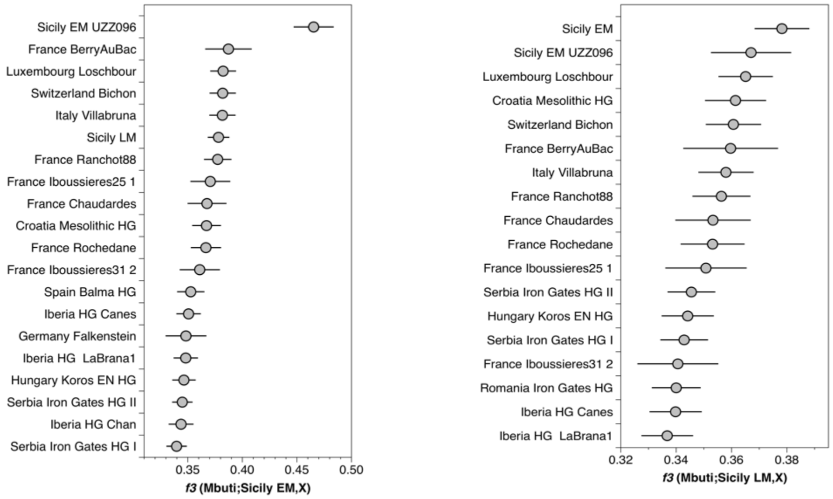


**fig. S4.1. $F_3$-outgroup statistics for the Mesolithic Sicilian HGs**. Error bars reflect 3 SEs.

.

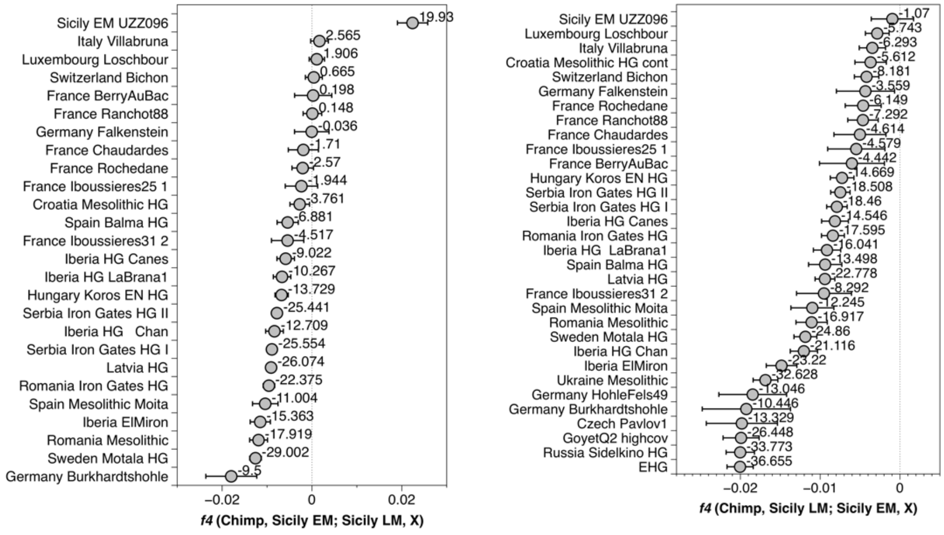


**fig. S4.2**. **$F_4$-cladality statistics for the Mesolithic Sicilian HGs.** Error bars reflect 3 SEs, z-values are given.

Subsequently, we aimed to compare the ancestry as found in *Villabruna* with that in the Mesolithic Sicilian HGs and their respective affinities to various West Eurasian HGs (*X*). Accordingly, we perfomed $f_4$-cladality tests of the form *$f_4$(Chimp, X; Sicily EM HGs, Villabruna),* and *$f_4$(Chimp, X; Sicily LM HGs, Villabruna)*. Notably, comparing the ancestry of Sicily EM HGs and *Villabruna* results in a similar geographical separation in their genetic affinitties to western and eastern European HGs as found for *$f_4$(Chimp, X; Sicily EM HGs, Sicily LM HGs)* (Fig. 4). Also here, Sicily EM HGs share an excess of alleles with western European HGs, including the majority of Villabruna cluster individuals, whereas *Villabruna* does with (south-)eastern European HGs (fig. S4.3 - Left). This suggests that *Villabruna* and Sicily LM HGs behave genetically similar in relation to Sicily EM HGs.

If the gene pool of the Sicily LM HGs is very similar to that of *Villabruna*, we do not expect any HGs from Eurasia to significantly share more alleles with either of them. In an $f_4$-cladality test, we hence expect *$f_4$(Chimp, X; Villabruna, Sicily LM HGs)* $\approx 0$. However, EHG is marginally closer to Sicily LM HGs, whereas, Sicily EM HGs and some other West-Eurasian HGs from the Villabruna and Gravettian cluster are closer to *Villabruna* (fig. S4.3 - Right).

Taken together, these *F*-statistics suggest that *Villabruna*, Sicily EM HGs and Sicily LM HGs share significant ancestry but differ in their affinities towards eachother and to HGs from south(-eastern) Europe and HGs with Magdalenian-related ancestry from southwestern Europe.

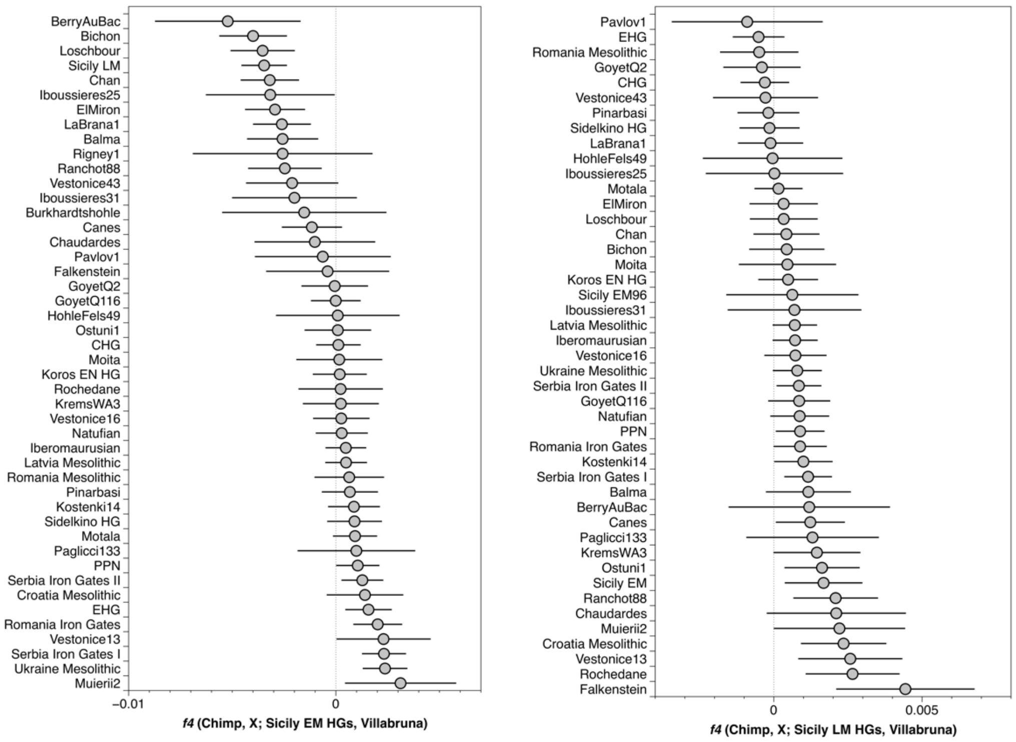


**fig. S4.3. Cladality $f_4$-statistics to compare the ancestry in Villabruna to that of the Sicilian Mesolthic HGs with *$f_4$(Chimp, X; Sicily EM/LM HGs, Villabruna).*** Error bars reflect 2 SEs.

### S5. Investigating the phyologenetic position of the Early Mesolithic Sicilian HGs

Admixture graph models fit allele frequency correlations and allow us to hierarchically build an increasingly complex framework of ancestry streams that fit the genetic diversity observed. Here, we used the *qpGraph* program (*57*) to construct a phylogeny of ancestry lineages found among Palaeolithic and Mesolithic West-Eurasian HGs to further clarify the phylogenetic position of Sicily EM HGs in relation to *Villabruna*, other Early Mesolithic HGs from continental Europe (EM WHGs), and Magdalenian-associated HGs (e.g. *El Miron* and *GoyetQ2*).

We built the phylogeny models with increasing complexity by fitting representative West Eurasian HG ancestry lineages one by one to the phylogeny roughly in order of their respective $^{14}$C dates. We added each of them to all possible nodes as a branch without admixture or as a binary admixture between two branches. We selected models that did not include trifurcations or 0% ancestry stream estimates, and for which the difference between the observed and fitted *f*-statistics were the lowest (the maximum deviation falls within 3.5 SEs for our preferred models). We preferred a model that fits a HG lineage as a branch without admixture over one with additional admixture if both of them fit the observed *f*-statistics equally well.

For this analysis, we grouped individuals that have similar ancestry:

| | |
|---|---|
| **Sicily EM** | *I2158/OrienteC* (~14 kyBP), *UZZ05054* (~10.5 kyBP) |
| **EM WHGs** | Early Mesolithic WHGs:<br>*Bichon* (~13.5 kyBP), *Rochedane* (~13 kyBP), *Iboussieres25* (~12 kyBP), *Iboussieres31* (~11.5 kyBP) |

We started with a core model (CM-5) phylogeny fitting five populations that separates African (Mbuti) from non-African ancestry (*UstIshim* ~45 kyBP (*27*)), followed by a major split between basal West Eurasian (*Kostenki14*, ~36 kyBP (*127*)) and Ancient North Eurasian (ANE) ancestry (*Mal'ta*, ~24 kyBP (*36, 46*)) Subsequently, we added ~30 kyBP *Vestonice16* (*16*) associated with the Gravettian.

CM-5: 1) Mbuti, 2) *Ust Ishim*, 3) *Kostenki14*, 4) *Mal'ta*, 5) *Vestonice16*

We found two models that fit the data (fig. S5.1). In the least complex model *Vestonice16* is fitted as branch without admixture as a sister lineage to *Kostenki14*, and *Mal'ta* as an outgroup to both of them. In the alternative model, *Vestonice16* is on an admixed branch with a source related to *Mal'ta* contributing 92% and *Kostenki14* contributing 8%.

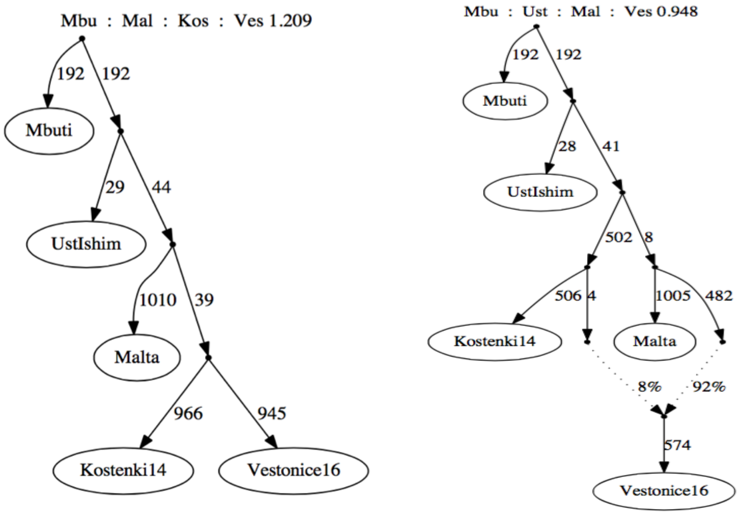


**fig. S5.1. CM-5 models fitting 5 populations.** Left: *Vestonice16* is fitted as unadmixed on a branch with *Kostenki14* with *Mal'ta* as an outgroup to both of them (1 outlier, max $|f_4$, expected – $f_4$, observed$|$ = 1.209). Right: *Vestonice16* is admixed (1 outlier, max $|f_4$, expected – $f_4$, observed$|$ = 0.948).

To the least complex model we then added either *GoyetQ116* (~35 kyBP Aurignacian (*16*)), *El Miron* (~19 kyBP Magdalenian (*16*)) or *GoyetQ2* (~15 kyBP Magdalenian (*16, 50*) as a representative of an ancestry lineage characteristic for Magdalenian-associated HGs (CM-6). Subsequently, we added ~18 kyBP *AfontovaGora3* (*16*) (CM-7), a more recent representative of the ANE ancestry lineage.

CM-6 = CM-5 + *GoyetQ116* or *El Miron* or *GoyetQ2*
CM-7 = CM-6 + *AfontovaGora3*

*GoyetQ116* and *El Miron* can be fitted as branches without admixture in both core models (fig. S5.2 - Left & Center, only the least complex models are shown). In contrast *GoyetQ2* is best modeled as a mixture between 47% *Vestonice16* ancestry and 53% ancestry from a basal lineage that branches of basal to HGs with ANE ancestry. In all three models, *AfontovaGora3* is best fitted on a branch with *Mal'ta* without additional admixture (fig. S5.2 - Right).

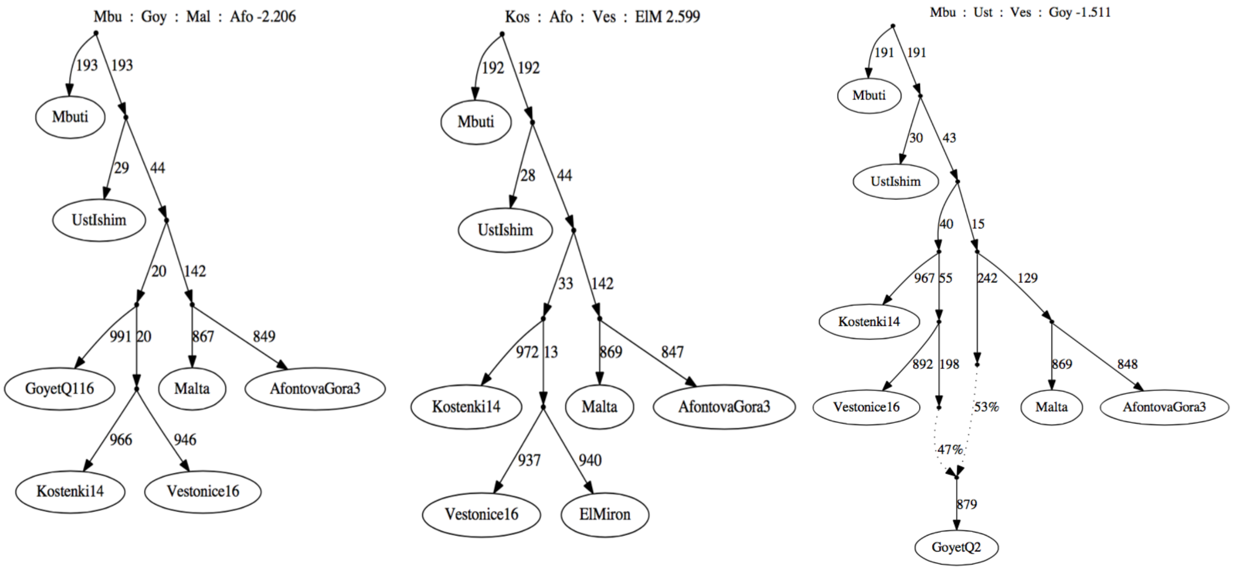


**fig. S5.2. CM-7 models fitting different HGs as representatives of an ancestry found in Magdalenian-associated individuals.** Left: *GoyetQ116* as a branch without admixture (1 outlier, |$f_4$, expected – $f_4$, observed| = 2.206). Center: *El Miron* as a branch without admixture (1 outlier, |$f_4$, expected – $f_4$, observed| = 2.599). Right: *GoyetQ2* as a mixture between *Vestonice16* and a lineage basal to ANE (1 outlier: |$f_4$, expected – $f_4$, observed| = 1.511).

We then made an experimental model series to investigate how *Villabruna* and Sicily EM HGs relate to each other, and whether either of them requires an additional ancestry contribution from a Magdalenian-associated or ANE-related source. We added ~14 kyBP *Villabruna* (*16*) followed by the Sicily EM HGs, and *visa versa*, either as a branch without or with admixture to the seven population core models (EM-9a+b).

EM-9a: CM-7 + *Villabruna* + Sicily EM HGs
EM-9b: CM-7 + Sicily EM HGs + *Villabruna*

The least complex model that fits the gene pools of *Villabruna* and Sicily EM HGs is the same regardless of the order in which we added them (fig. S5.3 - Left): Sicily EM HGs and *Villabruna* form a clade, with *GoyetQ2* as an outgroup to both of them. Notably, models that include one admixture event fit the gene pools of both Sicily EM HGs and *Villabruna* approximately equally well. However, the branches that contributed ancestry are different for the two. When *Villabruna* is added to the graph first (EM-9a), Sicily EM HGs is fitted on an admixed branch that derives 37% ancestry from a *Villabruna*- and 63% from a *GoyetQ2*-related lineage (fig. S5.3 - Center). In contrast, when Sicily EM HGs is added first (EM-9b), *Villabruna* is fitted on an admixed branch that derives 97% ancestry from a lineage close to Sicily EM HGs and 3% ancestry from a lineage related to *AfontovaGora3* (fig. S5.3 - Right).

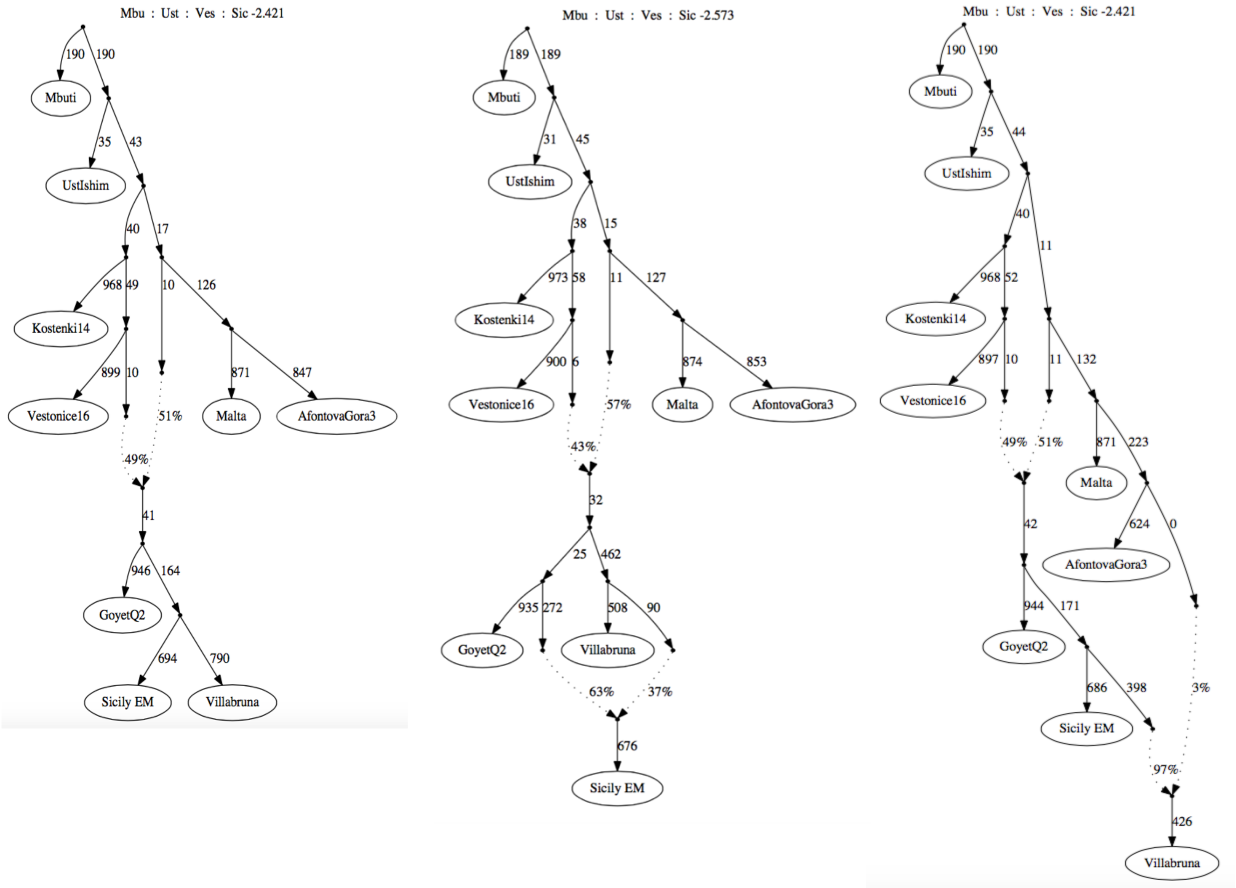


**fig. S5.3**. **EM-9 models fitting Sicily EM HGs and *Villabruna*.** Magdalenian-associated ancestry is represented by *GoyetQ2*, and ANE ancestry by *Mal'ta* and *AfontovaGora3.* Left: Sicily EM HGs and *Villabruna* form a clade with *GoyetQ2* as an outgroup to both of them (1 outlier, |$f_4$, expected – $f_4$, observed| = 2.421). Center: Sicily EM HGs as a mixture between branches related to *Villabruna* and *GoyetQ2* (1 outlier, max |$f_4$, expected – $f_4$, observed| = 2.573). Right: *Villabruna* as a mixture between Sicily EM HGs and a lineage related to *AfontovaGora3* (1 outlier, max |$f_4$, expected – $f_4$, observed| = 2.421).

Changing the proxy for Magdalenian-related ancestry to *GoyetQ116* or *El Miron* does neither result in a consistent tree topology. Using *GoyetQ116* results in Sicily EM HGs and *Villabruna* being fitted as being cladal on an admixed branch that derives 96% ancestry from a lineage that related to *GoyetQ116* and 4% from *Vestonice16* (fig. S5.4 - Left). An alternative model fits Sicily EM HGs on a branch without admixture, and *Villabruna* on an admixed branch between 97% Sicily EM HG-related and 3% *AfontovaGora3*-related ancestry (fig. S5.4 - Center). Modeling the Magdalenian-associated ancestry with *El Miron* fits the ancestry in *Villabruna* as a mixture between 91% Sicily EM HG-related and 9% *El Miron*-related ancestry (fig. S5.4 - Right).

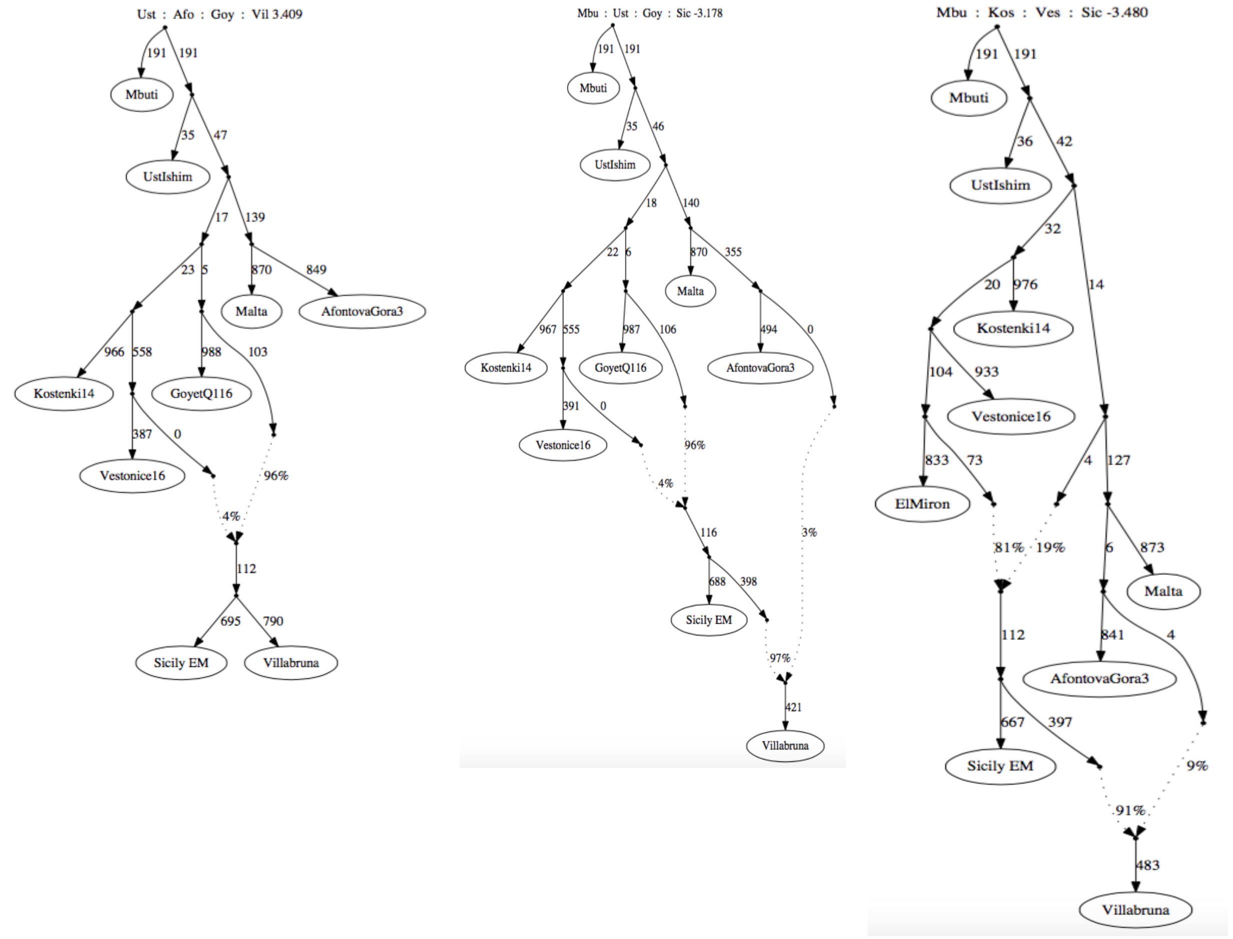


**fig. S5.4. EM-9 models fitting nine populations, including Sicily EM HGs and *Villabruna,* with Magdalenian-associated ancestry represented by *GoyetQ116* (Left, Center) or *El Miron* (Right).** Left: Sicily EM HGs and *Villabruna* form a clade on branch that derives 96% ancestry from a source close to *GoyetQ116* and 4% from *Vestonice16* (2 outliers, max |$f_4$, expected – $f_4$, observed| = 3.409). Center: *Villabruna* as a mixture between branches related to Sicily EM HGs and *GoyetQ116* (1 outlier, max |$f_4$, expected – $f_4$, observed| = 3.178). Right: *Villabruna* as a mixture between branches related to Sicily EM HGs and *El Miron*, respectively (4 outliers, max |$f_4$, expected – $f_4$, observed| = 3.480).

In a second experimental model series we additionally included a group of early Mesolithic continental HGs (EM WHGs) dating to ~13.5-11.5 kyBP, which are approximately contemporaneous to *Villabruna* and Sicily EM HGs and part of the Villabruna genetic cluster (*17, 32*). To the seven populations core model we hence added *Villabruna*, followed by Early Mesolithic continental HGs (EM WHGs) (EM-9c), and then Sicily EM HGs (EM-10), either as a branch without or with admixture. Since *GoyetQ2* as a proxy for Magdalenian- associated ancestry has sofar resulted in the smallest discrepancy between the observed and fitted allele frequencies, we proceeded with this model (fig. S5.3 - Left).

EM-9c: CM-7[*GoyetQ2*] + *Villabruna* + EM WHGs
EM-10: CM-7[*GoyetQ2*] + *Villabruna* + EM WHGs + Sicily EM HGs

When EM WHGs is added to the graph with *Villabruna (*EM-9c), the EM WHGs are fitted on a branch to which *Villabruna* contributes 95% ancestry and *GoyetQ2* contributed 5% (fig. S5.5 - Left). A model that fits *Villabruna* and Sicily EM HGs as a clade on a branch without admixture (similar to fig. S5.3 - Left) results in more significant outlier statistics (3 outliers, max $|f_4$, expected $- f_4$, observed$|$ = 4.388), and hence is less likely to reflect the true tree topology.

Subsequently, when Sicily EM HGs is added (EM-10) it is placed on a branch without admixture that falls basal to both *Villabruna* and EM WHGs, with *GoyetQ2* as the immediate outgroup (fig. S5.5 – Right). Notably, the ancestry contribution from *GoyetQ2* to EM WHGs increases to 14%, and the ancestry contribution from *AfontovaGora3* to *Villabruna* increases to 9%. We can therefore not rule out that Sicily EM HGs descend from a more basal lineage that admixed into Iberian HGs and Villabruna cluster individuals.

All in all, in our models there is a complex interaction between distal affinities to ANE- and Magdalenian-related ancestries in *Villabruna* and other Early Mesolithic HGs from continental Europe, and Sicily EM HGs. Depending on the populations included in the scaffold graph we obtained different ancestry contributions and different tree topologies. We hence could not accurately resolve the phylogeny of the Sicily EM HGs. However, eventhough *Villabruna*, Sicily EM HGs and EM WHGs share a large proportion of their ancestry, our results hint at population substructure among these HGs. *Villabruna* shows a stronger affinity to ANE ancestry, whereas Sicily EM HGs and EM WHGs show a stronger affinity to Magdalenian-related ancestry.

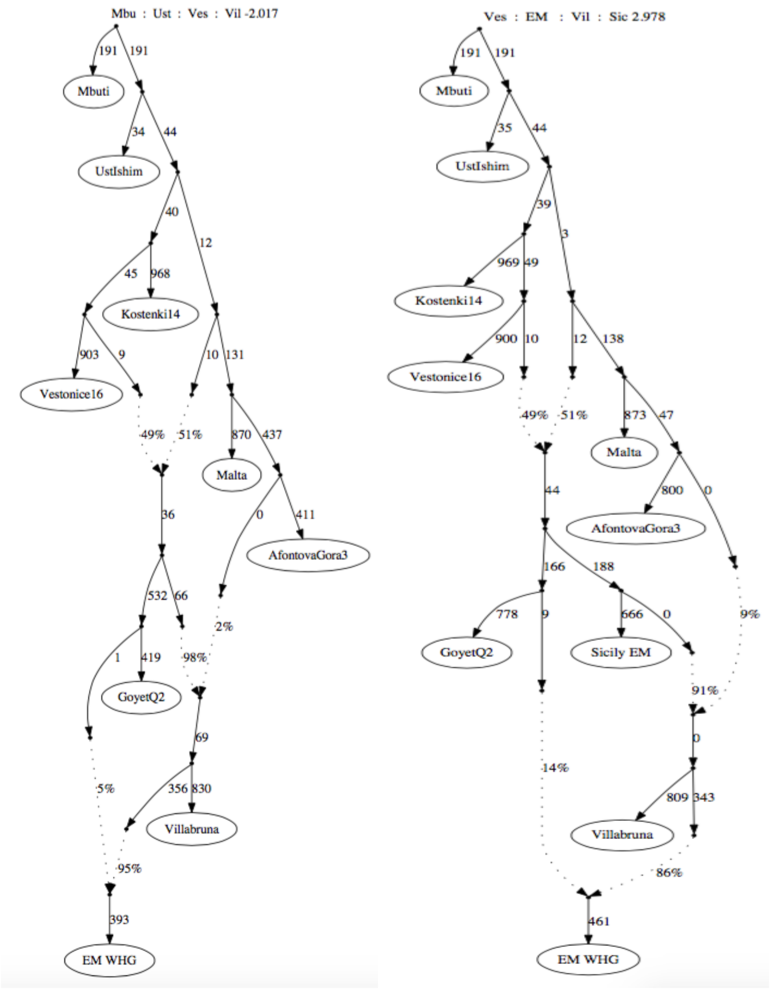


**fig. S5.5. Scaffold graphs to investigate the genetic relation of *Villabruna*, EM WHGs and Sicily EM HGs.**
Magdalenian-associated ancestry is represented by *GoyetQ2*. Left: EM-9c scaffold graph fitting 9 populations, including *Villabruna* and EM WHGs. EM WHGs is fitted on an admixed branch that derives 95% ancestry from a source close to *Villabruna* and 5% from *GoyetQ2* (1 outlier, max $|f_4$, expected $- f_4$, observed$|$ = 2.017). Right: EM-10 scaffold graph fitting 10 populations to investigate the phylogenetic position of Sicily EM HGs relative to *Villabruna* and EM WHGs. Sicily EM HGs is fitted on a branch that falls basal to both *Villabruna* and EM WHGs. Sicily EM HGs contributed ancestry to *Villabruna*, and *Villabruna* contributed ancestry to EM WHGs (1 outlier, max $|f_4$, expected $- f_4$, observed$|$ = 2.978).

### S6. Characterizing the Sicilian early farmer ancestry using *F*-statistics

First, we tested with an admixture $f_4$-statistic whether the Early Neolithic Sicilians contain a HG ancestry component in addition to their shared ancestry with early farmers from Anatolia Barcin of Greece Peloponnese. Accordingly, we performed *$f_4$(Chimp, X; Greece EN Peloponnese/Anatolia EN Barcin, Sicily EN),* where *X* are various West Eurasian HG lineages (fig. S6.1). By using Greece EN Peloponnese or Anatolia EN Barcin as a baseline for the early farmer ancestry in Sicily EN (*38*), we downweighted any shared ancestry related to this that is abundant in many Mesolithic HG lineages from southern Europe (e.g. Iron Gates HGs (*17*), and see fig. S6.2). We found that Sicily EN shows significant admixture signals for various HGs from Europe, including preceding local Sicily EM ($z_{max}$ = 4.12) and Sicily LM HGs ($z_{max}$ = 4.09) (fig. S6.1). Notably, alongside the Sicilian HGs, Mesolithic HGs from France (e.g. *Chaudardes*: $z_{max}$ = 2.89, *Ranchot88:* $z_{max}$ = 3.64), Croatia ($z_{max}$ = 2.61), and Iberia (*LaBrana*: $z_{max}$ = 2.96) are among the strongest signals for HG admixture in Sicily EN (fig. S6.1).

To test whether the Sicilian early farmers are closer to the Mesolithic HGs from France than to the preceding Sicily LM HGs, we performed $f_4$-cladality statistics of the form *$f_4$(Chimp, Sicily EN; Sicily LM HGs, X).* For this test we found no significantly positive $f_4$-values for the Mesolithic HGs from France (fig. S6.2). However, given the low number of ABBA and BABA trees, the similar ancestries in Sicily LM HGs and the Villabruna-cluster individuals from France may be driving the non-significance for this test. Notably, Mesolithic HGs from southeastern Europe, including *Croatia Mesolithic HG*, Koros EN HG and various Iron Gates HG groups, do share significantly more alleles with Sicily EN than the local Sicily LM HGs do. However, due to the pre-Neolithic gene flow between southeastern Europe and the Near East, the positive signals in this test for the shared genetic drift between Sicily EN and southeastern European HGs might either reflect an ongoing direct gene flow between these regions, e.g via maritime contact (*5*), or an indirect signal from the farmer ancestry that was brought in.

Subsequently, we aimed to find the closest proxy for the non-HG ancestry component in Sicily using *$f_4$(Chimp, X; Sicily LM HGs, Sicily EN),* where *X* are various Early Neolithic groups from West Eurasia (fig. S6.3). We here took the Sicily LM HG ancestry as a baseline for any HG ancestry in Sicily EN, and any HG ancestry broadly similar to it in *X*, if present. Hence, any Sicily LM-like HG ancestry is downweighted in this test and does not contribute to the $f_4$-statistic. We expected all Neolithic groups to result in a positive value for this test. However, the test group with the highest relative $f_4$-value was assumed to be the best proxy for the farmer ancestry component in Sicily EN. Again, we found the highest levels of shared genetic drift for various Early Neolithic farmers from the Balkan, followed by Hungary Koros and Anatolia Barcin (fig. S6.3).

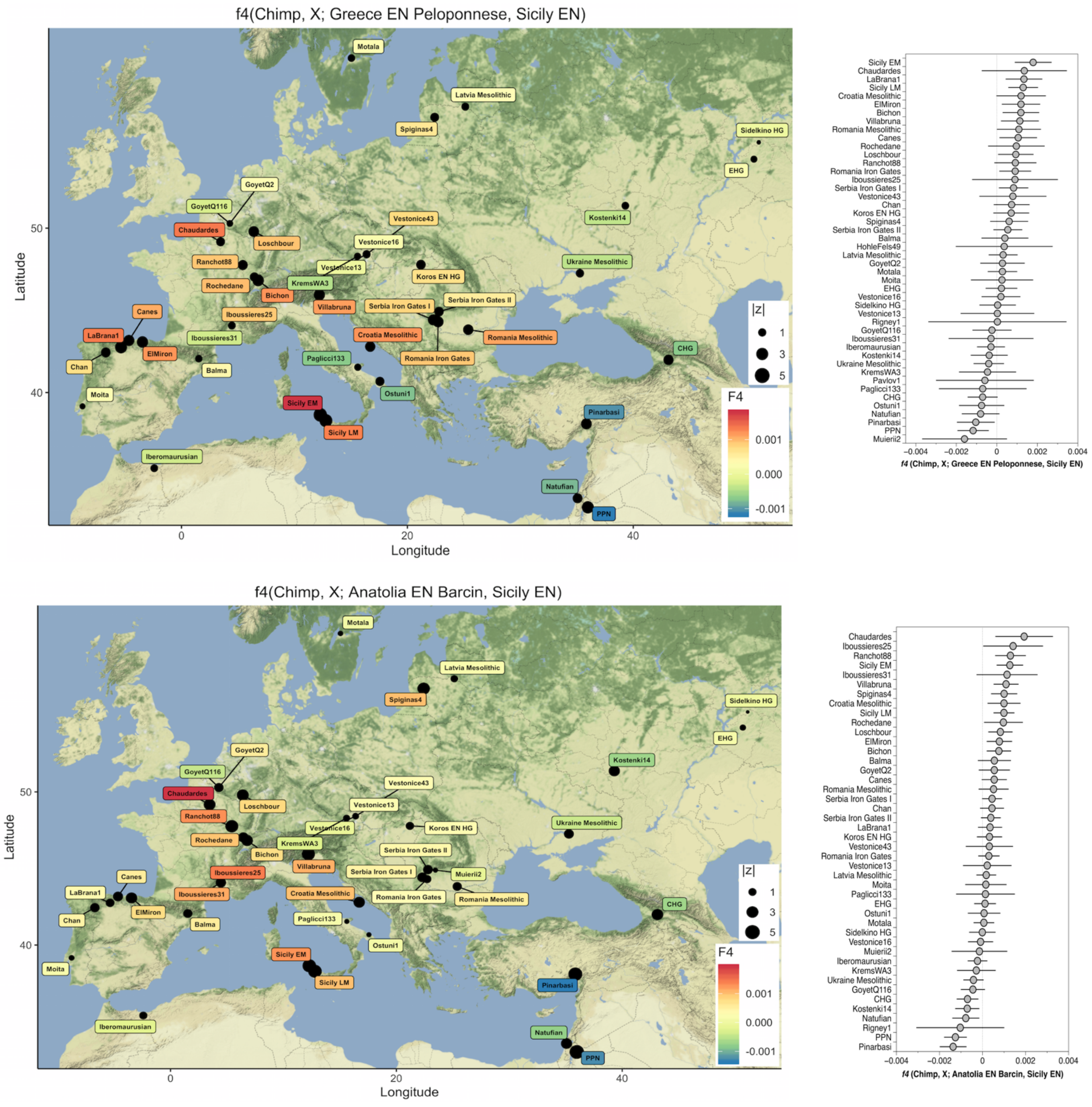


**fig. S6.1.** ***F$_4$*-admixture statistics for various HG ancestry sources in Sicily EN, relative to Greece EN Peloponnese (top) and Anatolia EN (bottom), of the form *f$_4$(Chimp, X; Greece EN Peloponnese/Anatolia EN Barcin, Sicily EN).*** Warmer colours reflect stronger signals of admixture. Dot sizes reflect |z|-scores and error bars 2 SEs.

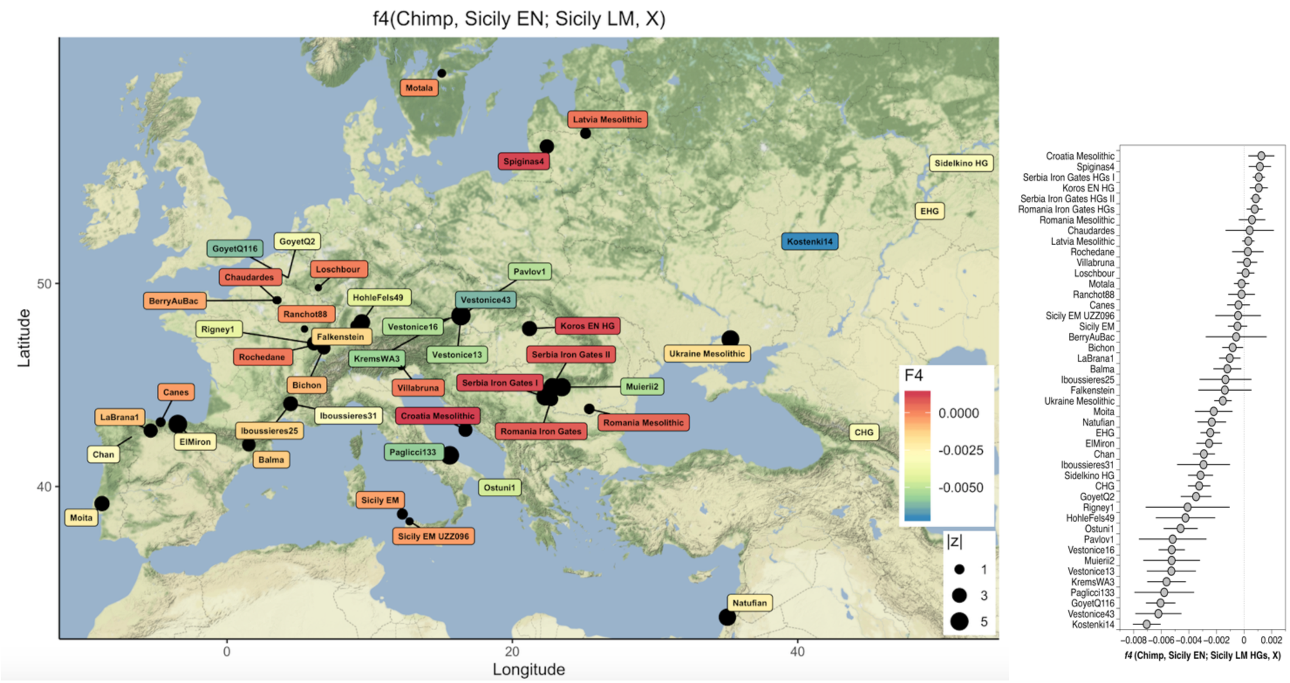


**fig. S6.2. *F4*-symmetry statistics of the form *f4(Chimp, Sicily EN; Sicily LM HGs, X)* to compare the ancestry in Sicily EN farmers to that in Sicily LM HGs and various West-Eurasian HGs (*X*).** There is no indication that Mesolithic HGs from France are genetically closer to the Sicilian early farmers than Sicily LM HGs are. Dot sizes reflect |z|-scores and error bars 2 SEs.

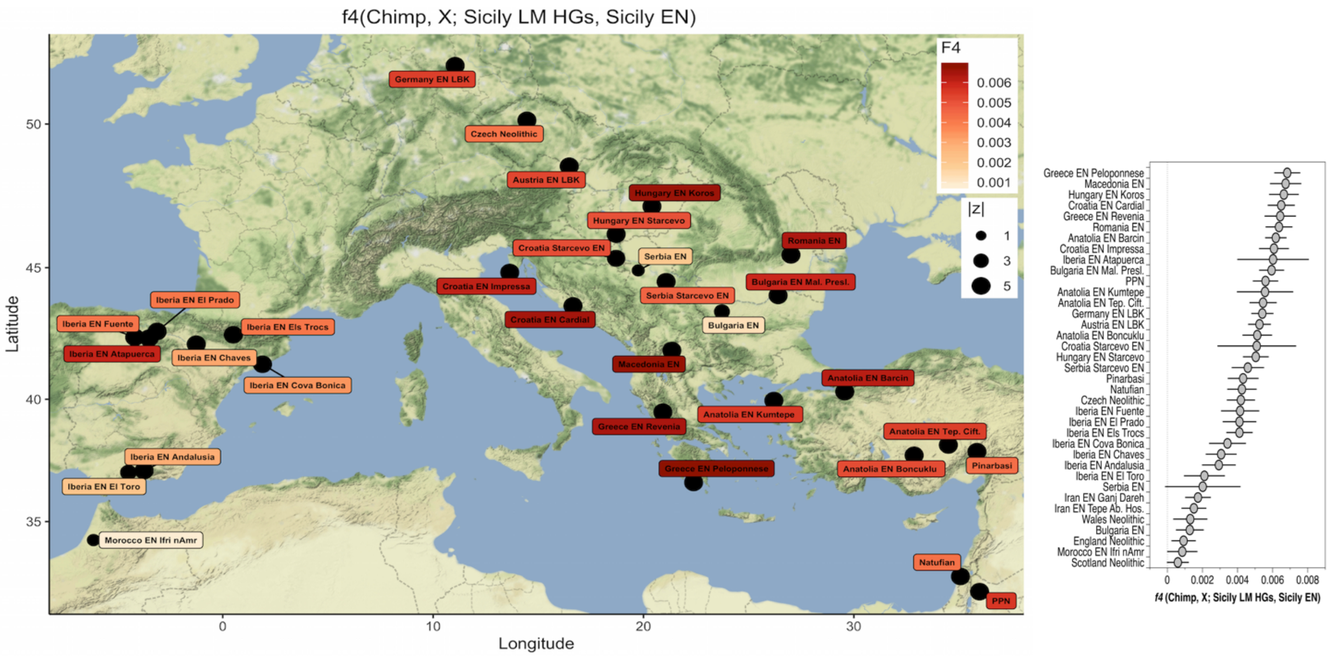


**Fig. S6.3. *F4*-admixture statistic of the form *f4(Chimp, X; Sicily LM HGs, Sicily EN)* to find the closest proxy to the early farmer ancestry in Sicily, using Sicily LM HG as a baseline for the HG ancestry.** Redder colours reflect higher levels of shared genetic drift between Sicily EN and the tested early farmer group X. Dot sizes reflect |z|-scores and error bars 2 SEs. Early farmers from the Balkan (Greece, Macedonia, Croatia) and Hungary Koros share the highest excess of alleles with Sicily EN.

Alternatively, we aimed to find the closest proxy for the non-HG ancestry in Sicily EN by measuring levels of shared genetic drift to various Early Neolithic European groups *X*, using Greece EN Peloponnese as a baseline, with $f_4$*(Chimp, Sicily EN; Greece EN Peloponnese, X)* (fig. S6.4). This test is similar to the $f_3$-outgroup statistic $f_3$*(Mbuti, Sicily EN, X)* (Fig. 4B), except that it downweights the ancestry that is shared between Sicily EN and Greece EN Peloponnese, and test group *X* and Greece EN Peloponnese. This has the advantage that any HG ancestry that is part of the gene pool shared between Sicily EN, Greece EN Peloponnese and *X* does not contribute to the *f*-statistic. In this way we can separate the population genetic affinities from distant HG admixture (e.g. from southeastern European HGs or Near Eastern HG groups), which may have resulted in population genetic substructure within the European Early Neolithic founder groups, from admixture from local HG groups *en route* as the Early Neolithic farmers expanded. Similar to the $f_3$-outgroup statistic results, we find various Early Neolithic farmers from the Balkan and Central Europe to be genetically most similar to Sicily EN (fig. S6.4).

Lastly, we performed the cladality test $f_4$*(Chimp, X; Greece EN Peloponnese, Sicily EN),* where *X* are various Early Neolithic groups from West Eurasia (fig. S6.5). By taking an immediate genetic outgroup to the Early Neolithic groups in Europe, Greece EN Peloponnese, any *en route* and locally admixed HG ancestry that is part of gene pool of Sicilian EN and is shared with test group *X* will contribute to a positive $f_4$-statistic. The test group with the most positive $f_4$-statistic hence will have a gene pool that is most similar in allele frequencies and variances compared to that of Sicily EN. We find that none of the Early Neolithic group from Europe shares significantly more alleles with Sicily EN than with Greece EN Peloponnese (fig. S6.5). Sicily EN and Greece EN Peloponnese either form a clade to the exclusion of other European Early Neolithic farmers, or the latter are genetically closer to Greece EN Peloponnese. This result could imply that the Early Neolithic Sicilians derive from an ancestral lineage that falls outside the genetic variation of other Early Neolithic groups in Europe. However, for statistics of the form $f_4$*(Chimp, Sicily EN; Greece EN Peloponnese, X)*, Sicily EN shares more alleles with various EN groups from the Balkan (Macedonia, Serbia, Croatia, Romania) and Central Europe than with Greece EN Peloponnese (fig. S6.4). Since the proportion of HG ancestry in Sicily EN exceeds that in Greece EN Peloponnese, the HG ancestry may cause outgroup attraction in the statistic $f_4$*(Chimp, X; Greece EN Peloponnese, Sicily EN*), forcing the outcomes more negative. We can, however, not exclude the possibility that the early farmer ancestry in Sicily EN falls partly outside of the broader genetic diversity of the early farmers from Europe, Greece EN Peloponnese or Anatolia Barcin. Assuming that the Late Mesolithic HGs from Sicily and the Balkan substantially overlapped both in lithic industry and genetic composition, parallel admixture events from local HGs in these regions with incoming early farmers would result in similar ancestry profiles for the early farmer groups from the Balkan and Sicily, respectively.

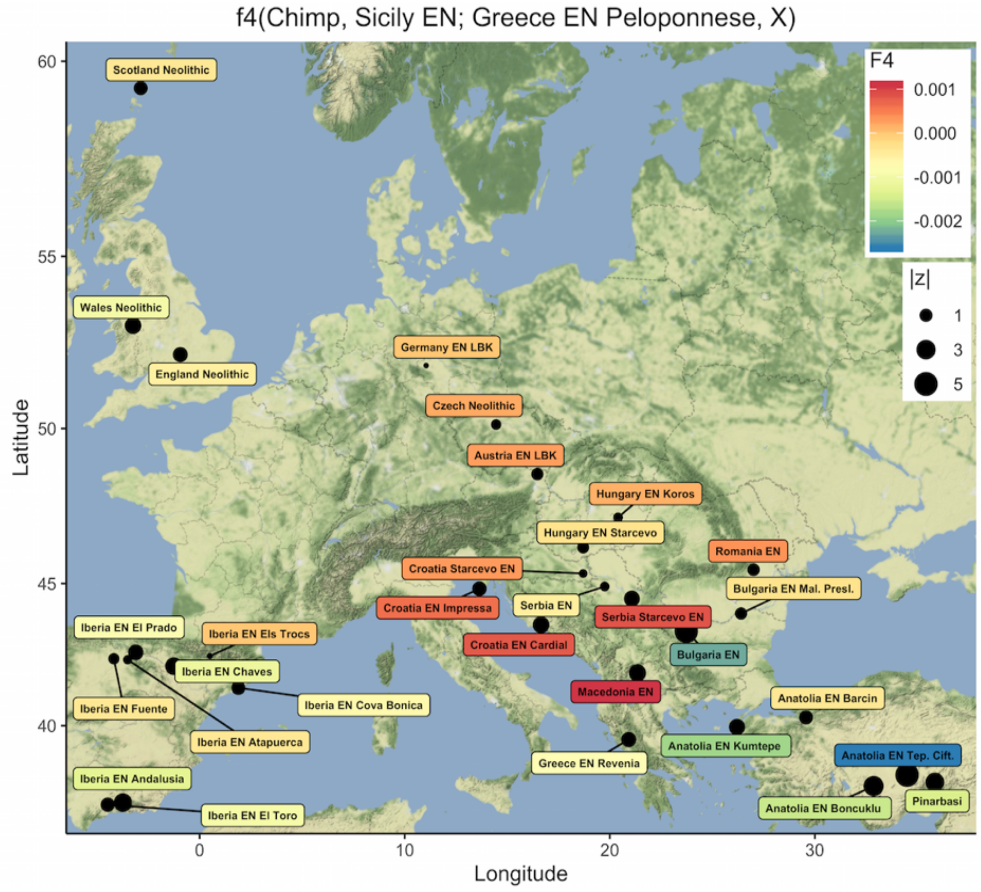


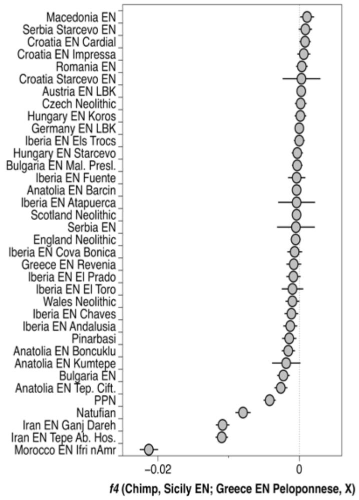


**fig. S6.4. Cladality tests for various Early Neolithic groups from West Eurasia and Greece EN Peloponnese with respect to Sicily EN, of the form: *$f_4$(Chimp, Sicily EN; Greece EN Peloponnese, X)*.** Warmer colours correspond to positive $f_4$-values and indicate that Sicily EN gene pool is closer to that of the Early Neolithic test group *X*, whereas cooler colours correspond to negative $f_4$-values and indicate Sicily EN is closer to Greece EN Peloponnese. Compared to Greece EN Peloponnese, Early farmers from the Balkan and Central Europe share a moderate excess of alleles with Sicily EN. Dot sizes reflect |z|-scores and error bars 2 SEs.

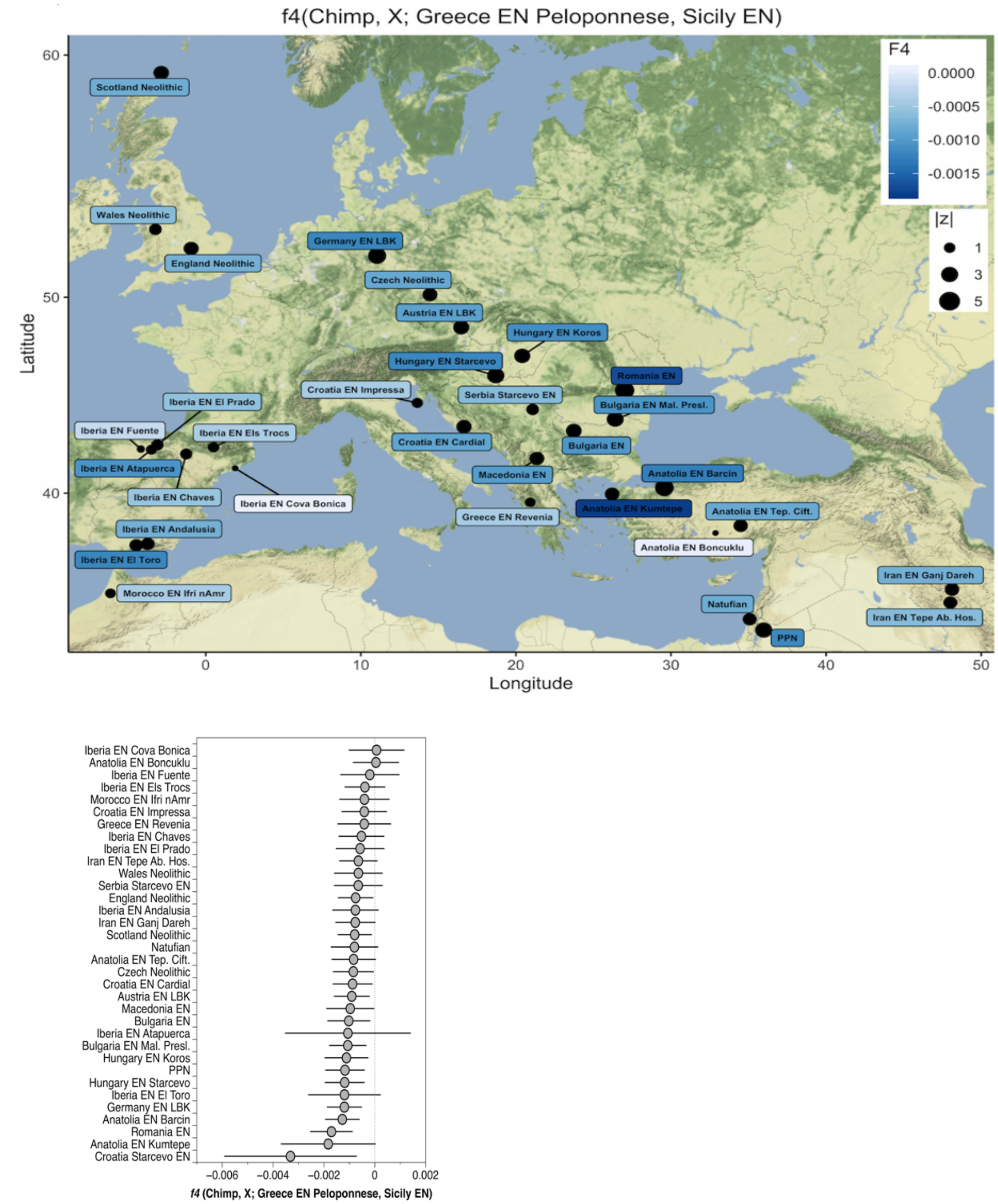


**fig. S6.5. Comparing the ancestry in Greece EN Peloponnese and Sicily EN to various early farmer groups (*X*), using an $f_4$-cladality statistic of the form *$f_4$(Chimp, X; Greece EN Peloponnese, Sicily EN)*.** All the early farmer groups are either symmetrically related to Sicily EN and Greece EN Peloponnese (white) or genetically closer to the latter (dark blue). Dot sizes reflect |z|-scores and error bars 2 SEs.

### S7: Uniparental marker haplotyping

#### A. Mitogenome haplotypes

We could reconstruct the mitochondrial genomes for 17 individuals (table S7.1, 98-100% genome coverage, mean base coverage 7 - 1,034X).

*Sicily EM HGs*

The two oldest HGs in our dataset, *UZZ5054*, *UZZ96* carried mitogenome lineages that fall within the U2'3'4'7'8’9 branch, and show a high similarity to the U2'3'4'7'8’9 haplotype that was previously reported for an Epigravettian HG from OrienteC (*I2158 - OrienteC*, (*15, 17*) (table S7.2). The three HGs have nine lineage-specific mutations in common and differently relate to each other with regard to three additional private mutations (table S7.2). U2'3'4'7'8’9 mitogenome lineages have been reported for Upper Palaeolithic European HGs associated with the Gravetian in Italy (*Paglicci108)*, Magdalenian in France (*Rigney*) and Azilian in Spain (*Balma Guilanyà*) (*50, 52*).

| Individual ID | Site | $^{14}$C age (calBCE) | Study | Genetic group label | Genetic sex | Mitogenome haplogroup | % reference covered | Mean base coverage (X) | Sd. base coverage (X) |
|---|---|---|---|---|---|---|---|---|---|
| *I2158* | OrienteC | 12,836-7,923 (strat.) | (*15, 17*) | Sicily EM | F | U2'3'4'7'8'9 | 55.6 | 2.5 | 2.0 |
| *UZZ5054* | Uzzo | 8,790-8,635 | this study | Sicily EM | F | U2'3'4'7'8'9 | 100 | 401.4 | 122.1 |
| *UZZ96* | Uzzo | 9,150-6,550 (strat.) | this study | Sicily EM | F | U2'3'4'7'8'9 | 98.4 | 6.6 | 3.4 |
| *UZZ69* | Uzzo | 6,753-6,609 | this study | Sicily LM | F | U5b2b | 100 | 17.8 | 6.0 |
| *UZZ4446* | Uzzo | 6,599-6,477 | this study | Sicily LM | F | U5b2b | 100 | 326 | 95.1 |
| *UZZ71* | Uzzo | n.a | this study | Sicily LM | F | U5a2+16294 | 100 | 259 | 75.9 |
| *UZZ79* | Uzzo | 6,684-6,596 | this study | Sicily LM | F | U5b3d | 100 | 96.4 | 24.7 |
| *UZZ88* | Uzzo | 5,989-5,850 | this study | Sicily LM | F | U5b3d | 100 | 106.6 | 31.2 |
| *UZZ81* | Uzzo | 6,682-6,595 | this study | Sicily LM | M | U5b3d | 100 | 39.4 | 12.3 |
| *UZZ80* | Uzzo | 6,683-6,596 | this study | Sicily LM | F | U5b2b1a | 100 | 344.2 | 86.8 |
| *UZZ82* | Uzzo | 6,628-6,481 | this study | Sicily LM | F | U5a1 | 100 | 889.5 | 228.5 |
| *UZZ40* | Uzzo | 6,416-6,251 | this study | Sicily LM | M | U4a2f | 100 | 1034.3 | 248.8 |
| *UZZ61* | Uzzo | n.a | this study | Sicily EN | M | K1a2 | 100 | 27.1 | 8.2 |
| *UZZ77* | Uzzo | n.a | this study | Sicily EN | F | H | 100 | 493.8 | 134.4 |
| *UZZ33* | Uzzo | 5,570-5,180 (strat.) | this study | Sicily EN | M | U8b1b1 | 100 | 521.5 | 139.7 |
| *UZZ34* | Uzzo | 5,461-5,231 | this study | Sicily EN | F | U8b1b1 | 100 | 247.2 | 87.9 |
| *UZZ74* | Uzzo | 5,326-5,220 | this study | Sicily EN | F | N1a1a1 | 99.6 | 32.3 | 12.6 |
| *UZZ75* | Uzzo | 5,327-5,220 | this study | Sicily EN | F | J1c5 | 100 | 221.6 | 65.2 |

**table S7.1. Details on the reconstructed mitogenomes and the assigned haplogroups.**

| Individual ID | MT haplogroup | Variants for called MT haplogroup (against rCRS) | Private mutations |
|---|---|---|---|
| *UZZ5054* | U2'3'4'7'8'9 | 73G 263G 750G 1438G A1811G 2706G 4769G 7028T 8860G 11467G 11719A 12308G 12372A 14766T 15326G | 1406C 5999C 6152C 6498A 7403G 9991G 10020C 14152G 15466A 16274A 16297C |
| *UZZ96* | U2'3'4'7'8'9 | 73G 263G 750G 1438G A1811G 2706G 4769G 7028T 8860G 11467G 11719A 12308G 12372A 14766T 15326G | 895T 5999C 6152C 6498A 7403G 10020C 14152G 15466A 16274A 16297C |
| *I2158/ OrienteC* | U2'3'4'7'8'9 | 750G 11719A 12308G 12372A 14766T 15326G | 14152G 15466A 16274A 16297C |

**table S7.2**. **Details on the mitogenome haplotypes for the Sicilian EM HGs.**

| Private mutations | *OrienteC (I2158)* | *UZZ5054* | *UZZ96* |
|---|---|---|---|
| 895T | no coverage | absent! 517X | 9X |
| 1406C | absent! 1X* | 321X | absent! 6X |
| 5999C | no coverage | 381X | 12X |
| 6152C | no coverage | 321X | 7X |
| 6498A | no coverage | 435X | 9X |
| 7403G | no coverage | 351X | 1X* |
| 9991G | no coverage | 208X | absent! 3X |
| 10020C | no coverage | 150X | 1X* |
| 14152G | 2X | 357X | 2X |
| 15466A | 2X | 241X | 1X* |
| 16274A | 2X | 156X | 3X |
| 16297C | 1X* | 109X | 2X |

**table S7.3. Private mutations with their coverage for the U2'3'4'7'8'9 mitogenome sequences in the Sicily EM HGs.** When a private mutation is absent, the coverage for the reference allele is given.

*Sicily LM HGs*

We found that all the individuals in the Sicily LM genetic group carried U4a, U5a, and U5b mitogenome haplogroup lineages. All of these are characteristic for West Eurasian Mesolithic HGs (*52, 128*).

Two Castelnovian-associated HGs carried haplogroup U5b2b and one a more derived variant U5b2b1a (table S7.4). The individuals who harboured U5b2b (*UZZ69* and *UZZ4446*) shared five private mutations (5585A, 9833C, 12477C, 16311C, 16355T). None of these mutations are typically found on a more derived branch, including U5b2b1a. U5b2b haplotypes were frequently observed among Villabruna cluster individuals high in WHG ancestry (*52*). The oldest individuals found so far to have carried U5b2b are two Italian Epigravettian individuals from *Grotta Paglicci* and Villabruna, and two Epipalaeolithic HGs from *Rochedane* and *Aven des Iboussières* in France (*52*). The haplogroup was also found in low frequency among Mesolithic HGs from southeastern Europe such as Croatia and Iron Gates fishermen from Serbia (~7,300-6,000 calBCE (*17*).

We also found haplogroup U5b3/U5b3d in two Castelnovian-associated HGs and in one individual tentatively contemporanous to early Impressa Ware (table S7.4). Notably, these individuals carried only one of the three expected variants that define U5b3d, and had three additional mutations in common (11836G, 16278T, 16385G). The two Castelnovian HGs, a genetic male (*UZZ79*) and female (*UZZ81*) also show a pairwise mismatch rate (PMMR) for autosomal SNP sites that is half of that found for unrelated individuals from this time period (see Extended Data Table-4). This underlines a first-degree genetic relatedness for these two individuals via at least the maternal side. Interestingly, the U5b3/U5b3d haplogroup has not been reported in European Mesolithic HGs thus far. However, Pala et al. (*129*) suggested an origin for U5b3 in the Italian Peninsula based on their analysis on the mitochondrial DNA variation observed among modern individuals. Notably, U5b3 has been found in an early Cardial farmer from the *El Portalon* cave at Sierra de Atapuerca in Spain, with a high amount of local HG ancestry (*77*). Additional sampling of Sicily Mesolithic HGs should indicate whether this haplogroup can be viewed as a general maternal lineage for the Mesolithic population in Sicily, or whether the individuals sampled here are genetic isolates.

In addition, we found U5a haplogroups in one Castelnovian-associated HG (*UZZ82*) and one individual tentatively contemporaneous to Impressa Ware (*UZZ71*) (table S7.4). *UZZ82* carried U5a1 with three additional private mutations (1007C, 3865G, 9380A). The U5a1 haplogroup has been reported for Mesolithic HGs from Russia and northern Europe (*39, 130*). *UZZ71* harboured U5a2+16294, a basal lineage to U5a2a. The more basal U5a2 haplogroup has been found in two Mesolithic hunter-gatherers from *Los Closeaux* and *Les Vignolles* in France (*52, 130*). The more derived haplogroup U5a2a is found in relatively higher frequency among Mesolithic HGs in general, more specifically in those from Ukraine, Serbia and Romania (*17*).

Lastly, for one Castelnovian-associated HG, *UZZ40* we found the rare haplogroup U4a2f without one of the four expected variants (G15172A is missing, table S7.3). Intriguingly, haplogroup U4a2f has been found also in a Cardial Ware individual from *Cueva de Chaves*, Iberia (*131*). U4a haplogroups are mostly found among Mesolithic HGs from northern Europe, the Baltic and Russia (*17, 41, 130*).

| Individual ID | MT haplogroup | Variants for called MT haplogroup (against rCRS) | Private mutations | Missing mutations |
|---|---|---|---|---|
| **UZZ79** | U5b3d | 73G 150T 263G 750G 1438G 2706G 3197C 4769G 7028T 7226A 7768G 8860G 9477A 11467G 11719A 12308G 12372A 13617C 14182C 14766T 15326G 16192T 16270T 16304C 16311C! | 11836G 16278T 16385G | 13830C 16067T |
| **UZZ81** | U5b3d | 73G 150T 263G 750G 1438G 2706G 3197C 4769G 7028T 7226A 7768G 8860G 9477A 11467G 11719A 12308G 12372A 13617C 14182C 14766T 15326G 16192T 16270T 16304C 16311C! | 11836G 16278T 16385G | 13830C 16067T |
| **UZZ88** | U5b3d | 73G 150T 263G 750G 1438G 2706G 3197C 4769G 7028T 7226A 7768G 8860G 9477A 11467G 11719A 12308G 12372A 13617C 14182C 14766T 15326G 16192T 16270T 16304C 16311C! | 11836G 16278T 16385G | 13830C 16067T |
| **UZZ80** | U5b2b1a | 73G 150T 263G 750G 1438G 1721T 2706G 3197C A3861G 4769G 7028T 7768G 8860G 9477A 11467G 11653G 11719A 12308G 12372A 13617C 12634G 13630G 13637G 14182C 14766T 15326G 15497A 16192C! 16270T 16362C | | |
| **UZZ4446** | U5b2b | 73G 150T 263G 750G 1438G 1721T 2706G 3197C 4769G 7028T 7768G 8860G 9477A 11467G 11653G 11719A 12308G 12372A 13617C 12634G 13630G 13637G 14182C 14766T 15326G 16192C! 16270T | 5585A 9833C 12477C 16311C 16355T | |
| **UZZ69** | U5b2b | 73G 150T 263G 750G 1438G 1721T 2706G 3197C 4769G 7028T 7768G 8860G 9477A 11467G 11653G 11719A 12308G 12372A 13617C 12634G 13630G 13637G 14182C 14766T 15326G 16192C! 16270T | 5585A 9833C 12477C 16311C 16355T | |
| **UZZ82** | U5a1 | 73G 263G 750G 1438G 2706G 3197C 4769G 7028T 8860G 9477A 11467G 11719A 12308G 12372A 13617C 14766T 14793G 15218G 15326G 16192T 16256T 16270T A16399G | 1007C 3865G 9380A | |
| **UZZ71** | U5a2 + 16294 | 73G 263G 750G 1438G 2706G 3197C 4769G 7028T 8860G 9477A 11467G 11719A 12308G 12372A 13617C 14766T 14793G 15326G 16256T 16270T 16294T 16526A | 3523G 5460A 15297C 16192C! | |
| **UZZ40** | U4a2f | 73G 195C! 263G 310C 499A 750G 1189C 1438G 1811G 1978G 2706G 4646C 4769G 5999C 6047G 7028T 8818T 8860G 11332T 11467G 11719A 12308G 12372A 12397G 14620T 14766T 15326G 15693C 16356C | | 15172A |

**table S7.4. Details on the mitogenome haplotypes for the Sicily LM HGs.**

*Sicily EN*

The early Sicilian farmers in our transect harboured mitogenome haplogroups characteristic for early farmers: U8b1b1 (n=2), K1a2 (n=1), N1a1a1 (n=1), H (n=1), and J1c5 (n=1) (table S7.5). All these haplogroups have previously been reported in early farmers from the Balkan, and in aceramic and ceramic Neolithic individuals from *Barcin* in north-western Anatolia (*17, 39*). Subsets of these were found among early farmers from all over Europe, albeit in different combinations and frequencies in the Balkan, Central Europe and Iberia (*132*).

U8b1b1, found in two of the early Sicilian farmers, has been reported for Starcevo early farmers from Croatia (17). Haplogroup K1a2 has been reported for early farmers from Romania, Germany LBK and northern Greece (*17, 30, 38*). In addition, K1a2 and the derived K1a2a haplogroup appear frequently among early farmers from Iberia. This includes a ~5,400 calBCE Cardial individual from *Cova Bonica* and a ~5,100 calBCE Epicardial individual from *Cova de Els Trocs* in northeastern Spain, and a ~5,000 calBCE individual from *Cueva del Toro* in southern Spain associated with 'boquique' and 'almagra' technique pottery (*26, 44, 54*).

The rare haplogroup N1a occurs at a relatively high frequency in LBK early farmers from Central Europe, but is much lower in Iberia (*132-134*). The N1a1a1 haplotype that we found in one Sicilian farmer was reported in Germany EN LBK and Hungary EN Starcevo farmers, and for one individual from *Cova de Els Trocs* (*17, 38, 54*). Interestingly, the more basal haplogroup N* was found in three Early Neolithic Cardial farmers from the *Can Sadurní* Cave in Catalonia, northern Spain.

| Individual ID | MT haplogroup | Variants for called MT haplogroup (against rCRS) | Private mutations |
|---|---|---|---|
| **UZZ33** | U8b1b1 | 73G 195C! 263G 750G 1438G 1811A! 2706G 3480G 4769G 5165T 7028T 8860G 9055A 9698C 11467G 11719A 12308G 12372A 14053G 14167T 14766T 15326G 16189C! 16234T 16324C | |
| **UZZ34** | U8b1b1 | 73G 195C! 263G 750G 1438G 1811A! 2706G 3480G 4769G 5165T 7028T 8860G 9055A 9698C 11467G 11719A 12308G 12372A 14053G 14167T 14766T 15326G 16189C! 16234T 16324C | |
| **UZZ61** | K1a2 | 73G 263G 497T 750G 1189C 1438G 1811A! 2706G 3480G 4769G 7028T 8860G 9055A 9698C 10398G! 10550G 11025C 16224C 16311C! 11299C 11467G 11719A 12308G 12372A 14167T 14766T 14798C 15326G | 152C! 9604G |
| **UZZ77** | H | 263G 750G 1438G 2706A 4769G 7028C 8860G 15326G | |
| **UZZ75** | J1c5 | 73G 185A 228A 263G 295T 462T 489C 750G 1438G 2706G 3010A 4216C 4769G 5198G 7028T 8860G 10398G! 11251G 11719A 12612G 13708A 14766T 14798C 15326G 15452a 16069T 16126C | |
| **UZZ74** | N1a1a1 | 199C 204C 669C 750G 1438G 1719A 2702A 2706G 3336C 4769G 5315G 7028T 8860G 8901G 10238C 10398G! 11719A 12501A 12705T 13780G 14766T 15043A 15326G 16147A 16172C 16223T 16248T 16248T | 5460A 11884G |

**table S7.5**. **Details on the mitogenome haplotypes for the Sicilian early farmers.**

### B. Y-chromosome haplotypes

We could determine the Y-haplogroup for four males (table S7.6). Two Sicilian LM HGs associated with the Castelnovian carried haplogroups I and I2a2, which both are characteristic for Upper Palaeolithic and Mesolithic HGs from West-Eurasia. Haplogroup I is commonly found among individuals associated with the Gravettian and part of the Vestonice genetic cluster, such as *Paglicci133* from Italy and *KremsWA3* from lower Austria, and in Magdalenian-associated individuals, such as *Hohlefels49* and *Burkhardtshohle (16).* In addition, I and the more derived I2, and I2a haplogroups are the most frequent haplotypes found among European Mesolithic HGs related to the Villabruna cluster. This includes from France the Epipaleolithic *Rochedane* (haplogroup I), Mesolithic *Chaudardes* (haplogroup I) and Mesolithic *BerryAuBac* (haplogroup I), from Hungary the *Koros* individual from a Neolithic context but an ancestry profile characteristic for WHG (haplogroup I2a), and Mesolithic *Loschbour* (haplogroup I2a1b) from Luxembourg (*16, 37, 39*). Moreover, haplogroup I2 was reported for *Bichon* from Switzerland associated with the Azilian (*32*). The haplogroup I2a2 that we find in one Sicily LM HG occurred relatively frequently among Mesolithic HGs from the Iron Gates and Latvia (*17*).

The two Sicilian early farmers carried haplogroups C1a2 and H. Interestingly, although C1 haplotypes are found among early farmers, these are in general considered to be more typical for pre-

Neolithic West-Eurasians. One of the oldest individuals to have carried C1a are *GoyetQ116* and *Vestonice16* associated with the Gravettian *(16).*The more derived haplogroup C1a2 that we find in one of the Sicilian early farmers, was found in the Mesolithic *LaBrana* HG from Iberia (*42*). However, C1a2 haplotypes have also been reported for early farmers from *Barcin* and *Tepe Ciflik*, as well as in the ~13,300 calBCE Pınarbaşı HG in Anatolia (*25, 34, 39*). C1a2 haplotypes were also found in early Cardial farmers from Croatia and early LBK farmers from Austria (*17*).

The Y-haplogroup H that we find in one of the Sicilian early farmers has been proposed to be among the genetic markers of the early farmer populations of Middle East that were introduced to Europe during the Neolithic transition (*135*).

| **Individual ID** | **Genetic group label** | **Mitogenome haplogroup** | **Y-chromosome haplogroup** | **Y derived SNPs supporting haplogroup determination** |
|---|---|---|---|---|
| **UZZ40** | Sicily LM | U4a2f | I | I: L578 L755 L758 CTS48 CTS646 CTS7502 CTS8742 CTS9860 PF3640 PF3660 PF3665 PF3668 PF3796 PF3797 PF3809 PF3817 PF3822 PF3837 FGC2412 FGC2414 |
| **UZZ81** | Sicily LM | U5b3/U5b3d | I2a2 | I: V218.2 L578 L751 L755 L758 CTS4848 CTS6231 CTS6265 CTS7329 CTS7831 CTS8876 CTS9618 CTS11540 PF3640 PF3661 PF3668 PF3814 PF3837 FGC2413<br>I2a2: M436 P217 P218 L35 L37 |
| **UZZ33** | Sicily EN | U8b1b1 | H | HIJK: M578<br>H: M2713 M2896 M2936 M2945 M2992 M3035 M3058 M3062 M3070 Z4309 |
| **UZZ61** | Sicily EN | K1a2 | C1a2 | C: P255 P260 V77 V183 V199 V232<br>C1a: CTS11043<br>C1a2: Z28922 |

**table S7.6. Details on the Y-chromosome haplogroup assignments for genetic males.**
